# Measuring intramolecular connectivity in long RNA molecules using two-dimensional DNA patch-probe arrays

**DOI:** 10.1101/2023.03.12.532302

**Authors:** Timothy K. Chiang, Ofer Kimchi, Herman K. Dhaliwal, Daniel A. Villarreal, Fernando F. Vasquez, Vinothan N. Manoharan, Michael P. Brenner, Rees F. Garmann

## Abstract

We describe a simple method to infer intramolecular connections in a population of long RNA molecules in vitro. First we add DNA oligonucleotide “patches” that perturb the RNA connections, then we use a microarray containing a complete set of DNA oligonucleotide “probes” to record where perturbations occur. The pattern of perturbations reveals couplings between different regions of the RNA sequence, from which we infer connections as well as their prevalences in the population. We validate this patch-probe method using the 1,058-nucleotide RNA genome of satellite tobacco mosaic virus (STMV), which has previously been shown to have multiple long-range connections. Our results not only indicate long duplexes that agree with previous structures but also reveal the prevalence of competing connections. Together, these results suggest that globally-folded and locally-folded structures coexist in solution. We show that the prevalence of connections changes when pseudouridine, an important component of natural and synthetic RNA molecules, is substituted for uridine in STMV RNA.

## Main text

Long sequences of ribonucleotides not only can carry genetic information but can also adopt functional structures that catalyze reactions and regulate cellular pathways. RNA secondary structures span a range of scales: nucleotides nearby in the sequence can form local structures such as hairpins, and nucleotides separated by large distances along the sequence can pair to form long-range connections. These long-range connections can significantly alter the overall size and shape of an RNA molecule^1, 2^ and the accessibility of its local structures^3^. Furthermore, a population of RNA molecules can have a distribution of competing long-range connections that might reflect different biologically relevant conformations^4–7^.

Determining long-range connectivity and its variations is challenging. Direct structural techniques, such as X-ray crystallography^8^ and cryo-electron microscopy^9^, are not well suited to long and heterogeneous RNA molecules. And indirect techniques, such as chemical probing by SHAPE^10^ and DMS^11^, measure the conformational flexibility of each nucleotide in the sequence, which is related to the probability that each nucleotide is connected to another in the sequence but does not directly reveal the endpoint of the connection or if competing connections are present^12^. Thermodynamic folding models can be used to infer this missing information^13–15^, but uncertainties in the model parameters affect the accuracy of detecting long-range connections^16^.

Multidimensional probing techniques^17–21^ address some of these challenges by more directly measuring connections between nucleotides. These techniques involve perturbing an RNA molecule at specific points and then detecting the effects of the perturbations elsewhere. For example, mutate-and-map^17^ involves introducing point mutations into the sequence and then measuring corresponding changes in the conformational flexibility of the other nucleotides by chemical probing. While this technique can detect distributions of connections, the range of connections is limited by the length of sequencing reads, which is currently 250 nucleotides (nt) or so.

Proximity ligation techniques^22–27^, such as PARIS^23^, avoid read-length limitations by covalently linking connected nucleotides and then detecting the linked segments by sequencing. The protocols involve cross-link formation, fragmentation, enrichment of the linked fragments, ligation, removal of the cross-links, reverse transcription, PCR amplification, and high-throughput sequencing, followed by computational analysis of the sequencing data. While some research groups have successfully adopted these protocols, other groups may not have access to—or expertise in—the required techniques. Therefore we aimed to develop a simpler method.

Our method of determining RNA connectivity is based on DNA probing, a technique in which RNA secondary structure is inferred from how strongly RNA molecules bind to complementary DNA oligonucleotides^28–34^. In contrast to the traditional DNA probing technique, developed over 50 years ago by Uhlenbeck *et al.*^28^, we take a multidimensional approach similar to that described by Kaplinski *et al.*^35^. First we bind DNA oligonucleotide “patches” to specific regions of the RNA molecule to perturb the intramolecular connections^36^. Then we determine whether and where perturbations occur by measuring the binding of the patched RNA molecules to DNA oligonucleotide “probes” contained on a microarray^37^. The exceptional specificity of oligonucleotide hybridization enables us to perturb the RNA at many points and read out the perturbations in parallel on a single array, without covalent modification, reverse transcription, amplification, sequencing, or folding models. From the pattern of perturbations, we can infer long-range connections and their prevalences in a population of long RNA molecules.

## Results

We apply the patch-probe method outlined in **Fig. 1A-C** to the 1,058-nucleotide (nt) RNA genome of satellite tobacco mosaic virus (STMV), a long-standing model system that has been studied using chemical probing^38–41^, computational modeling^42, 43^, and direct imaging^39, 44^. We use fluorescently-labeled STMV RNA (**Methods**), a series of 44 24-nt patches (**Methods**), and a single microarray containing all 12- and 24-nt probes that are complementary to the RNA (**Methods** and **Fig. 1D**). We discuss experiments with 12-nt probes throughout the main text of this paper, and we include experiments with 24-nt probes as **Extended Data**.

**Fig. 1:**
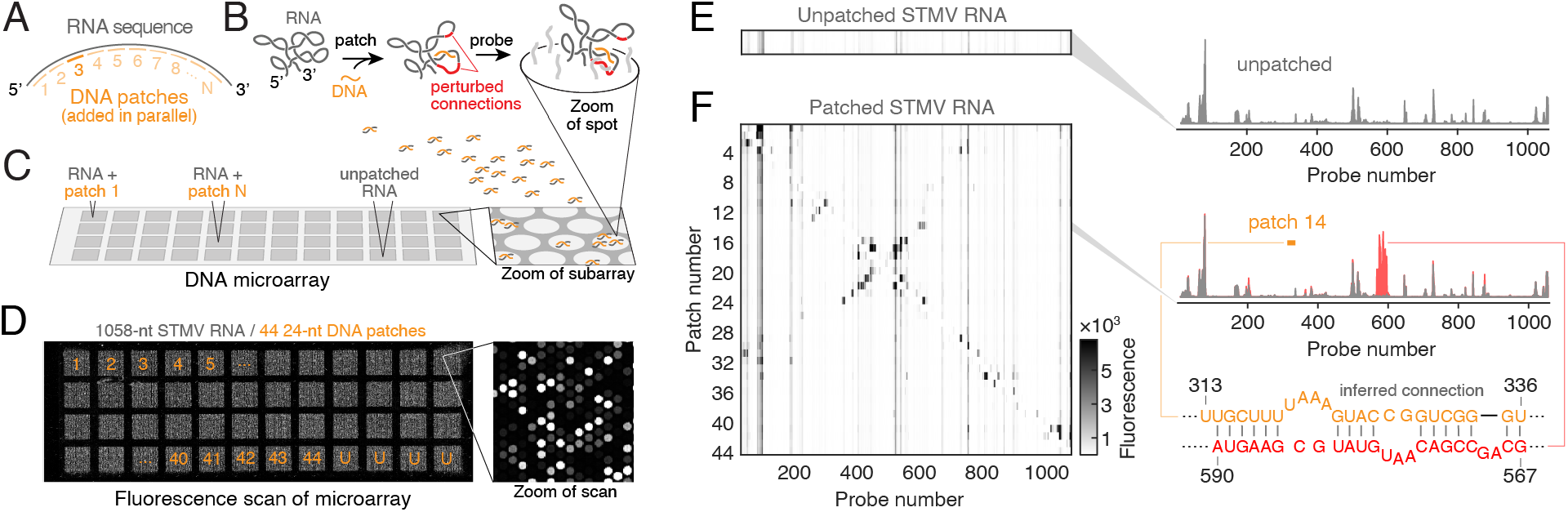
Basis of the patch-probe method. **A.** Diagram of DNA patches that are complementary to the RNA. **B.** Diagram of the patch-probe method, showing how a patch can perturb an intramolecular connection, affecting the binding of surface-tethered probes. **C.** We add each patched RNA to its own section of a DNA microarray and add unpatched RNA to a separate subarray. The microarray consists of spots, each of which contains multiple copies of a probe tethered at its 3’ end by a poly-T spacer. Within each section, every probe of a specified length that is complementary to the RNA appears in 3-12 spots. **D**. A fluorescence micrograph of a microarray containing 12- and 24-nt probes. We determine binding by labeling the RNA and measuring the fluorescence of each spot, which is proportional to the number of bound RNA molecules. Here we show the integrated fluorescence in 48 sections, 44 of which contain patched STMV RNA and 4 of which contain unpatched (“U”) STMV RNA. Each patch is 24 nt, and the 44 patches completely tile the STMV RNA sequence from nucleotide 1 to 1,056. **E**. Heatmap (left) and a bar plot (right) of the binding spectrum of unpatched RNA to 12-nt probes, as detailed in **Methods**. This 1D measurement reveals a broad range of fluorescence magnitudes for the probes. **F**. Heatmap of the binding spectrum of patched RNA to 12-nt probes (left). This 2D measurement consists of all the 1D patched-RNA spectra sorted by patch number. The vertical lines correspond to those in the unpatched RNA heatmap. Features that stand out above this background reflect patch-induced perturbations in RNA connectivity. The binding spectrum for RNA hybridized to patch 14 (right, top) is shown in red, with the spectrum of unpatched RNA overlaid in gray. The binding site of patch 14 (nucleotides 313-336) is shown in yellow. The difference between the patched and unpatched spectrum shows a large increase in binding at probes 200 nt downstream of the patch site, indicating a potential connection between these regions of the RNA sequence (right, bottom). See **Extended Data Fig. 1** for a detailed diagram of these patch and probe interactions, and **Extended Data Fig. 2** for binding spectra of 24-nt probes.

### Increases in probe binding after patching reveal intramolecular connections

Although the binding spectrum for unpatched RNA contains many peaks that, in principle, contain information about its connectivity (**Fig. 1E**), such information is in practice difficult to extract from these peaks alone. The fundamental problem is that 1D probing measurements do not directly determine which regions of the molecule are connected. Furthermore, any information about connectivity is convolved with variations in probe-binding affinity (**Extended Data Fig. 3**).

We resolve this problem by examining how the patches affect the binding, yielding a 2D measurement. The 2D spectrum has a background of vertical lines that correspond to peaks in the unpatched spectrum and therefore reflect the affinity of probes to the unpatched RNA (**Fig. 1E**). However, several patch-probe combinations show features that stand out above the background of vertical lines (**Fig. 1F**). These features correspond to an increase in probe binding, which we expect to occur if a patch disrupts a connection to a probe site in some fraction of the RNA molecules. For example, upon addition of patch 14, we observe an increase in binding for probes 560-590 (**Fig. 1F, right**). We interpret the coupling between patch 14 and probes 560-590 as evidence of a connection in the RNA involving segments that contain the patch and probe binding sites. Such couplings contain more direct information about the connections than the background itself.

While the peaks in the raw 2D binding spectrum shown in **Fig. 1F** suggest a pattern of connections in the molecule, this pattern is obscured by the background. We therefore separate changes in binding from the background. Specifically, for each patch-probe combination, we define the coupling signal 𝑆_𝑖𝑗_ to be the change in how probe number *i* binds to RNA attached to patch number *j*, relative to how probe *i* attaches to unpatched RNA: 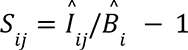, where 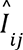 is an estimate of the true patch-probe fluorescence and 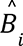 is an estimate of the true background. We use a Bayesian approach to infer the signal 𝑆_𝑖𝑗_ because the approach allows us to quantify the uncertainty on the signal using all of the measurements and to account for experimental effects such as outliers and variations in RNA amounts across subarrays. The statistical model and Markov-chain Monte Carlo sampling scheme are detailed in **SI**.

The resulting signals and uncertainties, shown in **Fig. 2**, are independent of variations in probe affinity and in RNA amounts across the microarray (see **Extended Data Figs. 4**), allowing us to directly compare coupling strengths between a given patch site and all probe sites. We do not expect the map of the signals to be perfectly symmetric, since patches and probes are different sizes, and there are variations in patch affinity. Nonetheless, the map shows a roughly symmetric pattern, with credible signals (as determined by the mean of the marginalized posterior of the signal divided by the standard deviation) occurring both near to and far from the diagonal. Signals near the diagonal, scattered from upper left to lower right, reflect short-range connections. Signals that extend far away from the diagonal, such as the long central ridge that cuts perpendicular to it, reflect long-range connections.

**Fig. 2:**
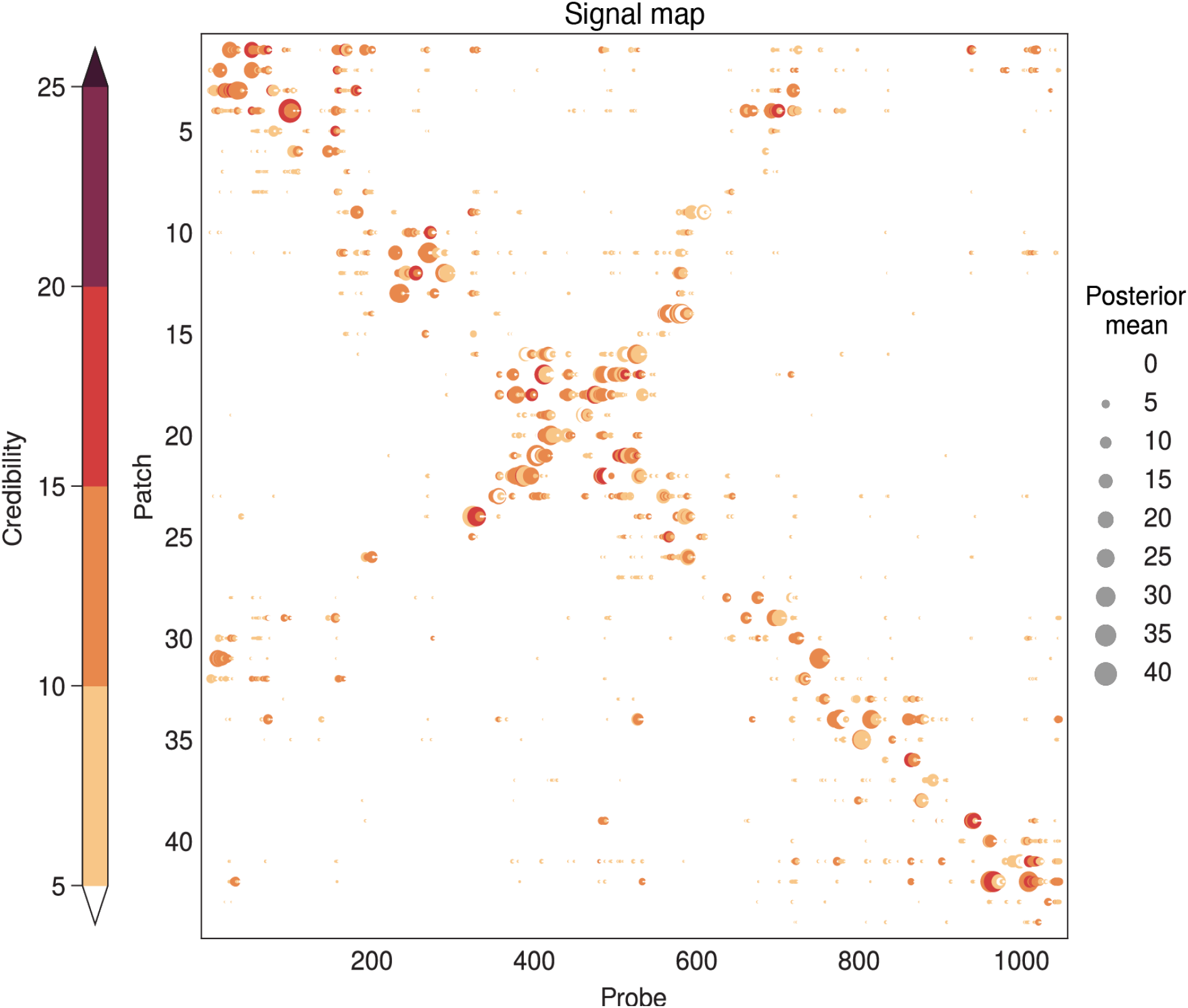
The patch-probe coupling signals suggest many possible connections in STMV RNA. The map of signals inferred from Fig. 1E, as described in **SI**. The size of each dot is proportional to the signal, and the color reflects the credibility, which we define as the ratio of the posterior mean to standard deviation. See **Extended Data Fig. 5** for the map corresponding to 24-nt probes.

### Dominant patch-probe signals reveal consensus duplexes of STMV RNA

The signal map (**Fig. 2**) yields a large amount of information about the couplings. We focus first on the dominant couplings, corresponding to the probes with the largest signals for each patch (**Methods**). The map of dominant couplings, shown in **Fig. 3A**, shows many of the features that stand out in the raw binding spectrum of **Fig. 1E**.

**Fig. 3:**
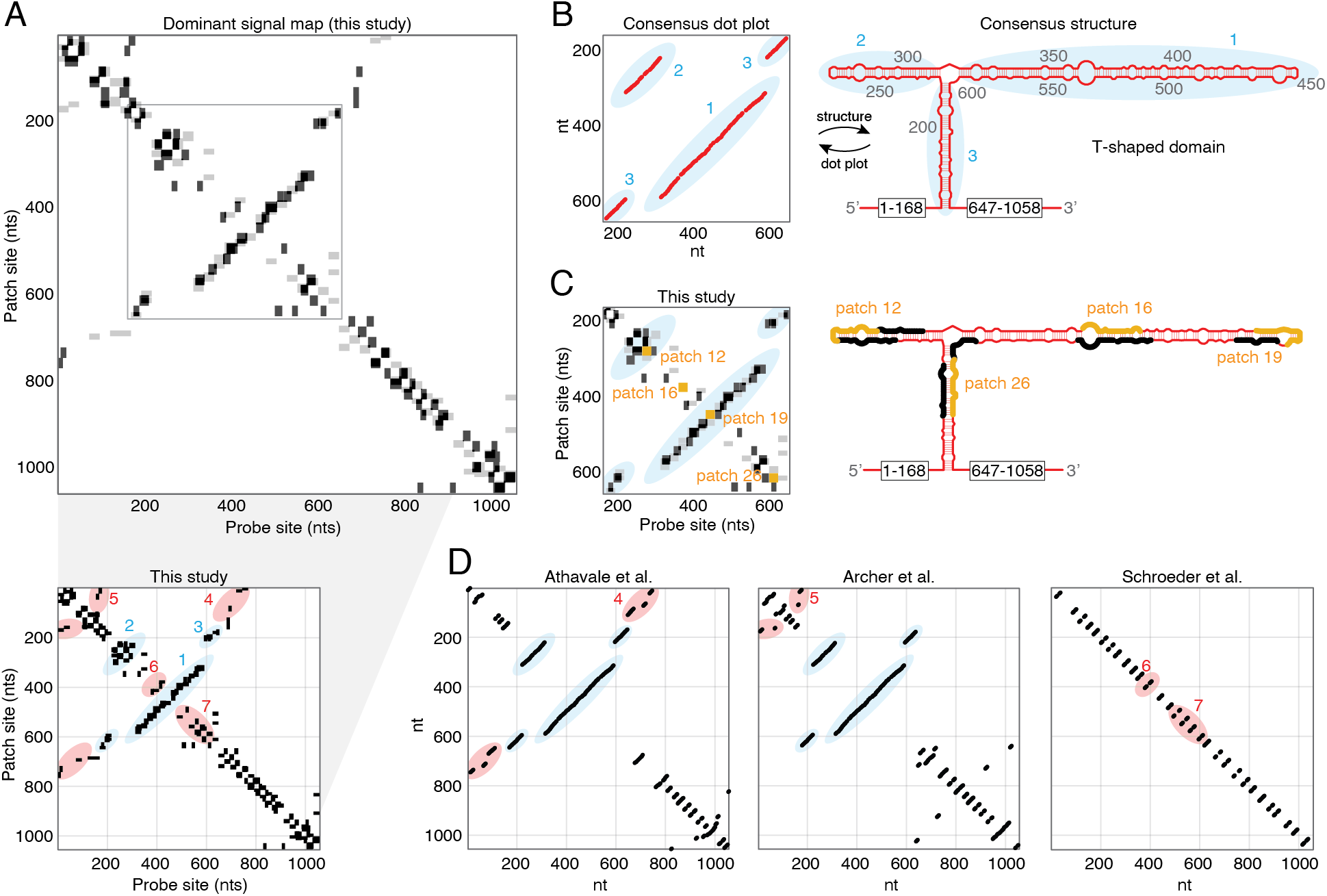
The dominant patch-probe signals are consistent with the consensus structure of STMV RNA and reveal competing structures in non-consensus regions. **A.** The map of dominant couplings (top) shows the five probes with the largest signals for each patch. Each dominant coupling is plotted as a black rectangle spanning the patch and probe binding sites. We plot the transpose of the map in gray to highlight couplings that may be obscured by larger features induced by other patches. The gray box shows the central region of the RNA, which adopts the consensus T-shaped structure. At bottom is the same plot in which ellipses highlight three long-range connections in the central region (blue ellipses 1-3), two long-range connections outside of the central region (red ellipses 4 and 5), and two stretches of short-range connections (red ellipses 6 and 7). **B.** A dot plot and a diagram of the consensus structure, with base pairs shown in red and three long-range connections highlighted by blue ellipses 1-3. **C.** Plot of the central region (gray box in panel A) of the dominant coupling map, showing consistency with three long-range connections, highlighted by blue ellipses. The binding sites of four patches are shown in yellow. A diagram of the T-shaped domain of the consensus structure is shown in red, with the binding sites of the four patches highlighted in yellow, and the largest changes in probe binding shown in black. **D.** Dot plots of the structures reported in previous chemical probing studies contain different sets of connections outside the consensus region. Some of these connections are seen in the set of dominant connections inferred from the patch-probe data: SHAPE probing of RNA transcribed *in vitro* by Athavale *et al.*^38^ shows connections in ellipse 4; SHAPE probing of RNA extracted from virus particles by Archer *et al.*^39^ shows connections in ellipse 5; DMS, kethoxal, and CMCT chemical probing and crystallographic analysis of RNA packaged in virus particles by Schroeder *et al.*^41^ shows connections in ellipse 6 and 7. See **Extended Data Fig. 6** for dominant couplings corresponding to 24-nt probes and arc plot representations of the data.

Couplings near the center of **Fig. 3A** provide a valuable point of comparison between our measurements and previous structural studies. Three different SHAPE chemical probing studies^38–40^, as well as direct measurements using atomic force microscopy^39^ and cryo-electron microscopy^44^, suggest that the central region of STMV RNA adopts a T-shaped domain containing three long-range connections: a 90-nt-long hairpin and a 270-nt-long hairpin branching from a 50-nt-long central duplex that connects regions over 470 nt apart (highlighted by ellipses in **Fig. 3B**).

Although the resolution of our map, set by the patch and probe size, is coarser than that of a dot plot, our map reveals couplings that are consistent with all three long-range connections in the consensus T-shaped domain (**Fig. 3C, left**). Visualizing these couplings on top of a structural model of the T-domain further demonstrates the good agreement between our patch-probe measurements and the consensus structure (**Fig. 3C, right**).

### Dominant patch-probe couplings detect competing connections

Having validated our results against the consensus structure, we now examine features outside the central T-shaped region. Here there is less consensus. Previous chemical probing studies by Athavale *et al*.^38^, Archer *et al*.^39^, and Schroeder *et al*.^41^ show considerable differences in connectivity (**Fig. 3D**). Some of these may be due to differences in the chemical probing protocol or the source of the RNA, and others might reflect differences in the folding models used to interpret the data. Both Athavale *et al*. and Archer *et al*. used thermodynamic folding models, but Archer *et al*. imposed a cutoff length of 600 nt for base-pair interactions. Schroeder *et al*. developed a cotranscriptional folding and assembly model with a short cutoff of 30 nt. The effects of these cutoffs are made clear by the dot plots in **Fig. 3D**. For example, the dot plot of the structure from Athavale *et al.* contains dots far from the diagonal that represent long-range connections, the longest of which connects nucleotides 12 and 746 (**Fig. 3D, ellipse 4**). These long-range connections are necessarily absent in the dot plots of the structures reported by Archer *et al.* and Schroeder *et al*. because they lie beyond the imposed cutoffs. Cutoffs can also affect the inference of shorter-range pairs by constraining the folding model. Such indirect effects could explain why mid-range base pairs in the dot plot from Archer *et al.* (**Fig. 3D, ellipse 5**) are absent in the dot plot from Athavale *et al*.

With the above differences in mind, we compare our map of dominant couplings (**Fig. 3A, bottom**) to the previously reported structures (**Fig. 3D**). Outside of the central T-domain, we observe couplings far from the diagonal that are consistent with the longest-range connections in the structure reported by Athavale *et al.* (**ellipse 4**; see, for example, features between nucleotides 1-27 and 721-768 in **Fig. 3A, bottom** and between nucleotides 12-24 and 735-746 in **Fig. 3D**). We also observe slightly off-diagonal couplings consistent with mid-range connections in the structure reported by Archer *et al.* (**ellipse 5**), and several near-diagonal couplings consistent with local connections in the structure reported by Schroeder *et al.* (**ellipses 6 and 7**). Each of these connections is found in only one of the previous structures and not the others.

Some of these connected regions cannot coexist within the same structure, suggesting that our measurements detect the presence of multiple structures within the population of RNA molecules. For example, the central T-domain shown by **ellipses 1-3** and the locally folded regions shown by **ellipses 6 and 7** cannot exist in the same molecule. Furthermore, the long-range connected region shown in **ellipse 4** cannot coexist with the shorter-range connection shown in **ellipse 5** in the same molecule. These coexisting features correspond to mutually exclusive connections with markedly different ranges, suggesting that the population of RNA molecules contains multiple structures with qualitatively different shapes and sizes.

That long RNA molecules might adopt a distribution of structures is not unexpected. Equilibrium base-pairing probabilities calculated by RNA folding algorithms also predict a distribution of structures. Although we do not claim that our system is in equilibrium, we note that predictions of folding algorithms are qualitatively consistent with our measurements. Specifically, RNAfold from the ViennaRNA software package^15^ shows that nucleotides in the central region between 400 and 600 can form both short- and long-range connections (**Extended Data Fig. 7**), consistent with dominant couplings in ellipses 1-3 and ellipses 6 and 7.

### Normalizing by the patch affinities reveals additional RNA connections and their prevalence

Our analysis thus far has focused only on the dominant couplings. The signal map (**Fig. 2**) reveals many other less strong yet credible couplings that reflect additional connections. But because the coupling signals depend on both the RNA connectivity and the patch affinity, the signal map does not distinguish between, for example, couplings that arise because a large fraction of RNA molecules are connected at the patch site and those that arise because a smaller fraction are connected, but the patch is more efficient at disrupting those connections. To separate these effects and glean information about the prevalence of connections, we must account for variations in the patch affinities.

We infer the patch affinities directly from the microarray data, and specifically from measurements of probes binding to sites that overlap completely with patch sites (these data correspond to diagonal elements of the signal map). We use a linear model and a Bayesian inference scheme, as detailed in **SI**. Briefly, we assume that the coupling signal is linearly proportional to the patch affinity—the simplest assumption we can make—and infer a normalized signal 𝑅_𝑖𝑗_ = 𝑆_𝑖𝑗_/𝑝_𝑗_, where 𝑝_𝑗_ is the patch affinity, or fraction of RNA molecules that are patched. We then rescale the normalized signal as 𝑓_𝑖𝑗_ = 𝑅_𝑖𝑗_/(1 + 𝑅_𝑖𝑗_) to produce values that lie between zero and one. Under a restrictive set of assumptions (see **SI, Supplementary Methods, section 4**), 𝑓_𝑖𝑗_ is the fraction of molecules in the population that contain a connection between probe site *i* and patch site *j*. But even when these assumptions are relaxed, we expect 𝑓_𝑖𝑗_ to scale monotonically (though not necessarily linearly) with how frequently the connection appears in the population, because the values are corrected for both patch and probe binding affinities. For this reason we call 𝑓_𝑖𝑗_ the “prevalence” of connection 𝑖𝑗. We can compare prevalences among all patch-probe combinations and even among different experiments.

The map of 𝑓_𝑖𝑗_ in **Fig. 4** reveals many features with high prevalence in the population, including some features that have not been reported previously. For example, we observe a cluster of features with high prevalence far from the diagonal, which point to long-range connections spanning nearly the entire RNA sequence (**Extended Data Fig. 8**). These very long-range connections have not been reported in previous studies, but are predicted by RNAfold^15^ to be present in the population of equilibrium structures (**Extended Data Fig. 7**). We also see isolated long-range features that are not part of a cluster and might reflect long-range pseudoknots (**Extended Data Fig. 8**). As with the dominant couplings, some features in the prevalence map point to competing connections that cannot appear in the same structure (**Extended Data Fig. 8**), providing further evidence that STMV RNA adopts multiple structures, at least under our experimental protocols.

**Fig. 4:**
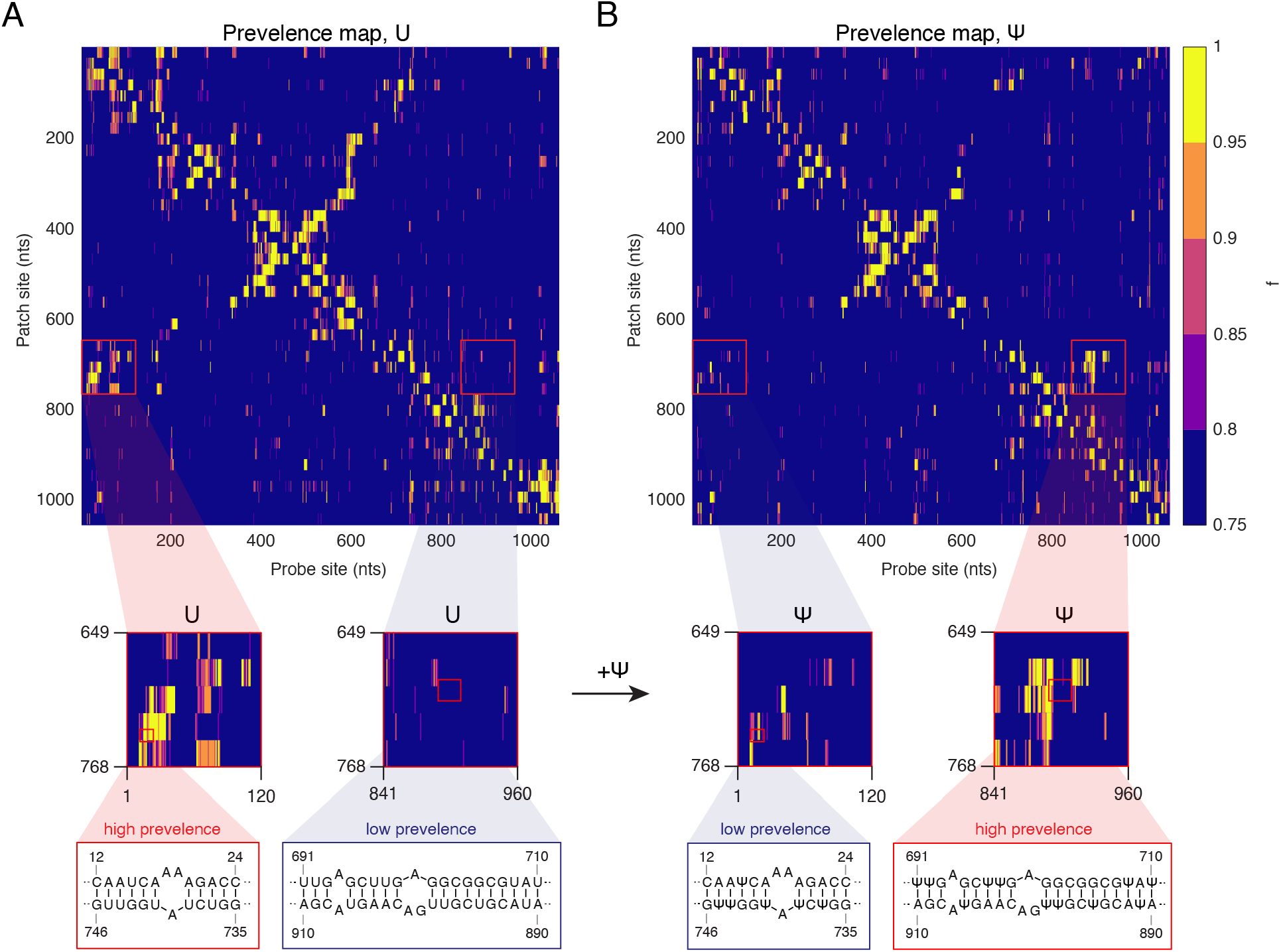
Normalized coupling signals provide information about the prevalence of connections in the population of STMV RNA molecules and enable comparisons between different experimental conditions. **A.** The map of normalized coupling signals for STMV RNA, which is corrected for patch and probe affinities (**Methods**), provides a measure of the prevalence of connections in the population. **B.** The map of normalized signals for STMV RNA containing pseudouridine (Ψ) reveals differences in the prevalence of certain connections relative to RNA containing normal uridine (U). Specifically, highly prevalent connections between patch sites at 649-768 nt and probe sites at 1-120 nt in U-containing RNA are less prevalent in Ψ-containing RNA. In contrast, low prevalence connections between patch sites at 649-768 nt and probe sites at 1-120 nt in U-containing RNA show higher prevalence in Ψ-containing RNA. These differences are highlighted by red boxes. Partially complementary sequences in these regions suggest specific connections that may be present. See **Extended Data Fig. 8** for additional plausible connections.

### The method reveals structural changes induced by modified nucleotides

Because the prevalences 𝑓_𝑖𝑗_ are corrected for patch and probe affinities, we can compare their values across experiments, enabling us to measure changes in RNA connectivity in response to changing conditions. To demonstrate this point, we apply the method to STMV RNA containing the modified nucleotide pseudouridine (Ψ) in place of normal uridine (U). Pseudouridine, an important component of natural RNA molecules^45^ and synthetic RNA vaccines^46^, forms stronger interactions with other nucleotides than uridine does^47^. These interactions are known to stabilize short duplexes^48^, but their effect on the connectivity of long RNA molecules is not understood. By comparing patch-probe experiments on STMV RNA molecules with and without Ψ, we aim not only to demonstrate that the method can detect changes in connectivity, but also to measure which connections are affected by the modified nucleotides.

The prevalence map of Ψ-containing molecules (**Fig. 4B**) shows some features similar to those of unmodified STMV RNA, including the central ridge of the T-domain, but also shows some differences. In particular, we observe a decrease in one cluster of long-range couplings (**Fig. 4A**), and the appearance of a new cluster of shorter-range couplings (**Fig. 4B**). The long-range couplings (involving regions of the sequence between nucleotides 1-100 and 650-750) are consistent with connections reported by Athavale *et al.* (**Fig. 3D, ellipse 4**), and the shorter-range couplings (involving the region between nucleotides 670-920) are consistent with connections predicted by RNAfold^15^ (**Extended Data Fig. 6**). Thus, the incorporation of Ψ appears to shift the connectivity of nucleotides 670–750 away from the longer-range connections toward the shorter-range connections, signifying a qualitative change in structure.

These changes occur in regions of the sequence that are thought to adopt functional structures. Downstream of nucleotide 700, the STMV sequence has high homology to the RNA of its helper virus, tobacco mosaic virus (TMV), which is known to fold into a functional transfer-RNA-like structure^49^ flanked by multiple short-range pseudoknots^50^. By detecting changes in the connectivity of this region, our results suggest structural changes that might affect the functionality of STMV RNA. These results could be used to inform *in vivo* studies that directly test the effect of Ψ on the biological properties of STMV. Similar comparative studies on Ψ-containing molecules used in RNA vaccines could reveal structural changes that shed light on their unique biological properties, such as their enhanced translational capacity and reduced immunogenicity^46^.

## Discussion

Throughout, we have been careful to distinguish between connectivity and structure. Two-dimensional methods—including multidimensional probing, proximity ligation, and the patch-probe method described here—measure intramolecular connections. If there were only one secondary structure, then the connections measured by these methods would determine that structure to within the resolution of the technique. But if there are variations in secondary structure within the population of molecules, these methods reveal only the connections that are present in the population. They do not reveal how those connections are grouped together into structures nor how many different structures exist, though such information could be gleaned through additional modeling and assumptions^50–52^.

Nonetheless, information about the connectivity of long RNA molecules can provide important clues about their biological function. For example, the RNA genomes of certain viruses form connections that are thought to direct a range of functions, including the production of viral proteins^54^ and the replication of new viral RNA strands^55^, and there is growing evidence that these connections can rearrange in response to changing conditions^56–59^, possibly triggering changes in functionality. Quantifying the prevalence of connections is therefore an important step in understanding how RNA virus genomes orchestrate infections, and could inform strategies for blocking infections by pathogenic viruses.

We use the patch-probe method to infer the prevalence of connections within a population of STMV RNA molecules, revealing several new features of the connectivity, including the coexistence of global and local folds. Previous chemical probing studies predict qualitatively different folding patterns for STMV RNA depending on the folding model used to interpret the data, with thermodynamic models predicting globally folded structures with extensive long-range connections^38–40^, and kinetic models predicting exclusively locally folded structures^41, 43^. Our prevalence map (**Fig. 4A**), obtained without reference to theoretical folding models of any kind, contains features that agree well with the globally folded T-shaped domain reported by Athavale *et al*.^38^, Archer *et al*.^39^, and Larman *et al.*^40^. However, we also find features that are consistent with locally folded structures reported by Schroeder *et al.*^41^, indicating that both global and local folds coexist in solution. This observation highlights the need for a statistical description of RNA structures^60^, and for experiments that measure competing connections.

Proximity ligation techniques, such as PARIS^23^ and COMRADES^26^, can detect competing connections in long RNA molecules, and we therefore consider how the patch-probe method compares to these techniques. Proximity ligation has the major advantage that it can address multiple sequences in parallel and *in vivo*, whereas the patch-probe method currently only works with a single sequence *in vitro*. Furthermore, the signals measured in a proximity ligation experiment arise directly from RNA connections, whereas the signals in the patch-probe experiment reflect perturbations to the RNA connectivity and are therefore less direct. However, inferring the prevalence of connections from proximity ligation data is not always straightforward. In principle, the magnitude of the ligation signal should scale with prevalence, but in practice some steps of the protocol, including ligation and PCR, can bias the measured signals in ways that are difficult to correct for^22, 52^. Inferring the prevalence from patch-probe data is simpler: after correcting for the patch and probe affinities, the magnitude of each coupling peak provides a quantitative measure of the prevalence.

To interpret the prevalence values we assume that each coupling peak corresponds to a pairwise connection in the RNA molecules. While the approximate symmetry of the 2D data (**Figs. 1F, 2, 3A, and 4**) suggests that this assumption holds for many of the couplings measured in the experiment, there are scenarios in which the assumption might not hold, which could lead to the identification of spurious connections. For example, in regions of the RNA sequence directly flanking the patch site, binding of the patch oligonucleotide could affect nearby probe binding even if those regions are not directly connected to the patch site. Furthermore, patch binding could induce nonlocal rearrangements of the RNA connectivity, involving cascades of multiple connections.

These scenarios highlight an important feature of the method: DNA oligonucleotides do not merely report on RNA structure; they can also modify it. The ability to modify and interact with the folded structure of an RNA molecule could enable new ways of measuring collective aspects of its folding process. Much as how DNA origami uses many staple oligonucleotides to direct a long DNA molecule to fold into prescribed structures^61^, it is possible that multiple patches could be used to drive a long RNA molecule away from its native folds in order to address larger regions of the folding landscape. For example, we imagine perturbing the RNA molecule in a way that mimics the unfolding and refolding effects of cellular enzymes during biological processes like transcription.

## Methods

### Buffers

**Hybridization buffer:** 50 mM Tris-HCl, pH 7.0; 1 M NaCl; 1 mM EDTA; 0.5% Tween-20. **TE buffer:** 10 mM Tris-HCl, pH 7.0; 1 mM EDTA. **TAE buffer:** 40 mM Tris-HCl, pH 8.3; 20 mM acetic acid; 1 mM EDTA.

### Preparation of fluorescently-labeled STMV RNA

We prepare fluorescently-labeled STMV RNA by *in vitro* transcription, using a DNA template derived from a plasmid containing the STMV sequence^38^ (gift from Steve Harvey, University of Pennsylvania). The plasmid contains a T7 promoter sequence upstream of the STMV sequence, and a HindIII restriction site downstream of the STMV sequence. We digest the plasmid with Hind III (New England Biolabs) to generate a linear template, purify the template by acid phenol–chloroform extraction and ethanol precipitation, and resuspend the template in molecular biology grade water. The sequence of the template is verified by Sanger sequencing (Genewiz).

We transcribe STMV RNA from the linear DNA template using a TranscriptAid T7 High Yield Transcription Kit (Thermo Fisher). A small amount of AlexaFluor-546 UTP (Thermo Fisher) is added to the transcription reaction (**Table 1**), such that the final RNA transcripts contain roughly one dye per transcript, as measured by UV-Vis spectrophotometry (**Supplementary Fig. S1**). To prepare STMV RNA containing pseudouridine (Ψ), we replace the non-fluorescent UTP in the transcription reaction with ΨTP (TriLink BioTechnologies).

**Table 1.** Transcription reaction. Components of the in vitro transcription reaction used to prepare fluorescently-labeled STMV RNA from a linearized plasmid DNA template.

| Reagent | Volume |
| --- | --- |
| Water | to 20 μL |
| Template DNA, 1 μg/μL | 1 μL |
| 5X TranscriptAid Reaction Buffer | 4 μL |
| ATP, 100 mM | 2 μL |
| UTP, 100 mM | 2 μL |
| CTP, 100 mM | 2 μL |
| GTP, 100 mM | 2 μL |
| Alexa Fluor 546-14-UTP, 1mM | 2 μL |
| T7 TranscriptAid Enzyme Mix | 2 μL |

Following transcription we digest the DNA template with DNase I (New England Biolabs), and purify the RNA transcripts using an RNeasy mini kit (Qiagen), eluting in TE buffer. Then we wash the RNA 5 times with additional TE buffer using a 0.5 mL 100kDa centrifugal filter (MilliporeSigma). The purified transcripts migrate as a single band in a native 1% agarose gel prepared in TAE buffer, signifying that they are full-length and not degraded (**Supplementary Fig. S1**). Finally, we dilute the RNA to a concentration of 2 µM and store it at −80°C prior to use.

### Sequence of the STMV RNA transcript

The sequence of the RNA transcript is shown below. Uppercase letters correspond to the 1,058-nt consensus sequence of STMV genome, and lowercase letters are extra nucleotides that get incorporated during transcription.

gggAGUAAACUUACCAAUCAAAAGACCUAACCAACAGGACUGUCGUGGUCAUUUAUGCUGUUGGGGGACAUAGGGGGAAAACAUAUUGCCUUCUUCUACAAGAGGCCUUCAGUCGCCAUAAUUACUUGGCGCCCAAUUUUGGGUUUCAGUUGCUGUUUCCAGCUAUGGGGAGAGGUAAGGUUAAACCAAACCGUAAAUCGACGGGUGACAAUUCGAAUGUUGUUACUAUGAUUAGAGCUGGAAGCUAUCCUAAGGUCAAUCCGACUCCAACGUGGGUCAGAGCCAUACCUUUCGAAGUGUCAGUUCAAUCUGGUAUUGCUUUUAAAGUACCGGUCGGGUCACUAUUUUCGGCAAAUUUCCGGACAGAUUCCUUUACAAGCGUCACAGUGAUGAGUGUCCGUGCUUGGACCCAGUUAACACCGCCAGUAAAUGAGUACAGUUUUGUGAGGCUGAAGCCAUUGUUCAAGACUGGUGACUCUACUGAGGAGUUCGAAGGGCGUGCAUCAAACAUCAACACACGAGCUUCUGUAGGGUACAGGAUUCCAACUAAUUUGCGUCAGAAUACUGUGGCAGCCGACAAUGUAUGCGAAGUAAGAAGCAACUGUCGACAAGUCGCCUUGGUUAUUUCGUGUUGUUUUAACUGAACCUCGACAUAAGCCUUUUGGAUCGAAGGUUAAACGAUCCGCUCCUCGCUUGAGCUUGAGGCGGCGUAUCUCUUAUGUCAACAGAGACACUUUGGUCUAUGGUUGUAUAACAAUAGAUAGACUCCCGUUUGCAAGAUUAGGGUUAACAGAUCUUGCCGUUAGUCUGGUUAGCGCGUAACCGGCCUUGAUUUAUGGAAUAGAUCCAUUGUCCAAUGGCUUUGCCAAUGGAACGCCGACGUGGCUGUAUAAUACGUCGUUGACAAGUACGAAAUCUUGUUAGUGUUUUUCCCUCCACUUAAAUCGAAGGGUUUUGUUUUGGUCUUCCCGAACGCAUACGUUAGUGUGACUACCGUUGUUCGAAACAAGUAAAACAGGAAGGGGGUUCGAAUCCCUCCCUAACCGCGGGUAAGCGGCCCAa

### Design of the DNA patches

We design the DNA patches to be complementary to the STMV genome and to collectively tile its primary sequence. We use 44 patch oligonucleotides of length 24 nt that tile the STMV genome from nucleotide 1 to 1,056. The patch sequences are listed in **Table 2**.

**Table 2.**
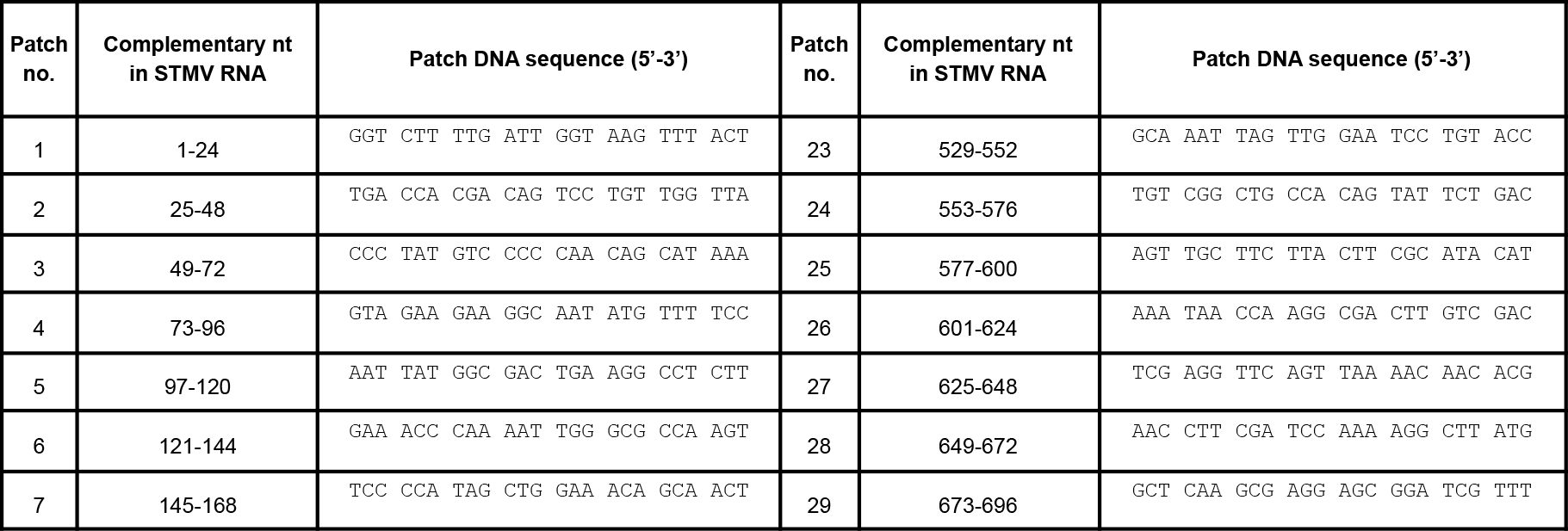
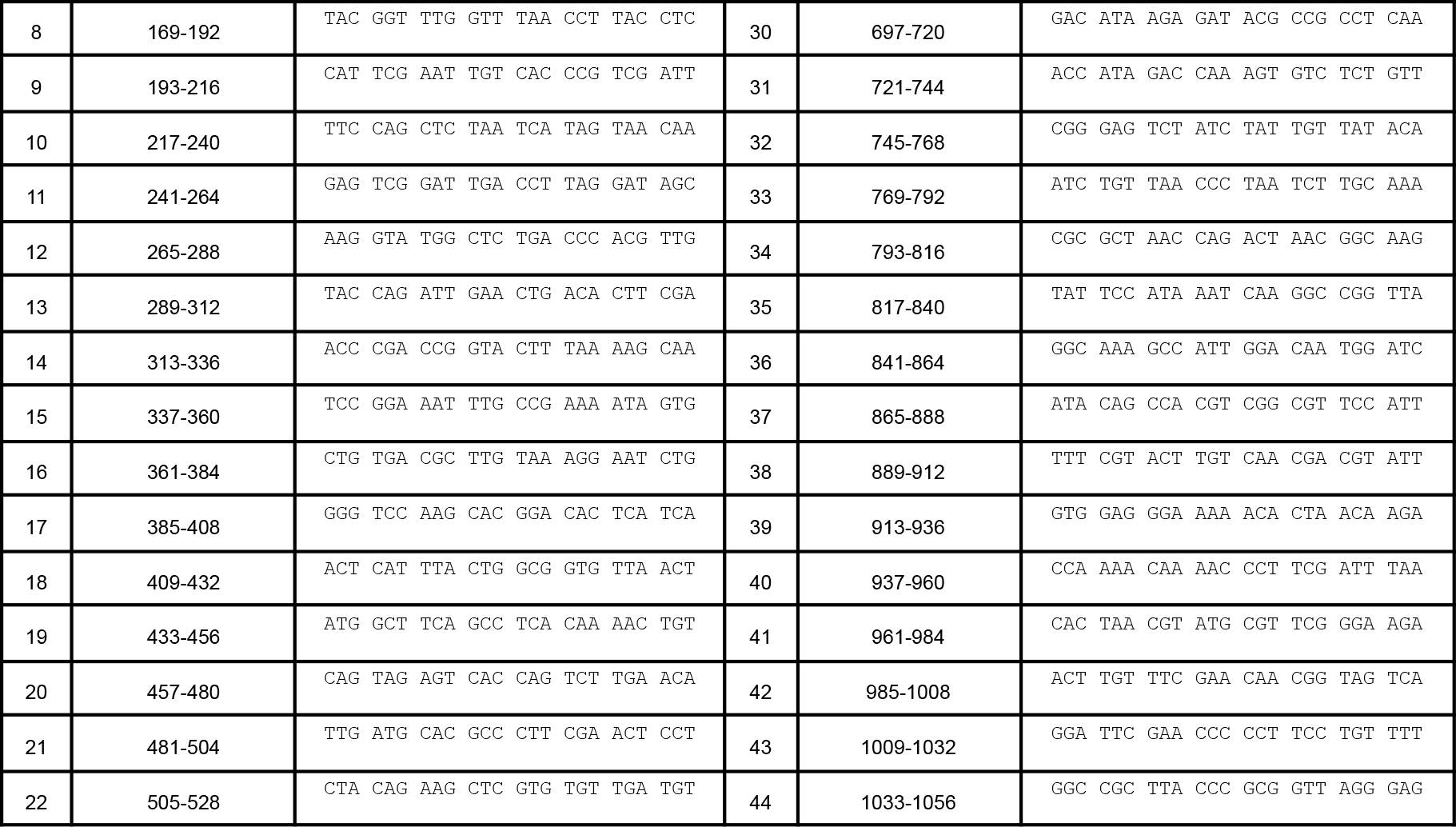
DNA Patches. The number, complementary nucleotides in the STMV RNA genome, and sequence of the 44 DNA patch oligonucleotides.

### Design of the DNA microarray and probes

The DNA microarrays used in our experiments are manufactured by Agilent Technologies (Supplier Item G4860A, SurePrint G3 Custom GE 1x1M Microarray). The arrays are prepared on 76 mm × 25 mm glass slides. Each slide contains 974,016 features arranged in a hexagonal grid with 1068 rows and 912 columns. Each feature is a circular spot with a nominal diameter of 30 micrometers in which many identical DNA oligonucleotides are tethered by their 3’-ends to the glass surface. Each oligonucleotide is 60 nt long.

We design two sets of DNA probes for the microarray: 12-nt probes and 24-nt probes. The 5’-ends of the probes contain either 12 or 24 nt that are complementary to the STMV genome, and the 3’-ends are poly-T spacers. We design each set of probes such that the complementary regions cover the entire STMV genome: there are 1,047 12-nt probes and 1,035 24-nt probes. The probes are numbered according to the first nucleotide in the STMV sequence that they are complementary to. As examples, the first and last three 12- and 24-nt probes are shown in **Table 3**. The complete list of probes is given in **SI**.

**Table 3.**
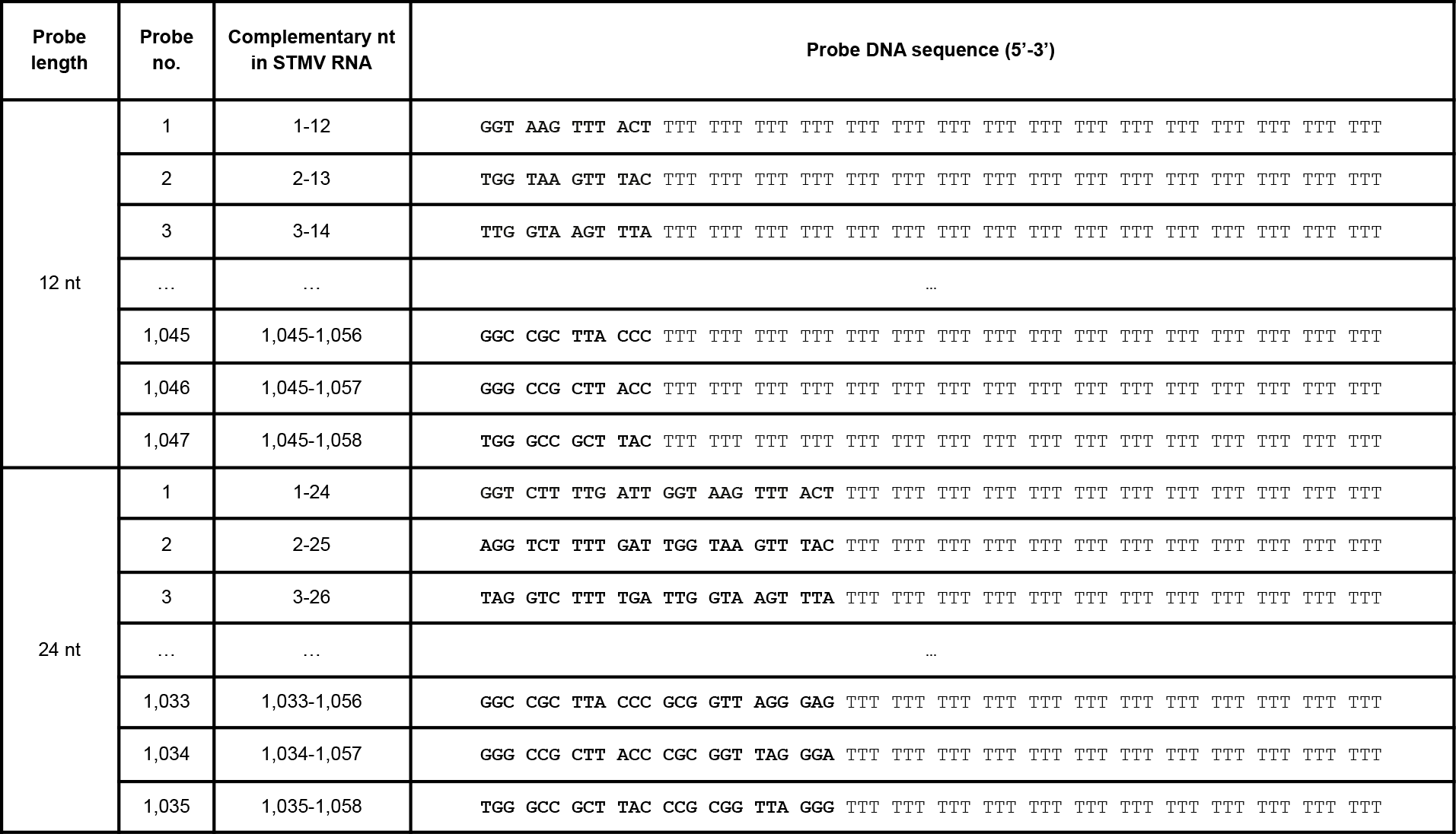
12- and 24-nt probes. The length, number, complementary nucleotides in the STMV RNA genome, and sequence (with complementary region in bold) of several DNA microarray probes. The complete list is given in **SI**.

To perform multiple patch-probe experiments on the microarray, we divide the array into identical subarrays that contain 3 replicates of each probe. Each subarray has 149 rows and 43 columns, for a total of 6407 spots. Of these spots, (3×1047 + 3×1035 =) 6246 are assigned to the 12- and 24-nt probes and their replicates. The remaining spots are assigned to various control oligonucleotides, including poly-T oligonucleotides, poly-A oligonucleotides, and oligonucleotides that are complementary to the start and end of the transcript but only partially complementary to the STMV genome. We position the poly-T and poly-A controls in the corners of the subarray, and assign random positions to the probes and their replicates, as well as the remaining controls. Because the nominal spacing between rows is 0.018330871 mm and between columns is 0.06349 mm, the subarrays are roughly square, with a height and width of 2.73 mm (**Supplementary Fig. S2**). We arrange the subarrays in a square grid across the microarray. The grid has 21 complete columns and 7 complete rows, for a total of 147 complete subarrays (**Supplementary Fig. S2**). The position of every spot and its corresponding probe sequence are provided as an XML file in **SI**.

### Design of the microarray gasket

We design a gasket to distribute the RNA samples to different subarrays. The gasket holds at least 10µL of sample volume and covers an area on the microarray that is greater than that of the subarray. Specifically, the wells cover 3.2 mm x 3.2 mm, with a separation of 1 mm between wells, and a separation of 0.5 mm from the edge of the array (**Supplementary Fig. S3**). A grid of wells consisting of 4 rows and 12 columns could fit on a single array.

A negative mold of the gasket is designed using FreeCAD (http://www.freecadweb.org) and printed using a Formlabs Form 3 3D printer (Formlabs). We cast the gaskets in the negative mold using polydimethylsiloxane (PDMS). A mixture of 10 parts DOWSIL 184 silicone elastomer base (Dow) and 1 part DOWSIL 184 silicone elastomer curing agent (Dow) are mixed vigorously, degassed under vacuum, and then poured into the mold. We cure the PDMS by incubating at 65°C overnight.

### Design of the hybridization clamp

We design a hybridization clamp to hold together the microarray slide and PDMS gasket (**Supplementary Fig. S4**). The clamp is designed using FreeCAD and printed using a Formlabs Form 2 3D printer. We fasten the clamp with 4 1”-binder clips (ACCO) (**Supplementary Fig. S5**). The handles of the binder clips are removed after assembly.

### Hybridization of the RNA and the DNA patches

In parallel, we mix the STMV RNA transcripts and each DNA patch in a 1:1 molar ratio in hybridization buffer such that the final concentration of RNA (and patch) is 10 nM and the final volume is 20 µL. We also add 24-nt poly-T DNA oligomers to a final concentration of roughly 1 µM, which reduces non-specific binding of the RNA to the poly-T spacers of the DNA probes during hybridization to the microarray. Before adding the RNA-DNA mixtures to the microarray, the mixtures are heated to 90°C and then cooled to 4°C at a rate of −1°C/s. This heating and cooling procedure breaks up RNA aggregates that can form during *in vitro* transcription and also helps drive hybridization of the RNA and the DNA patches.

As a control experiment, we perform RNA-patch hybridization without heating and cooling, by mixing 10-fold higher concentrations of RNA and patch (100 nM each) in hybridization buffer and incubating at 37°C for 1 h. Prior to patch hybridization, the RNA sample is heated and cooled by itself to break up aggregates and reproduce the thermal refolding process. Following patch hybridization but prior to adding the samples to the microarray, we dilute the concentration of RNA (and patch) back to 10 nM. This no-heat hybridization control gives similar results to the thermal hybridization experiments, as shown in **Supplementary Figs. S10, S11, S15, S17.**

### Hybridization of the RNA and the DNA microarray probes

We add the RNA-DNA mixtures to the microarray using a custom-built gasket. The gasket is made of polydimethylsiloxane (PDMS) and contains a 4 x 12 square grid of 48 wells. Each well covers a 3.2 mm x 3.2 mm square, and the separation between wells is 1 mm (**Supplementary Fig. S3**). We load 10 µL of each RNA-DNA mixture into its own well and place the microarray slide DNA-side down on top of the gasket. At this point in the experiment, the mixtures are not in contact with the array. We then clamp the microarray and gasket together using a custom-built chamber (**Supplementary Fig. S4 and S5**), invert the chamber, and spin it in a swinging bucket centrifuge at 1000 rcf for several seconds to bring the RNA-DNA mixtures in contact with the array. The microarray is incubated at 37°C for 100 minutes, washed for 1 minute with hybridization buffer, and dried completely.

### Imaging the microarray

We image the fluorescence of the microarray using an Agilent SureScan Microarray Scanner with the resolution set to 2 µm and the bit depth set to 20 (**SI**). The fluorescence integrated over each spot is determined using Agilent FeatureExtraction Software, using a non-standard extraction protocol provided by the Agilent development team upon request (**SI**). We also determined integrated fluorescence for each spot using our own image analysis protocol written in MATLAB (**SI**). The prevalences inferred with these different feature extraction protocols are in agreement, as shown in **Supplementary Fig. S18**.

### Compiling 1D and 2D binding spectra

We compile 1D and 2D binding spectra by averaging the integrated fluorescence over all spots corresponding to a given probe. For the 1D binding spectra (**Fig. 1E and Extended Data Fig. 2A**), we average over spots from four sections of the microarray containing replicate samples of unpatched RNA. For the 2D binding spectra (**Fig. 1F and Extended Data Fig. 2A**), each patch is added to its own section and we average over spots from that section only.

### Inferring the coupling signals and prevelances

A detailed account of the inference procedure is described in **SI Supplementary Methods**. Graphical depictions of the Bayesian models for inferring the signals and prevalences are shown in **Supplementary Figs. S6 and S7.** Detailed results of the MCMC approach to infer the signals are shown in **Supplementary Figs. S8-S13** and in **Supplementary Tables S1-S6**.

Detailed results of the MCMC approach to infer the prevalences are shown in **Supplementary Figs. S14-S16** and in **Supplementary Tables S7-S9**. A comparison of inference results from experiments involving annealed and unannealed patches is shown in **Supplementary Fig. S17.** A comparison of inference results from data in which features are extracted using Agilent’s software and our own software is shown in **Supplementary Fig. S18**. A comparison of inference results from two replicate experiments on the same RNA sequences is shown in **Supplementary Fig. S19**.

### Identifying dominant couplings

For each patch, we identify the top 5 probes with the highest signal, 𝑆_𝑖𝑗_. We define these patch-probe combinations as the dominant couplings. We plot each dominant coupling as a rectangle that spans the patch and probe binding sites in **Fig. 3A**.

### Measuring patch affinities by gel electrophoresis

We validate the patch affinities inferred from the microarray data by comparing them to affinities measured in bulk using gel electrophoresis. In contrast to the microarray measurements, the bulk measurements are performed using DNA patches that are fluorescently labeled and STMV RNA molecules that are unlabeled. The 5’-ends of the bulk patches are the same as the microarray patches, but the 3’-ends contain an additional 6-T spacer sequence terminated by a fluorescein dye (3’ 6-FAM, Integrated DNA Technologies). The 6-T spacer is designed to reduce nucleobase quenching of the fluorescence signal that can occur when the fluorescently-labeled patch binds to the RNA. To perform the bulk binding measurements we mix together 300 nM of STMV RNA with 300 nM of fluorescently-labeled patch in hybridization buffer. In one experiment, we heat the sample to 90°C, cool the sample to 4°C at a rate of −1°C/s, and then incubate the sample for 100 minutes at room temperature. This protocol corresponds to our normal patch-probe microarray experiment. In another experiment, we heat and cool the RNA molecules before adding the patches, and then incubate the mixture of RNA and patch for 100 minutes at room temperature. This protocol corresponds to our no-heat control. Following incubation, we perform native 1% agarose gel electrophoresis in TAE buffer and image the fluorescence of the gel (**Supplementary Fig. S20**). The slower-running fluorescence signal corresponds to bound patch, and the faster-running band corresponds to unbound patch. The patch affinity is calculated by dividing the bound signal by the total (bound plus unbound) signal (**Supplementary Fig. S21**). Comparisons of the microarray measurements and the bulk measurements, corrected for concentration according to the procedure described in **Supplementary Methods**, are shown in **Supplementary Figs. S22-S23**).

## Acknowledgments

We thank Ben Rogers and Sean Eddy for many helpful discussions. We thank Valentina Maran and Scott Leppanen from Agilent for their help with array design, and Steve Harvey for sharing the plasmid containing the STMV sequence. Research reported in this publication was supported by the National Institute of General Medical Sciences of the NIH (award number R00GM127751) (R.F.G.), the Peter B. Lewis ’55 Lewis-Sigler Institute/Genomics Fund through the Lewis-Sigler Institute of Integrative Genomics at Princeton University and the National Science Foundation through the Center for the Physics of Biological Function (PHY-1734030) (O.K.), the Office of Naval Research (N00014-17-1-3029) and the Simons Foundation (O.K. and M.P.B.), the NSF-Simons Center for Mathematical and Statistical Analysis of Biology at Harvard (1764269) and the Harvard Quantitative Biology Initiative, and the NSF through the Harvard University Materials Research Science and Engineering Center under NSF grants DMR-2011754. We acknowledge the California Metabolic Research Foundation for its support of biochemical research at San Diego State University, and the Bauer Core Facility at Harvard University for shared experimental facilities used in this study.

## Extended Data

**Extended Data Fig. 1:**
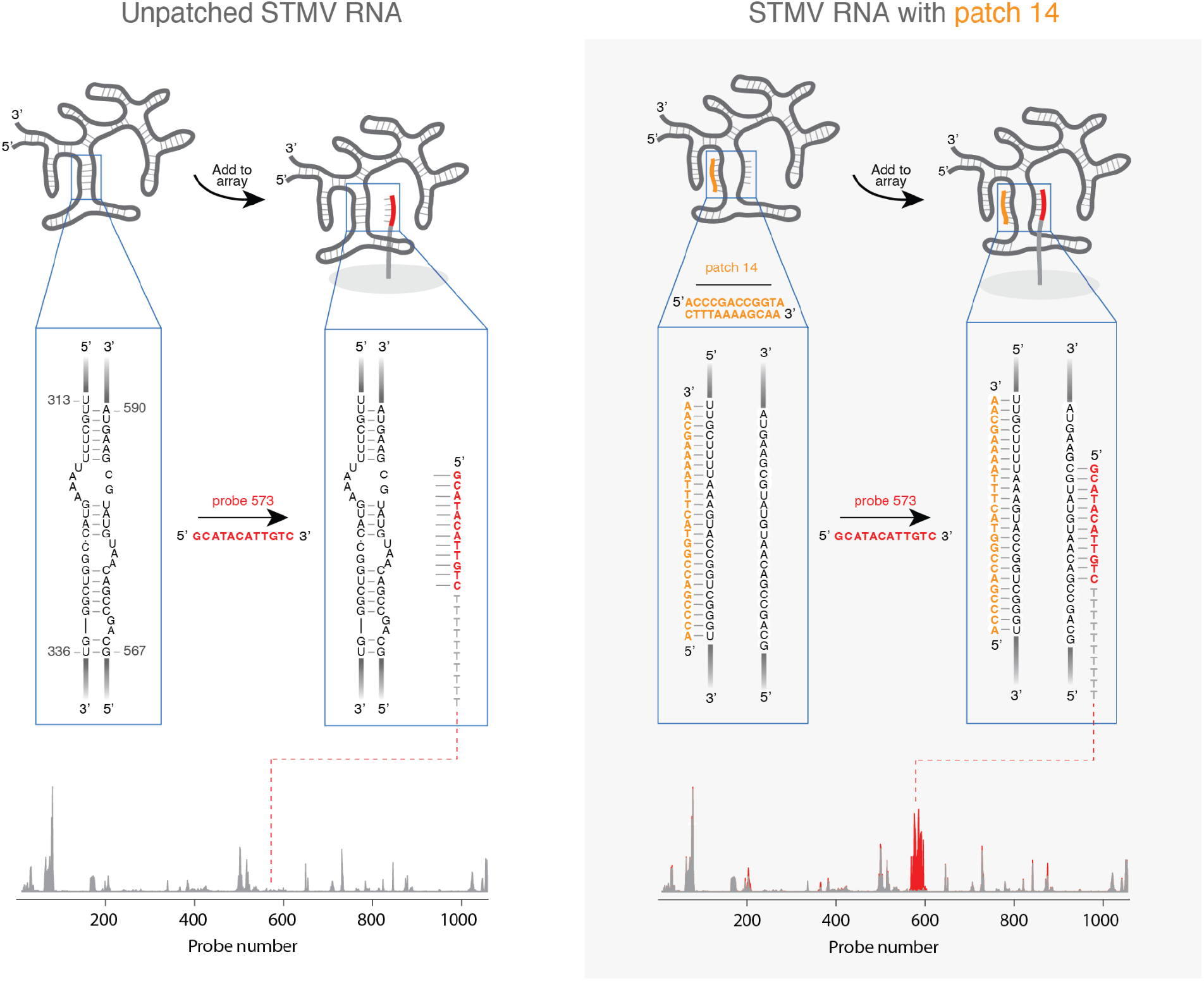
Plausible base pair interactions for the STMV RNA patch-probe experiment involving patch 14 and probe 573. Unpatched RNA binds weakly to microarray probe 573 owing to intramolecular RNA-RNA base pairs within the probe binding site. Hybridization of patch 14 disrupts these RNA-RNA base pairs, freeing up the binding site of probe 573 and increasing the binding of that probe.

**Extended Data Fig. 2:**
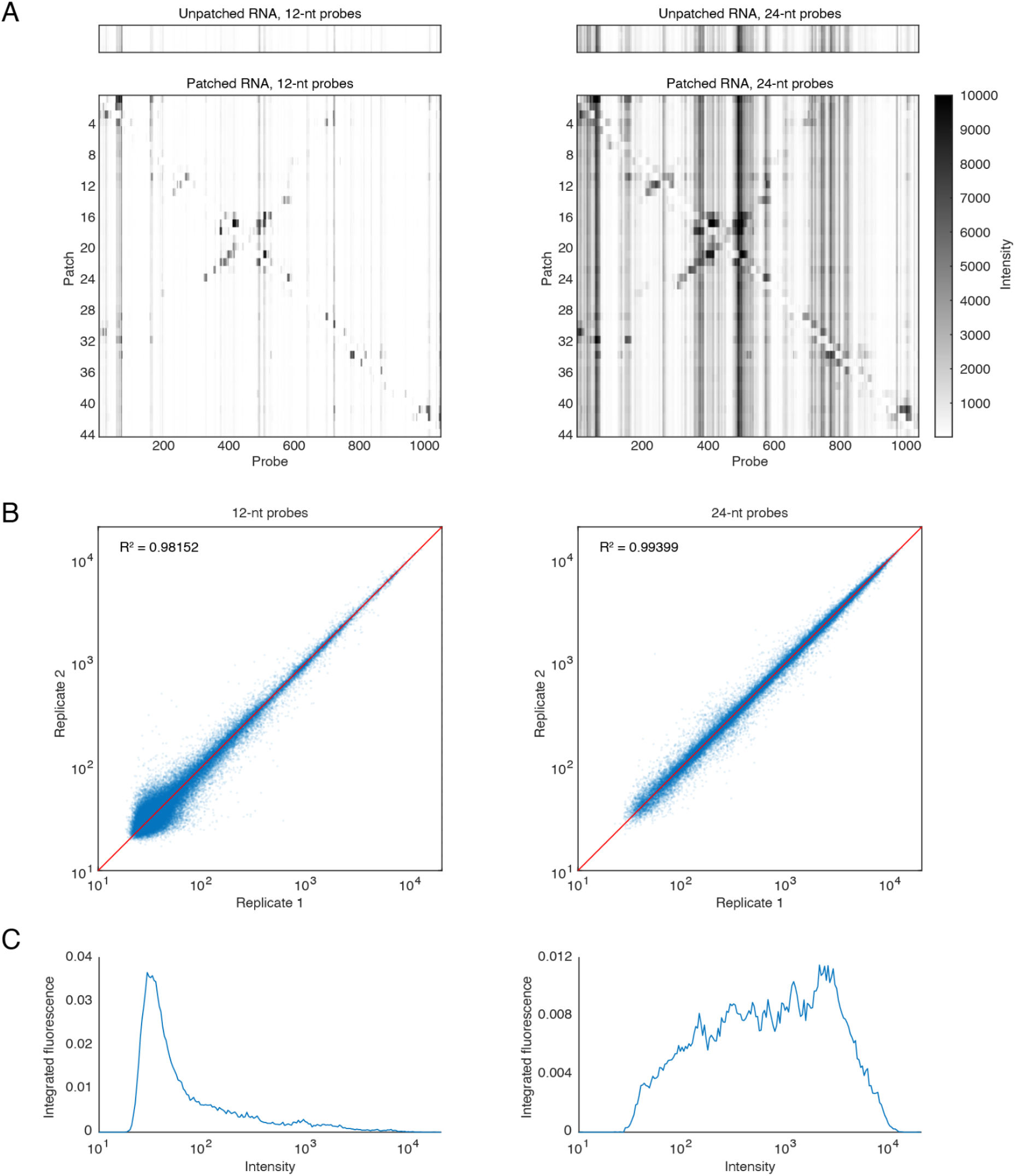
Binding spectra for 12- and 24-nt probes show reproducible changes in integrated fluorescence that span several orders of magnitude. **A.** Binding spectra of unpatched (top) and patched (bottom) STMV RNA. Spectra for both 12-nt (left) and 24-nt (right) long probes are shown. **B.** Replicate measurements of microarray spots are well correlated. **C.** Histograms of the microarray spot fluorescence measurements show that the dynamic range is several orders of magnitude. The measurements for 12-nt long probes are on average smaller than those of the 24-nt long probes.

**Extended Data Fig. 3:**
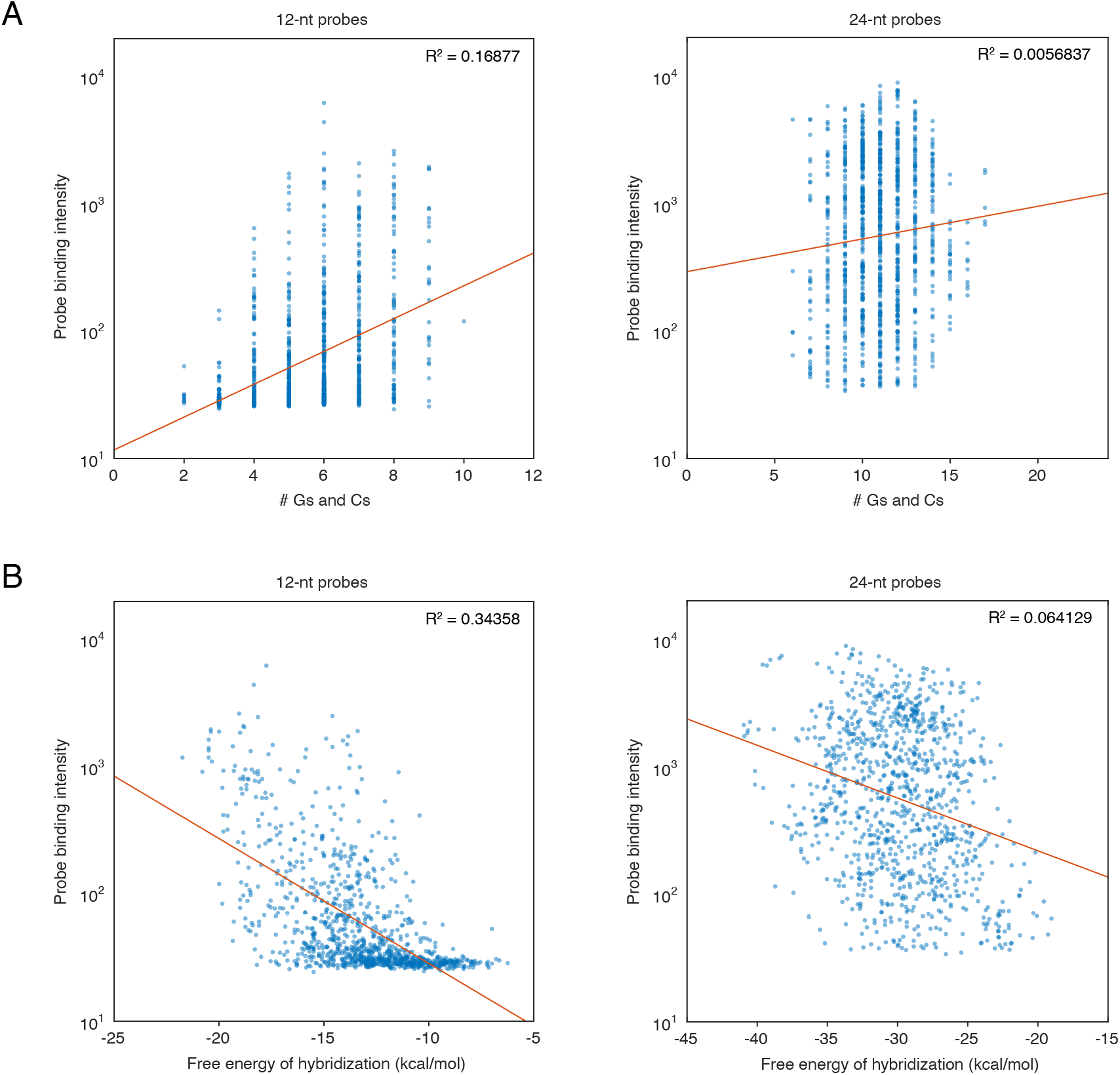
Probe GC content and hybridization free energy are not predictive of microarray binding. **A.** Microarray binding correlates weakly with the number of G and C nucleotides in the 12- (left) and 24-nt probes (right). Each point represents an individual probe, and the red lines show the best linear fits to the data. We would expect strong correlation in the absence of RNA secondary structure. **B.** Binding correlates weakly with the predicted free energy of hybridization in 12- (left) and 24-nt probes (right). Here, too, we would expect strong correlation in the absence of secondary structure. To predict the free energy of hybridization, we use LandscapeFold^62^ to calculate the minimum free energy of the RNA-DNA duplex in the absence of intramolecular base pairing. These results are virtually unchanged when we consider an ensemble of structures instead of the single MFE structure.

**Extended Data Fig. 4:**
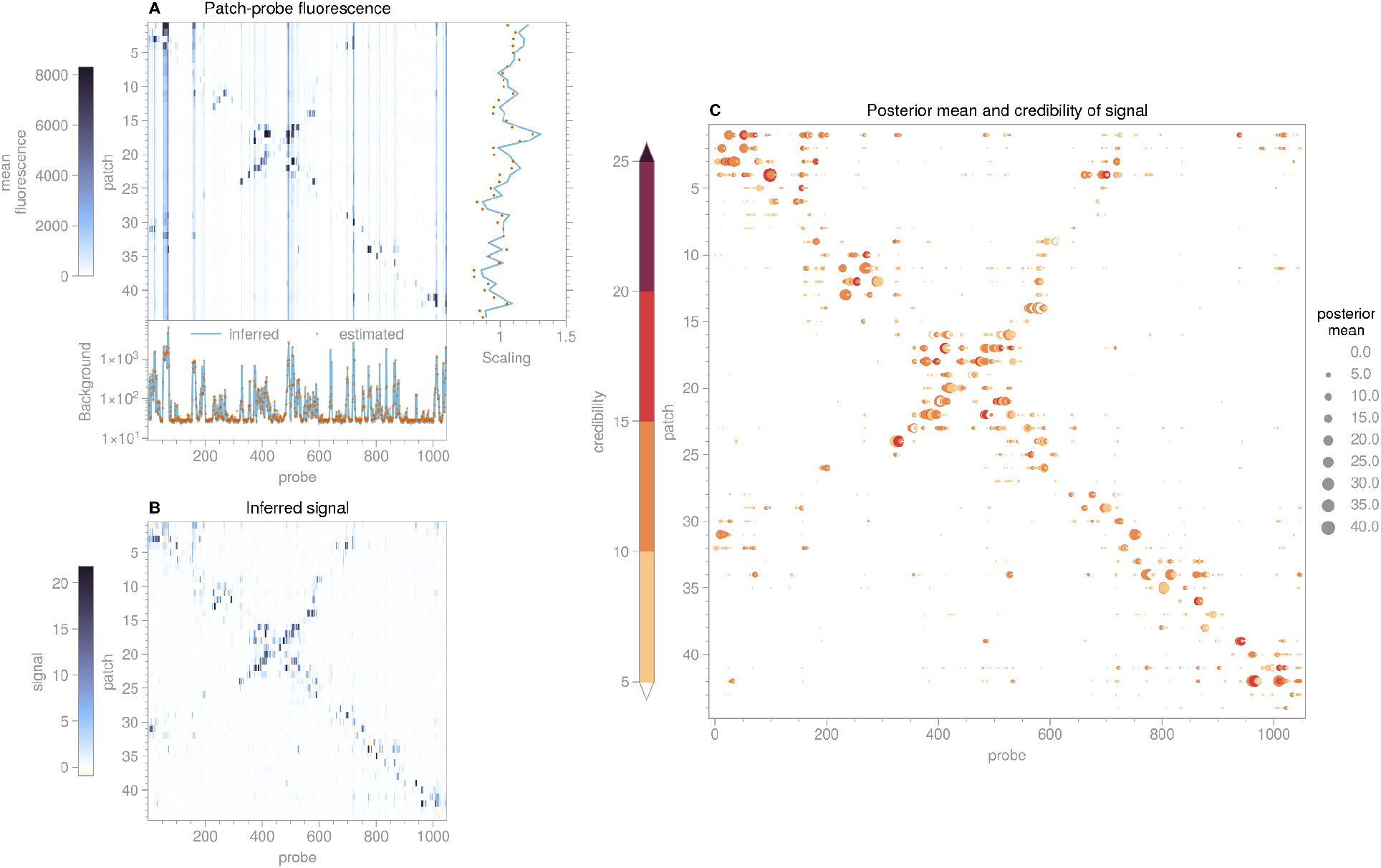
Signal for STMV RNA, 12-nt probes. **A.** Heatmap of patch-probe fluorescence data from microarray, averaged over spots for each (patch, probe) combination. Subplot at bottom shows the background estimated from a mean of the measurements taken in the four wells containing unpatched RNA, and the background inferred from the data using the Bayesian approach described in **SI**. Subplot at right shows the median values across all probes for each patch (dots) and the scaling values inferred from our Bayesian approach (blue line). **B.** Heatmap of signal inferred from the data in **A** using the Bayesian approach. In both **B** and **C**, the upper limit of the colormap corresponds to the 99.9th percentile of the values in the heatmap. **C.** Scatter plot showing the posterior mean (magnitude represented by size of symbols), and credibility (represented by color) of the inferred signals for each patch and probe combination. Only signal values with a credibility greater than 5 are shown.

**Extended Data Fig. 5:**
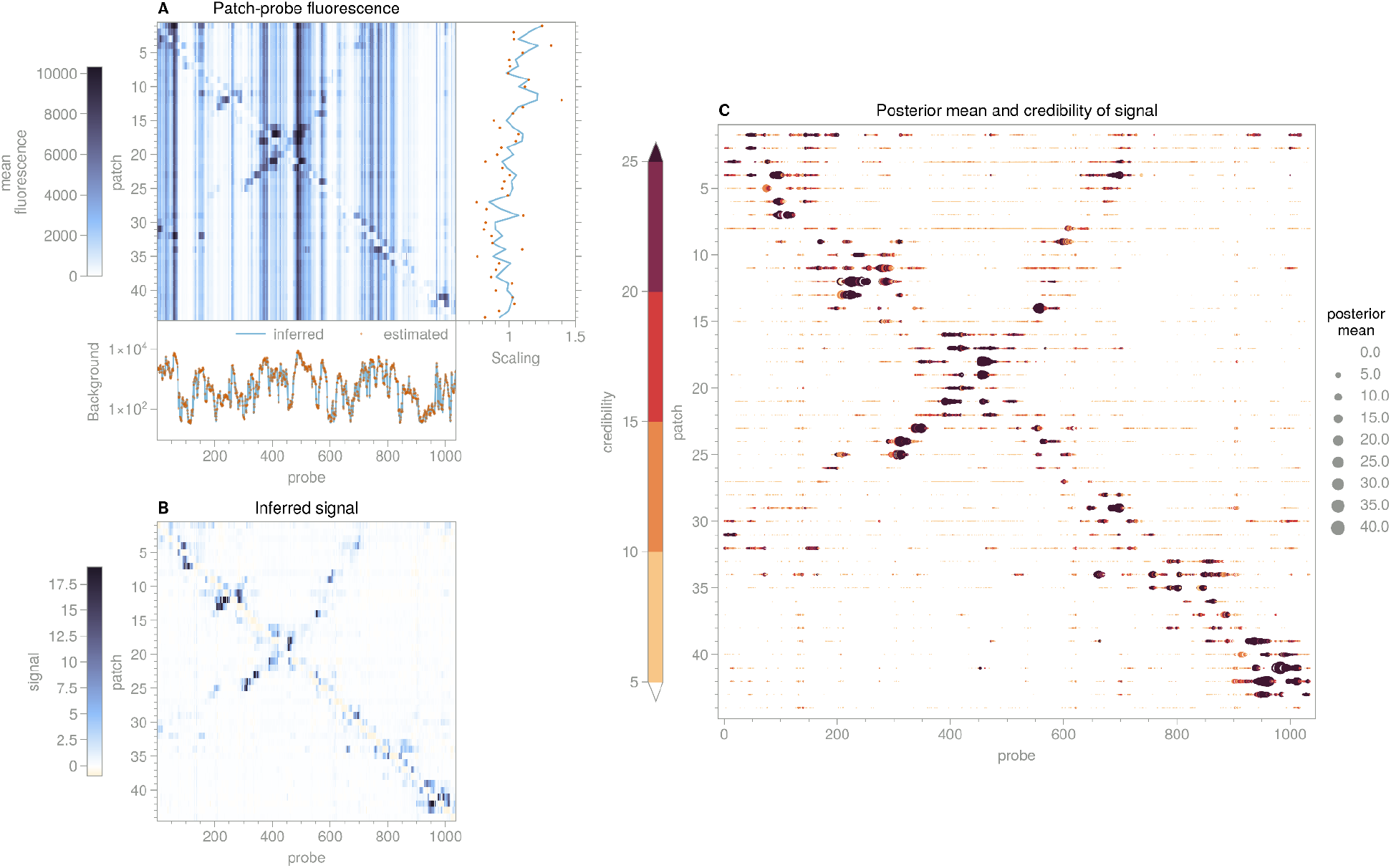
Signal for STMV RNA, 24-nt probes. See **Extended Data Fig. 4** for a detailed description of panels **A-C**.

**Extended Data Fig. 6:**
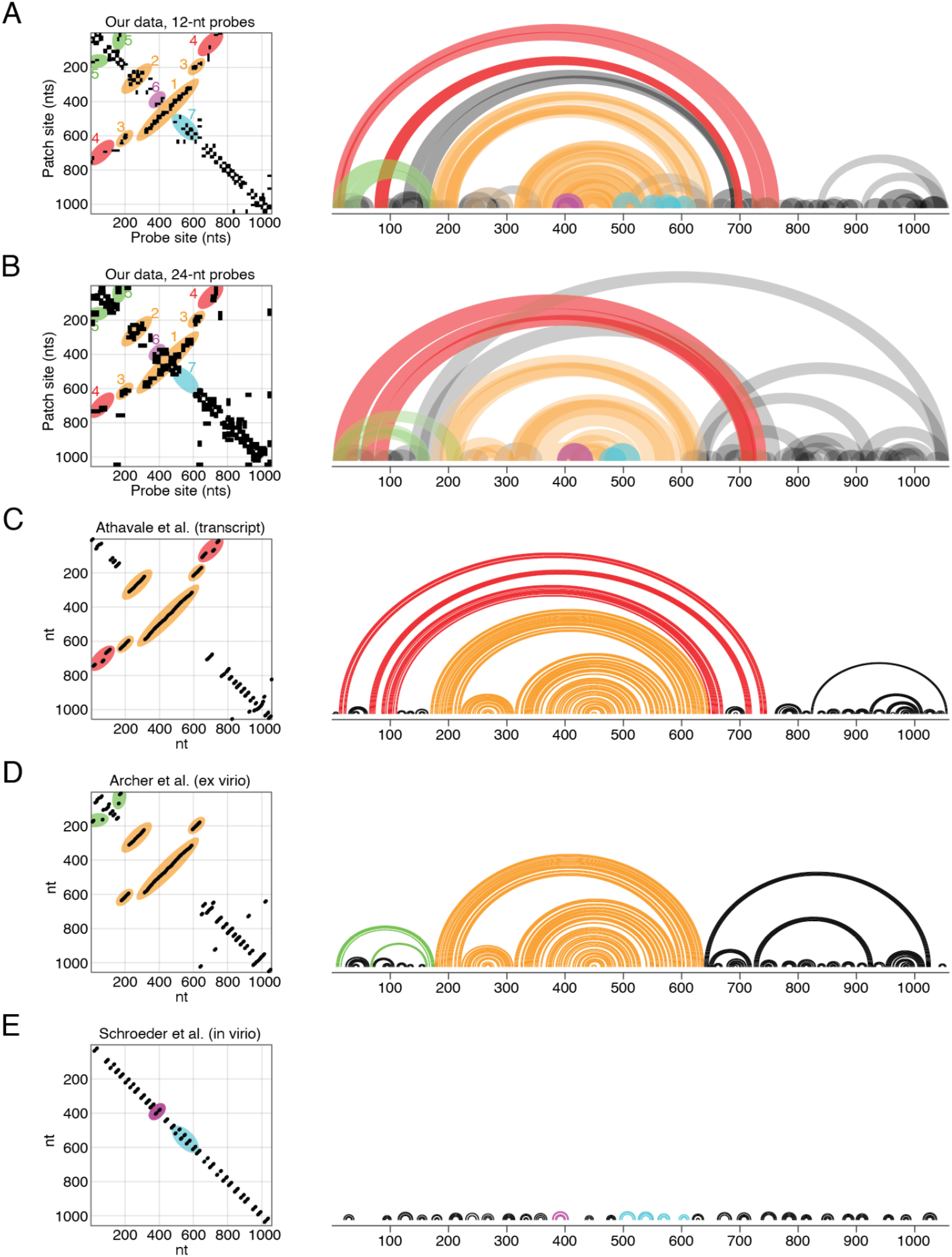
Many of the dominant patch-probe connections are consistent with those of previously reported structures. **A.** Matrix (left) and arc plot (right) representations of the dominant patch-probe couplings for 12-nt probes. The couplings within colored regions (1-7) of the matrix are shown with corresponding colors in the arc plot. **B**. Matrix and arc plot representations for the dominant patch-probe couplings for 24-nt probes, with the same coloring scheme as above. **C.** Dot and arc plot representations of the structure reported by Athavale *et al*. Connections in consensus regions 1-3 (orange) are present, as well as long-range connections in region 4 (red) spanning approximately 750 nucleotides. **D.** Dot and arc plot representations of the structure reported by Archer *et al*. Connections in consensus regions 1-3 (orange) are present, as are mid-range connections in region 5 (green) spanning approximately 170 nts. **E.** Dot and arc plot representations of the structure reported by Schroeder *et al*. Connections in regions 1-3 are not present. Rather, short range connections are present throughout the sequence, including regions 6 (purple) and 7 (cyan).

**Extended Data Fig. 7:**
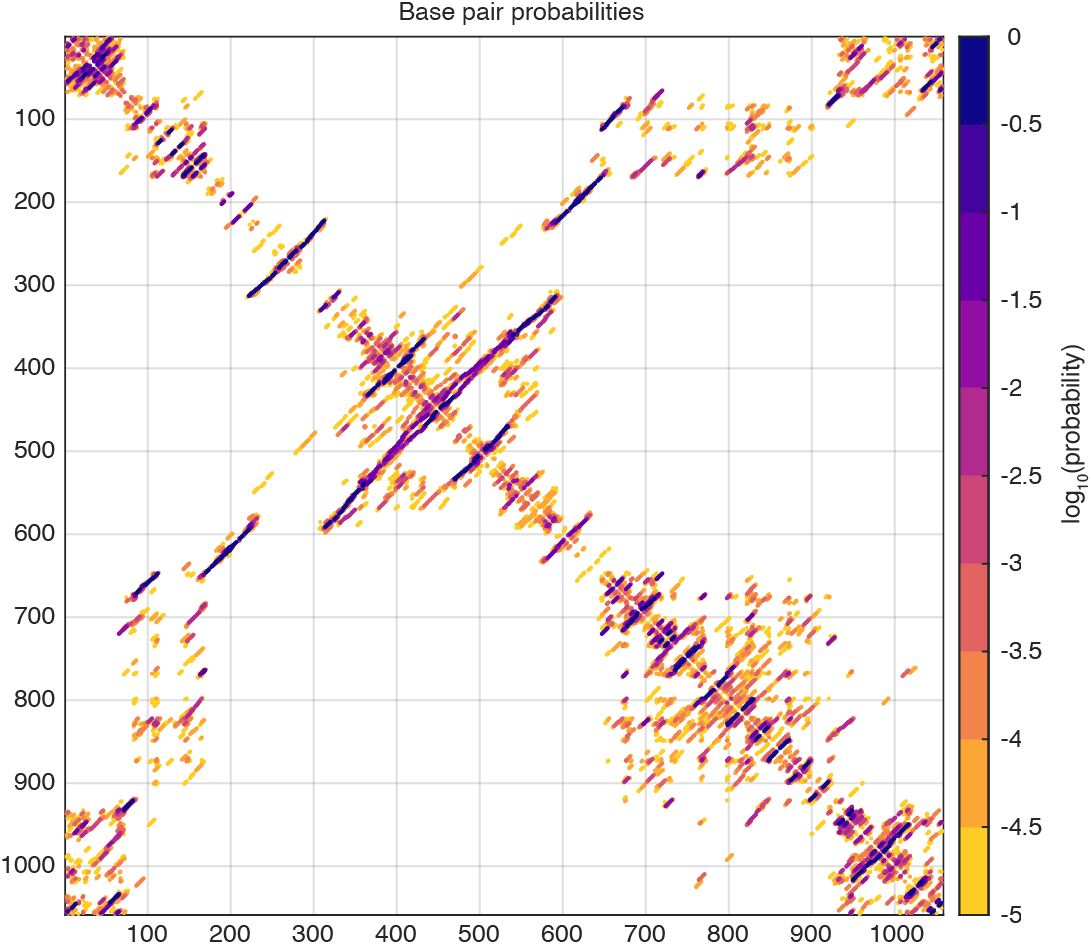
Equilibrium base pair probabilities for STMV RNA. Each base pair is shown as a dot whose color reflects the log_10_ of the pairing probability, as predicted by the thermodynamic folding algorithm RNAfold (ViennaRNA package^15^, version 2.4.17).

**Extended Data Fig. 8:**
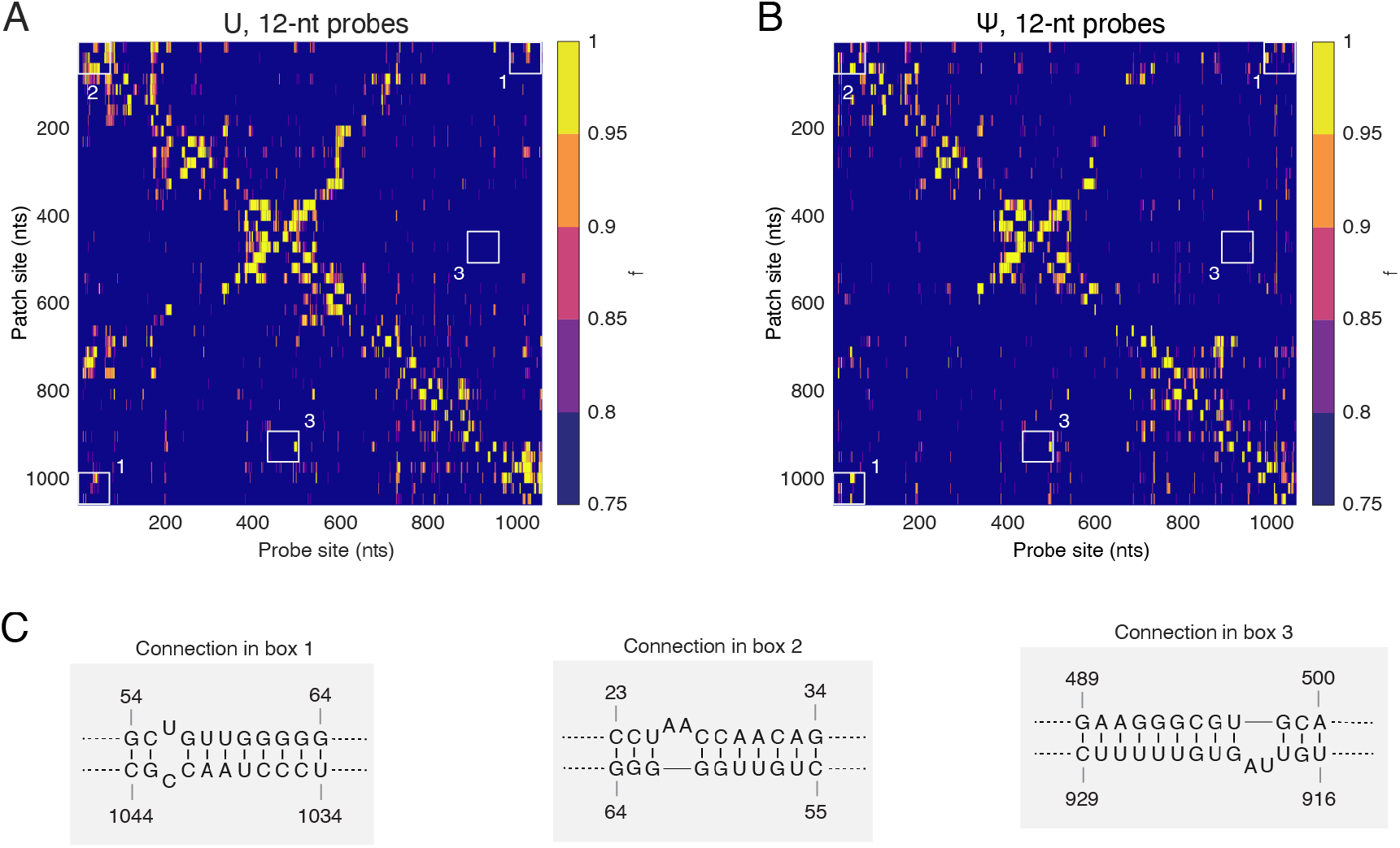
Several plausible connections in STMV RNA containing uridine (U) and pseudouridine (Ψ). **A.** Prevalence maps for U-containing RNA molecules inferred from measurements with 12-nt probes. )Boxes (1-3) highlight several features mentioned in the main text: (1) a very-long-range connection that is not observed in previous SHAPE studies but is predicted by RNAfold^15^ to be present in the population of equilibrium structures (see **Extended Data Fig. 7**); (2) a short-range connection that cannot coexist in the same structure as the connection in (1), and therefore suggests multiple structures are present in the population; and (3) an isolated connection that is not part of a contiguous cluster and might therefore reflect a yet unreported pseudoknot. The high credibility of this connection (signal/uncertainty = 14.92, **Fig. 2**) and its reproducibility in replicate experiments (**Supplementary Fig. S19**) indicate that it does not arise from measurement noise. The probe sites are within the central region of the sequence that forms the consensus T-domain, and the patch site is 250 nt downstream of the central region. For molecules in the population that adopt the T-domain, a connection between these sites would form a long-range pseudoknot. While it is possible that the T-domain and the isolated connection exist only in separate molecules, their high prevalence suggests they likely overlap in the same molecule some of the time. **B.** Prevalence maps for Ψ-containing RNA molecules inferred from measurements with 12-nt probes. **C.** The presence of partially complementary sequences within the regions highlighted in boxes (1-3) suggest specific connections that may be present in the population.

## Supplementary Information

Supplementary Information includes supplementary figures, supplementary tables, and supplementary methods.

