## Supplementary Information for "Measuring intramolecular connectivity in long RNA molecules using two-dimensional DNA patch-probe arrays"

**This PDF file includes:**

Supplementary Figures
Supplementary Tables
Supplementary Methods

### Supplementary Figures

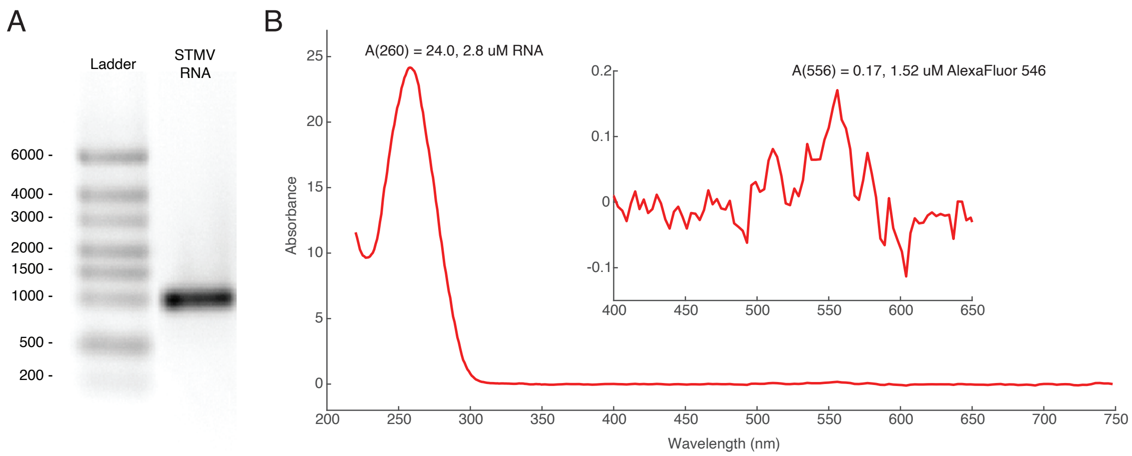


**Figure S1. Integrity, purity, and labeling density of STMV RNA transcripts. A**. In vitro transcribed STMV RNA runs as a single band in native agarose gel, indicating the RNA transcripts are not degraded. An RNA ladder with bands labeled by their length in nts is shown in the left lane. RNA bands are visualized by staining with ethidium bromide. **B**. UV-Vis absorbance spectrum of Alexa-Fluor-546-UTP-labeled STMV RNA shows an average of (1.5/2.8 =) 0.54 dye molecules per RNA molecule. For RNA, we take the extinction coefficient at 260 nm to be 25 $\mu L\ \mu g^{-1}\ cm^{-1}$. Taking the MW of STMV RNA to be 341,000 g/mol, we compute the molar extinction coefficient of STMV RNA to be 8.54 $\mu M^{-1}\ cm^{-1}$ at 260 nm. For Alexa Fluor 546-UTP, we take the molar extinction coefficient at 556 nm to be 0.11 $\mu M^{-1}\ cm^{-1}$.

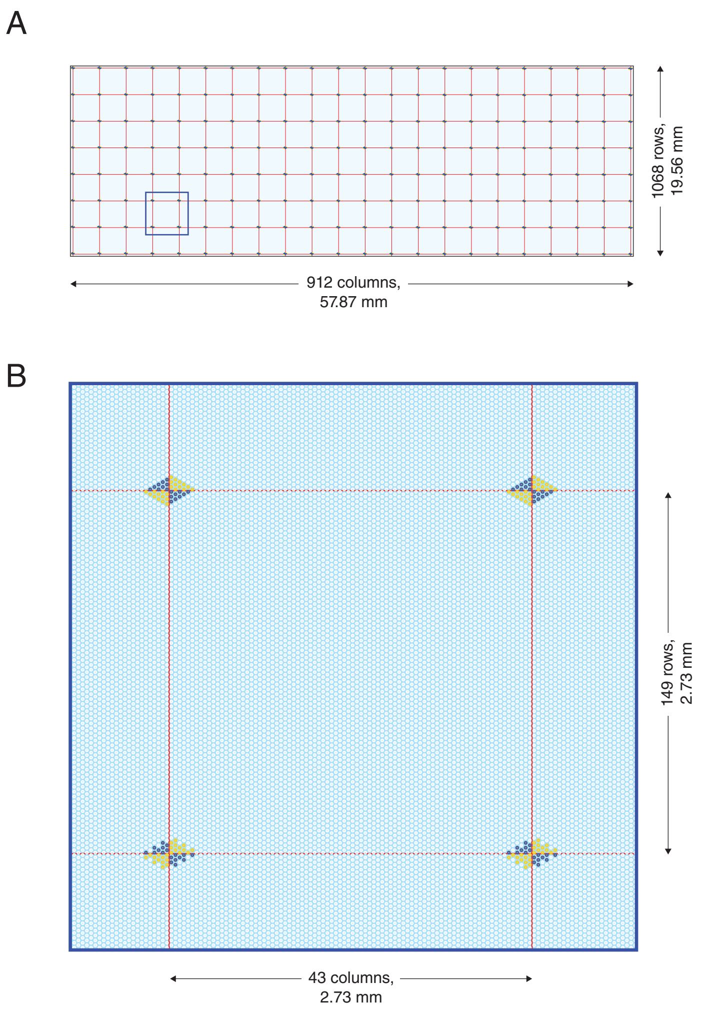


**Figure S2. Design of the microarray**. **A**. Layout and dimensions of the Agilent 3G microarray. The microarray contains spots arranged in a hexagonal grid. There are 1068 rows and 912 columns of spots, for a total of 974,016 individual spots. **B**. Layout and dimensions of a subarray. We design each subarray to contain 149 rows and 43 columns, as described in Methods. The upper left and bottom right corners contain a triangular pattern of control spots with poly-T sequences (dark blue spots), and the upper right and bottom left corners contain a triangular pattern of control spots with poly-A sequences (yellow spots). These spots help us identify the boundaries of each subarray.

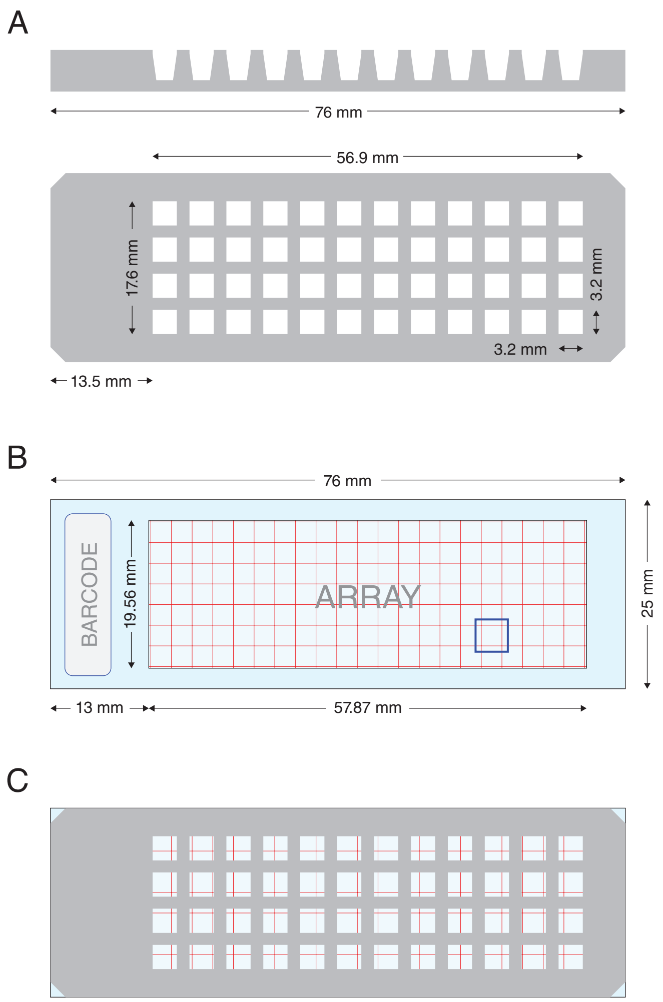


**Figure S3. Design of the gasket**. **A**. Dimensions of the PDMS gasket. **B**. Dimensions of the Agilent microarray glass slide. **C**. Overlay of the gasket and microarray.

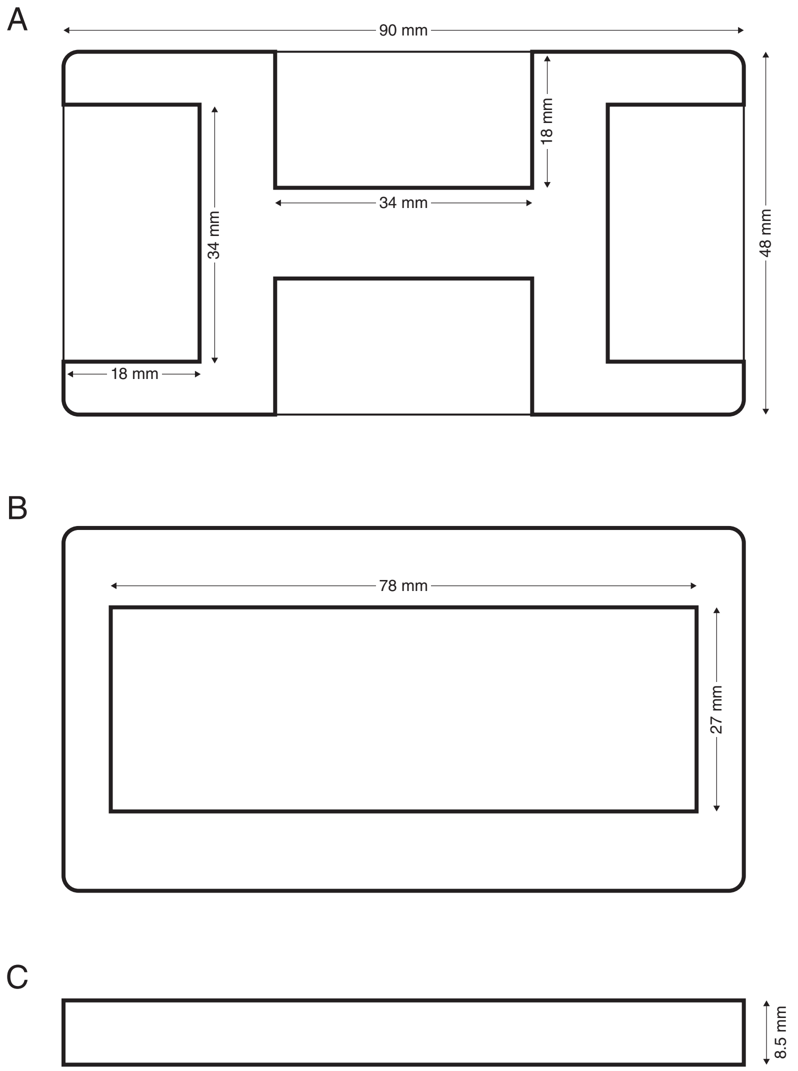


**Figure S4. Design of the clamp**. **A**. Top view. **B**. Bottom view. **C**. Side view.

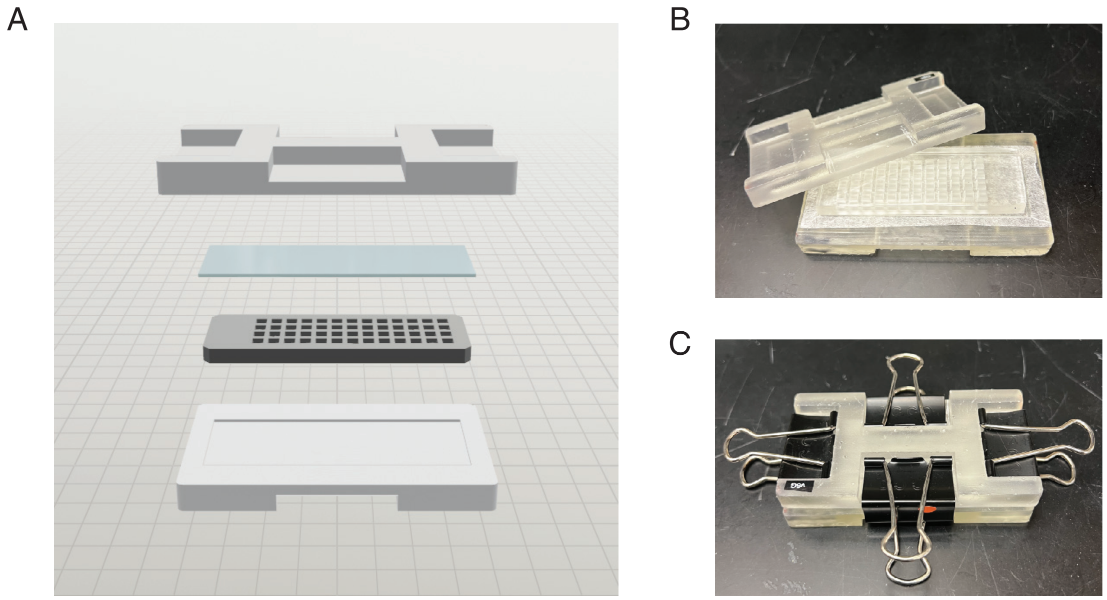


**Figure S5. Hybridization chamber**. **A**. Assembly diagram of the hybridization chamber shows the top clamp, gasket, and microarray slide, and bottom clamp. The gasket slide is placed face up and samples are pipetted into each well. Then we place the microarray slide DNA-side down on top of the gasket. The gasket and microarray are clamped together with binder clips. **B**. Photo of the unassembled chamber. **C**. Photo of the fully assembled chamber. We remove the handles of the binder clips before placing the chamber in the centrifuge.

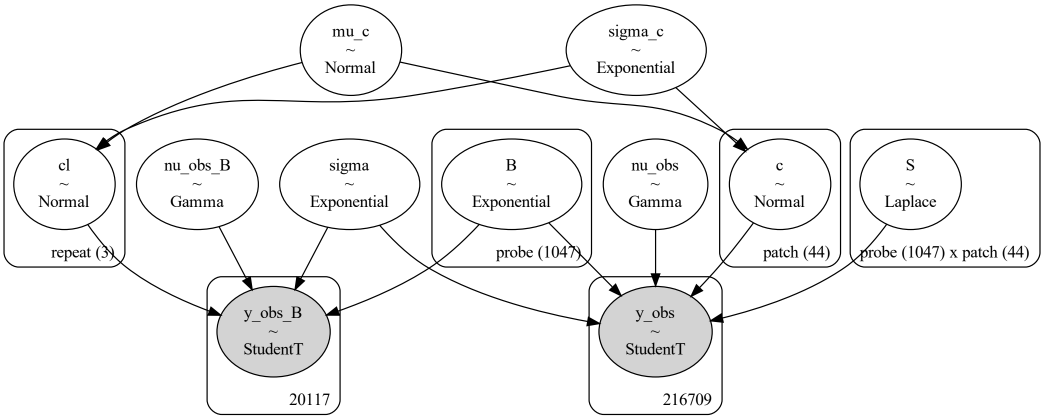


**Figure S6. Graphical depiction of Bayesian signal model** for the STMV RNA, 12-nt probe dataset. Shaded nodes at bottom correspond to the likelihoods. Numbers at the bottom of the likelihood nodes indicate the number of data points for the background measurements and patch-probe measurements. Other nodes indicate priors and, for arrays of parameters, their dimensions. Choices of priors and likelihood functions are discussed in section 2. Graphical depictions of the signal model for other datasets are the same, except that the number of data points differs from dataset to dataset, and the number of probes differs for datasets obtained with 12-nt probes and those obtained with 24-nt probes. Legend:

| | | |
|---|---|---|
| `mu_c` | $\mu_c$ | hyperparameter for mean of scalings |
| `sigma_c` | $\sigma_c$ | hyperparameter for standard deviation of scalings |
| `cl` | $c_l$ | scaling for background repeat measurement $l$ |
| `nu_obs_B` | $\nu_B$ | Student $T$ degrees of freedom for background measurements |
| `sigma` | $\sigma$ | uncertainty in measurements |
| `B` | $\hat{B}_i$ | inferred background for probe $i$ |
| `nu_obs` | $\nu_I$ | Student $T$ degrees of freedom for patch-probe measurements |
| `c` | $c_j$ | scaling for patch $j$ |
| `S` | $S_{ij}$ | signal for probe $i$ and patch $j$ |
| `y_obs_B` | $B_{ilm}$ | measured value for background for probe $i$, repeat $l$, spot $m$ |
| `y_obs` | $I_{ijk}$ | measured value for integrated fluorescence for probe $i$, patch $j$, spot $k$. |

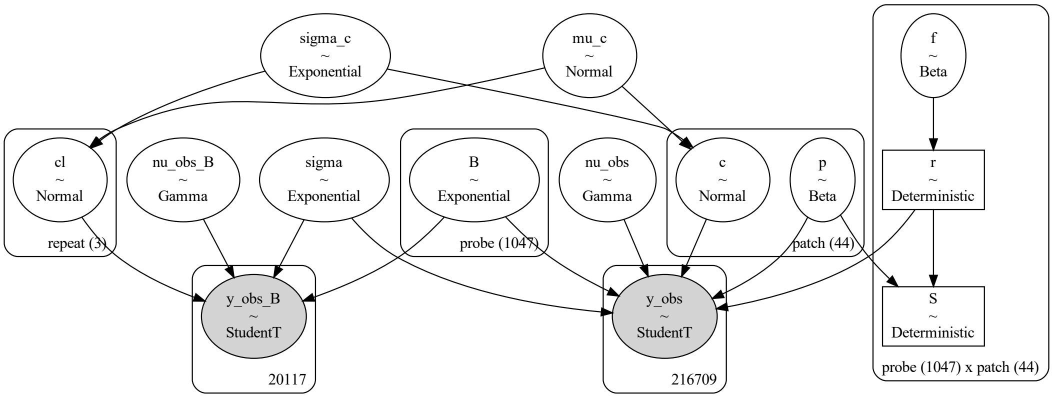


**Figure S7. Graphical depiction of Bayesian prevalence model** for the STMV RNA, 12-nt probe dataset. Description of nodes is the same as that described in Figure S6, and notation is the same with the addition of the following parameters:

| | | |
|---|---|---|
| `f` | $\underline{f_{i,j}}$ | prevalence for probe $i$ and patch $j$ |
| `r` | $\underline{r_{i,j}}$ | ratio for probe $i$ and patch $j$ |

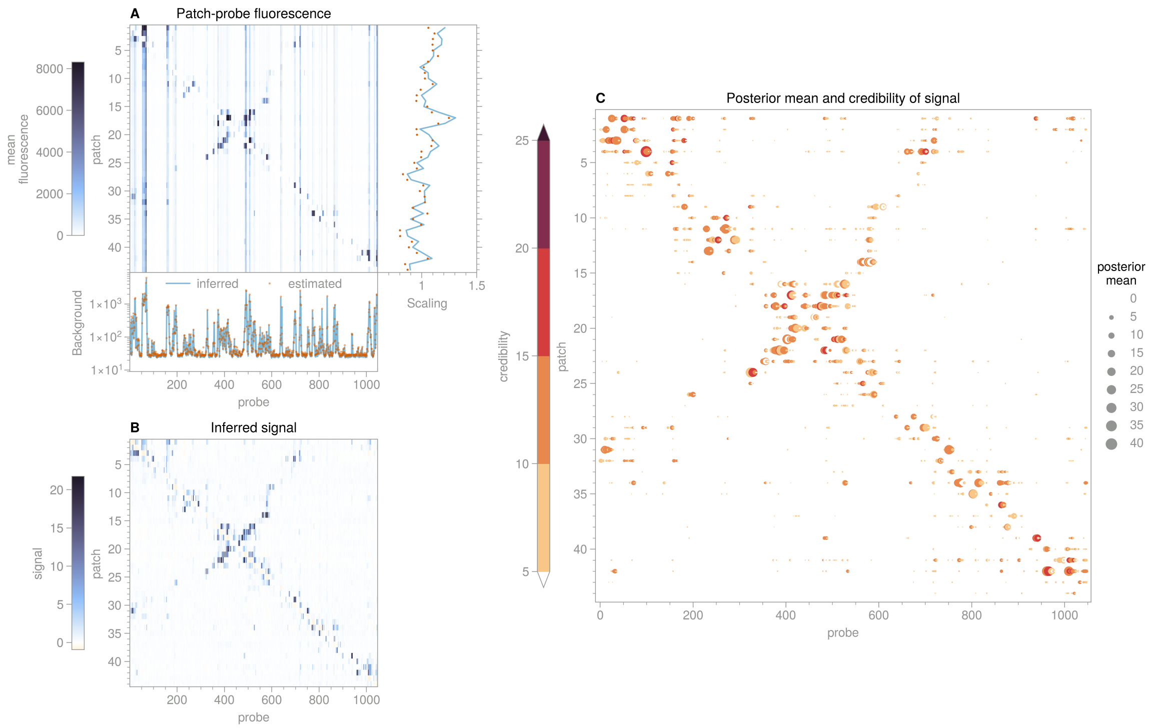


**Figure S8. Signal for STMV RNA, 12-nt probes**. **A**. Heatmap of patch-probe fluorescence data from microarray, averaged over spots for each (patch, probe) combination. Subplot at bottom shows the background estimated from a mean of the measurements taken in the four wells containing unpatched RNA, and the background inferred from the data using the Bayesian approach described in the Supplementary Methods. Subplot at right shows the median values of the fluorescence across all probes for each patch (dots) and the scaling values inferred from our Bayesian approach (blue line). **B**. Heatmap of signal inferred from the data in panel A using the Bayesian approach. In both panels A and B, the upper limit of the colormap corresponds to the 99.9th percentile of the values in the heatmap. **C**. Scatter plot showing the posterior mean (magnitude represented by size of symbols), and credibility (represented by color) of the inferred signals for each patch and probe combination. Only signal values with a credibility greater than 5 are shown.

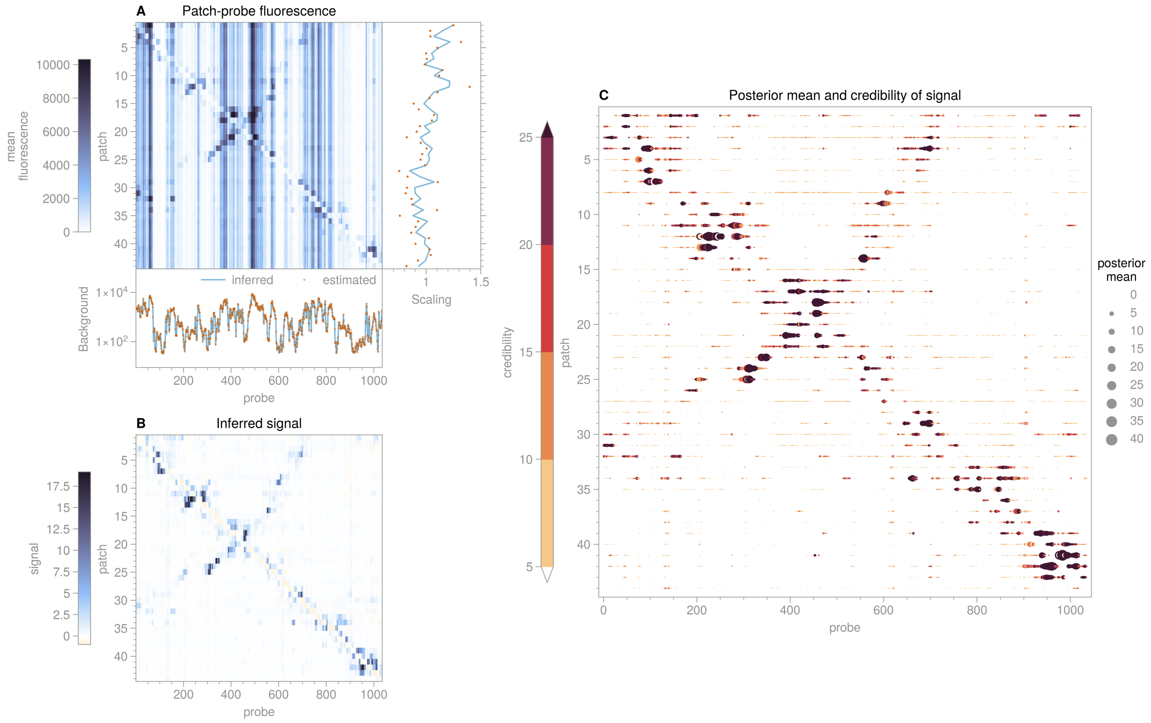


**Figure S9. Signal for STMV RNA, 24-nt probes**. See Figure S8 for detailed description of panels.

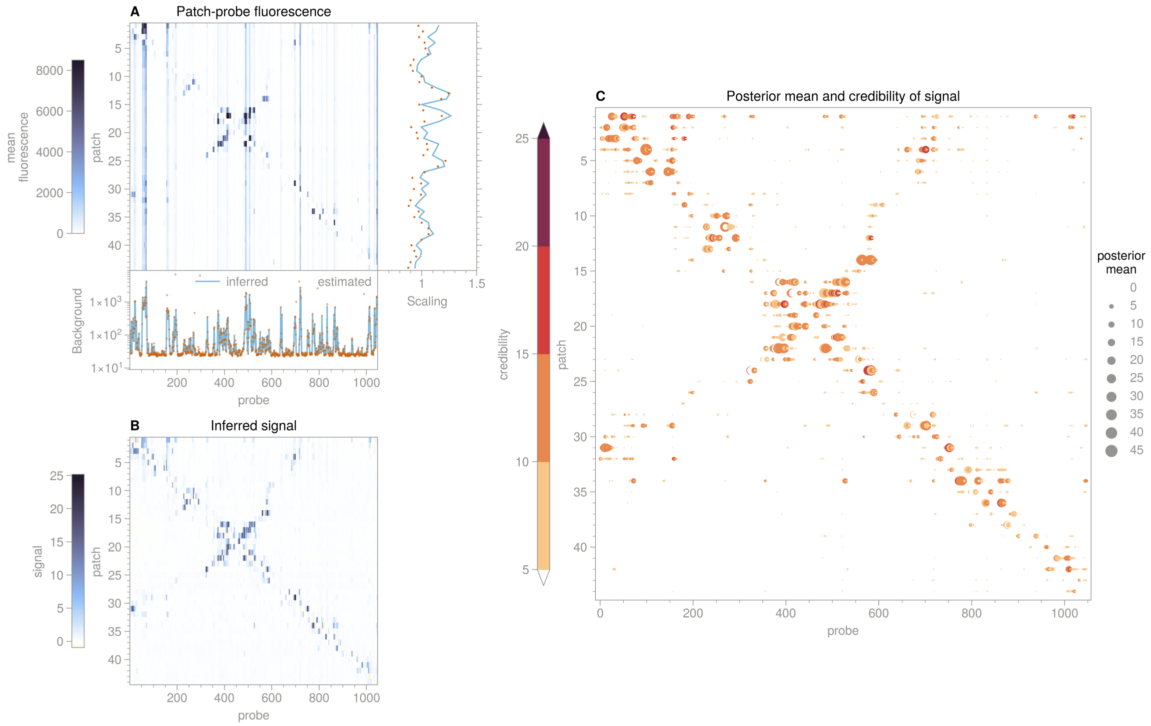


**Figure S10. Signal for STMV RNA, 12-nt probes, where patches are not annealed with the RNA**. See Figure S8 for detailed description of panels.

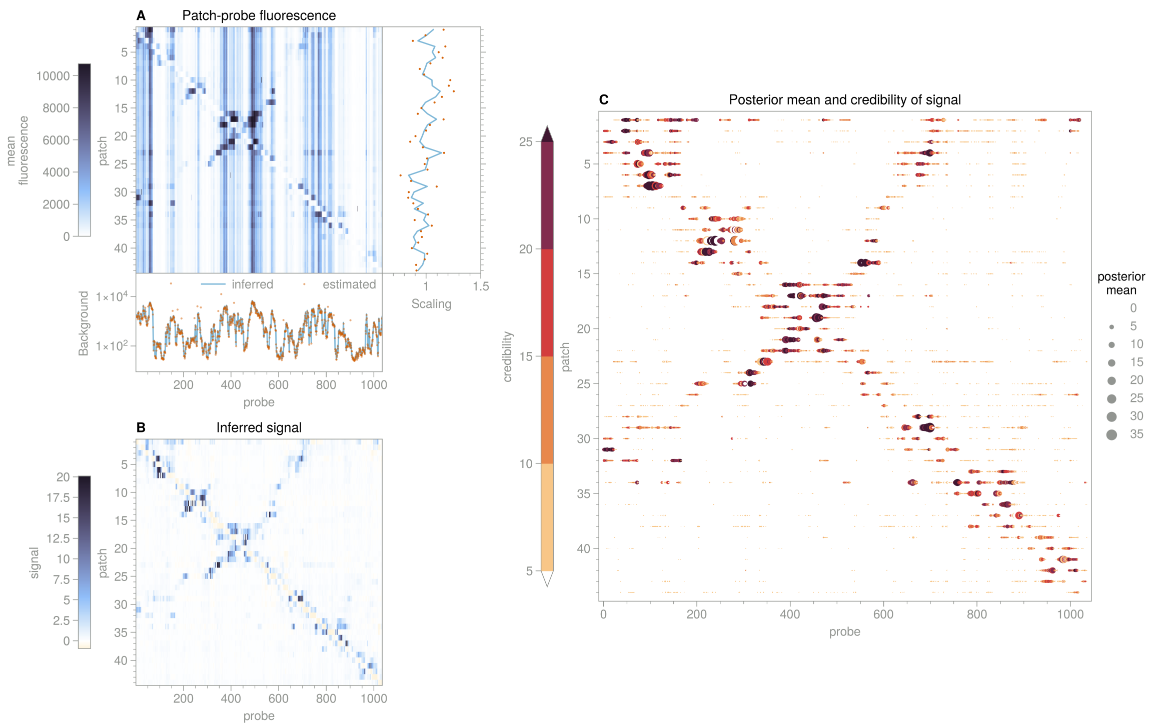


**Figure S11. Signal for STMV RNA, 24-nt probes, where patches are not annealed with the RNA**. See Figure S8 for detailed description of panels.

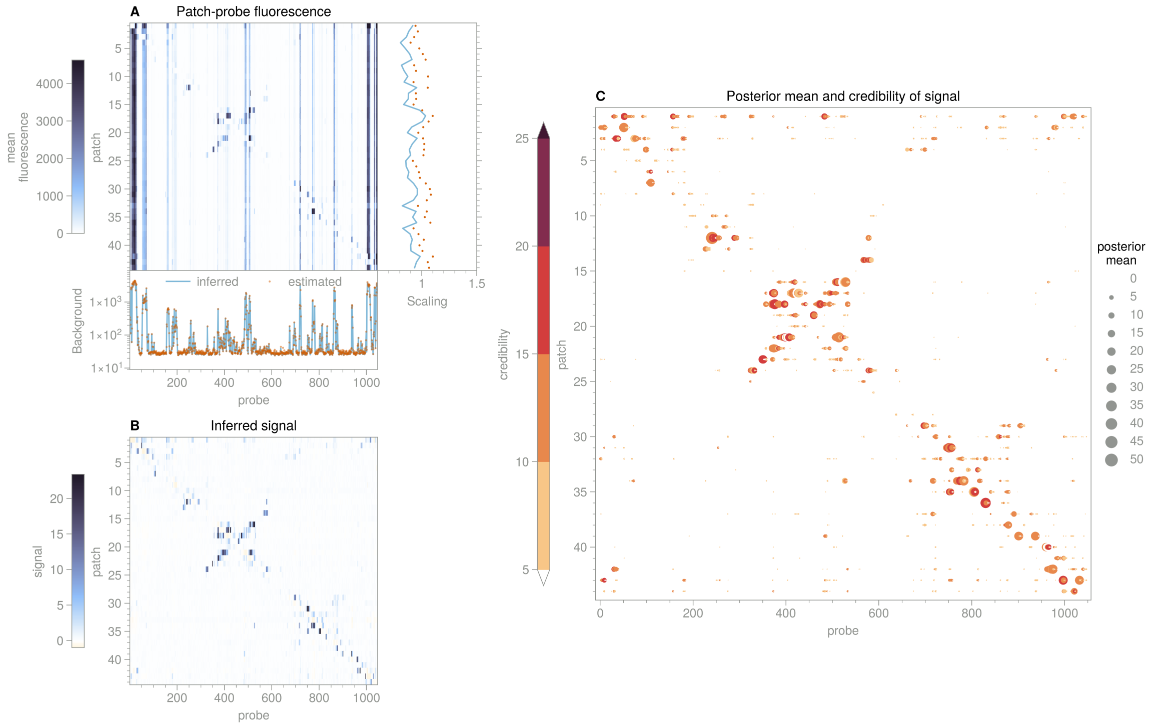


**Figure S12. Signal for STMV RNA with pseudouridine substitution and 12-nt probes.** See Figure S8 for detailed description of panels.

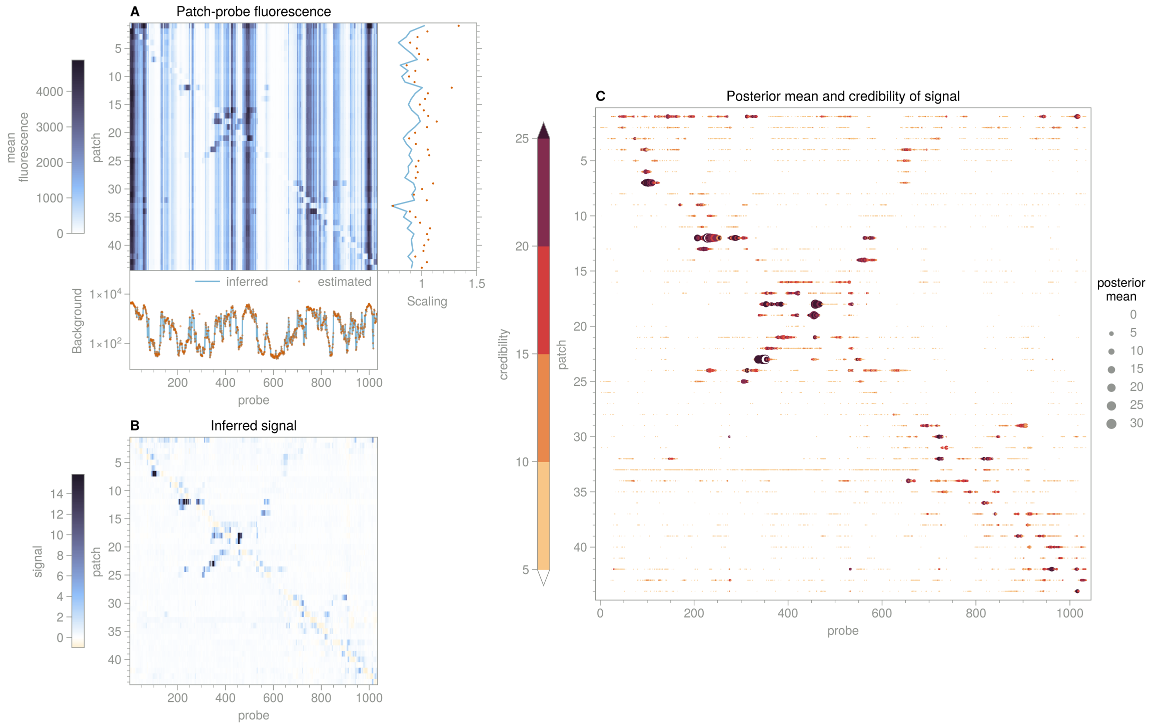


**Figure S13. Signal for STMV RNA with pseudouridine substitution and 24-nt probes.** See Figure S8 for detailed description of panels.

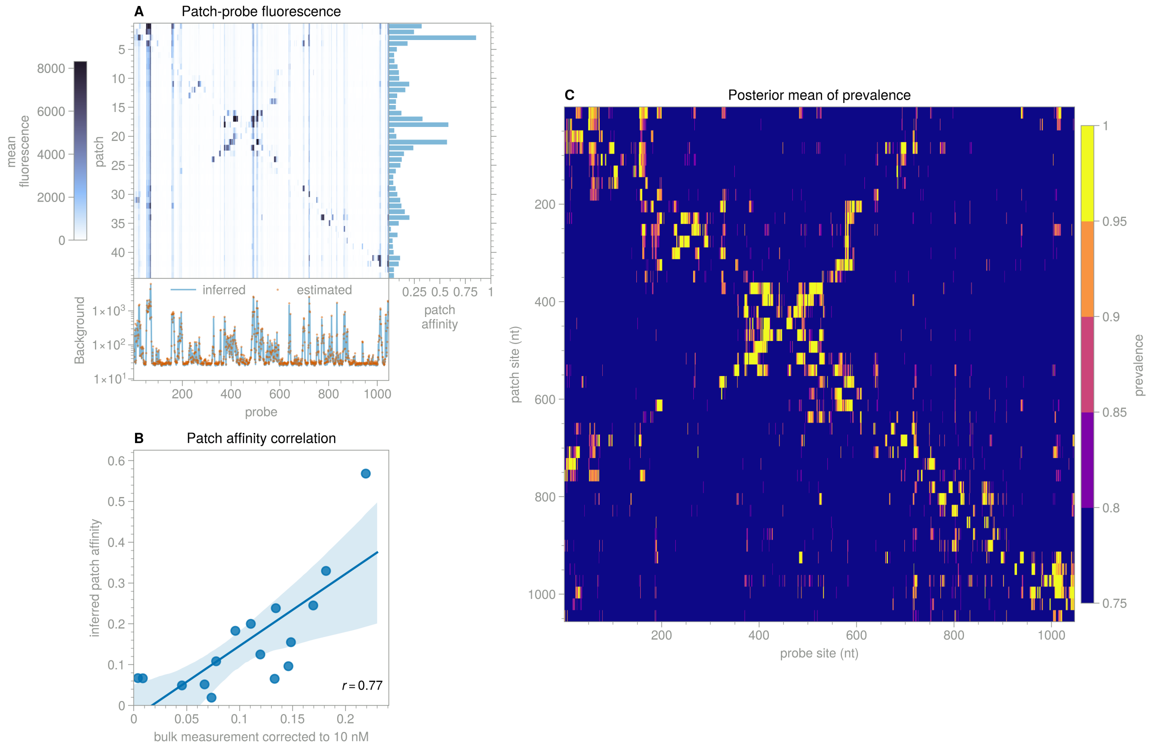


**Figure S14. Prevalence for STMV RNA, 12-nt probes**. **A**. Heatmap of patch-probe fluorescence data from microarray, averaged over spots for each (patch, probe) combination. The upper limit of the colormap corresponds to the 99.9th percentile of the values in the heatmap. Subplot at bottom shows the background estimated from a mean of the measurements taken in the four wells containing unpatched RNA, and the background inferred from the data using the Bayesian approach described in the Supplementary Methods. Subplot at right shows the inferred values (posterior mean) of the patch affinities. **B**. Correlation of patch affinities from the data in panel A with bulk measurements corrected to 10 nM. Pearson correlation coefficient is shown in the bottom right of the plot. Line is the best fit to a linear model, with 95% confidence interval shown by the blue region. **C**. Heatmap showing the posterior mean of the prevalence for each patch and probe combination.

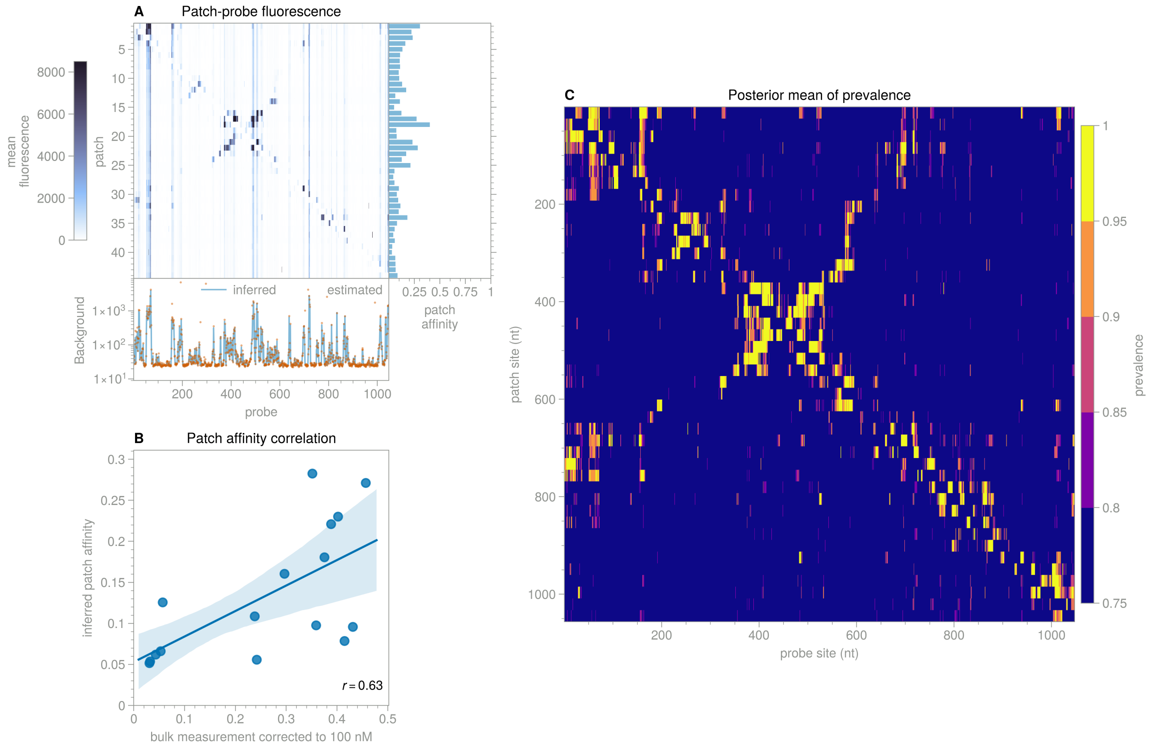


**Figure S15. Prevalence for STMV RNA, 12-nt probes, where patches are not annealed with the RNA**. See Figure S14 for detailed description of panels. Bulk patch affinities shown in panel B were corrected to 100 nM to match the patch concentation used in this protocol.

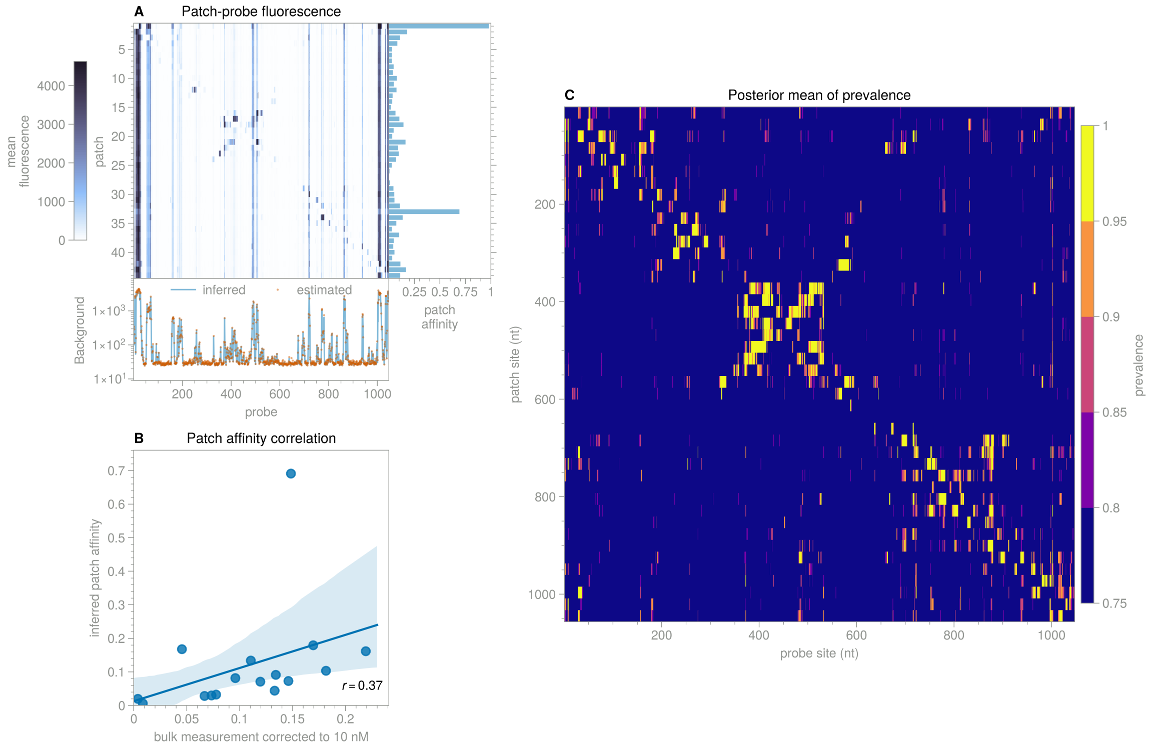


**Figure S16. Prevalence for STMV RNA with pseudouridine substitution and 12-nt probes.** See Figure S14 for detailed description of panels. The bulk measurements of patch affinity shown in panel B were carried out with unsubstituted RNA, and we do not expect to observe high correlation between these measurements and those inferred from microarray measurements with substituted RNA.

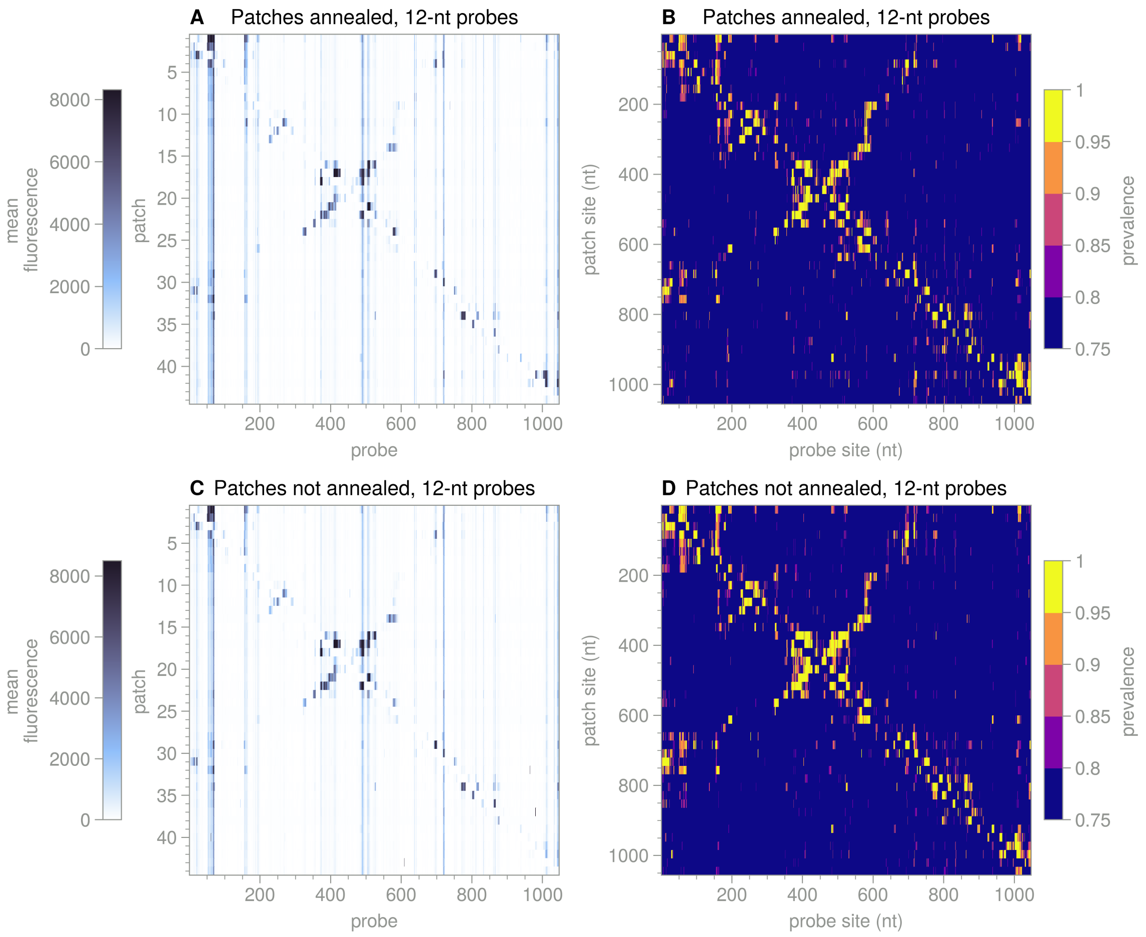


**Figure S17. Comparison of results for annealed and unannealed patches**. **A**. Heatmap of patch-probe fluorescence data for STMV RNA with 12-nt probes and patches that were annealed with RNA after mixing. Data is averaged over spots for each (patch, probe) combination. **B**. Heatmap showing the posterior mean of the inferred prevalence for each (patch, probe) combination in panel A. **C**, **D**. As in panels A and B, but for a microarray in which patches were not annealed with the RNA after mixing. In both panels A and C, the upper limit of the colormap corresponds to the 99.9th percentile of the values in the heatmap.

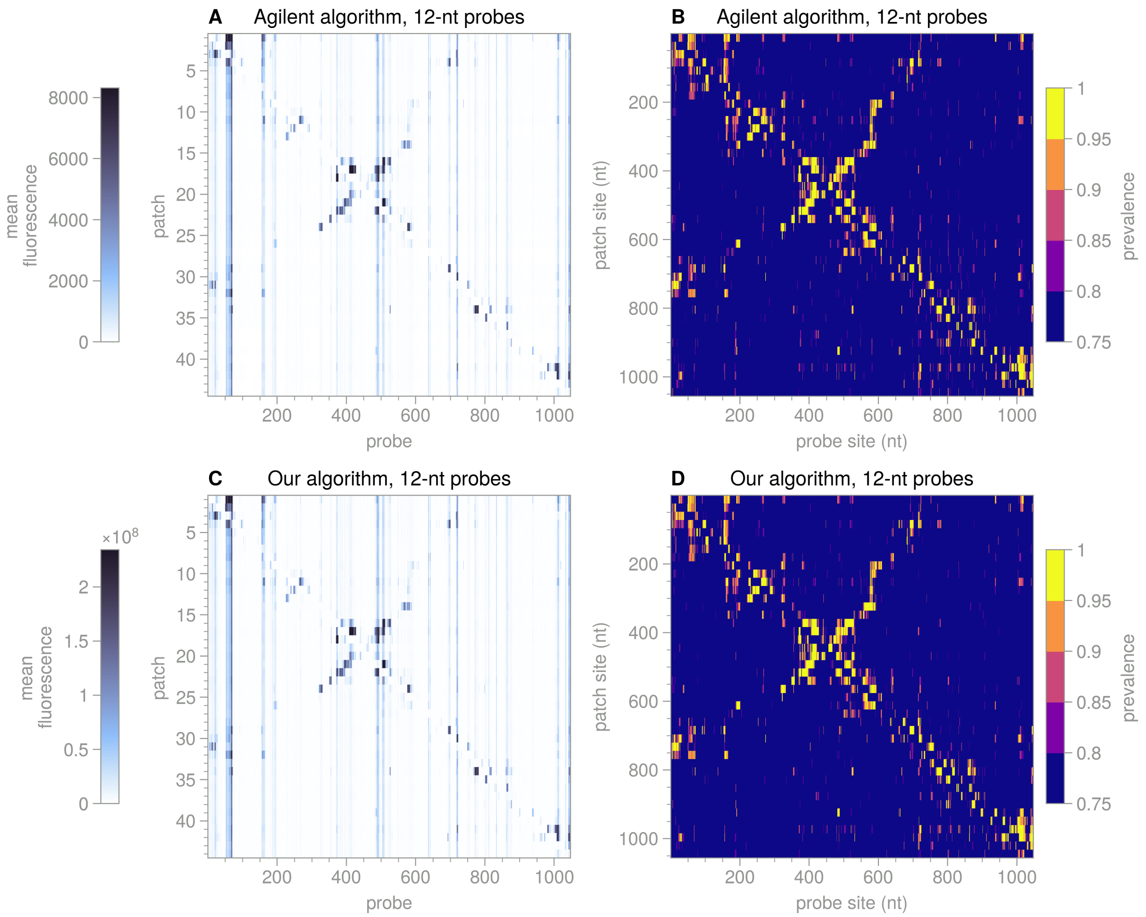


**Figure S18. Comparison of results from Agilent feature extraction and our own method**. **A**. Heatmap of patch-probe fluorescence data for STMV RNA with 12-nt probes, averaged over spots for each (patch, probe) combination. Spots and their integrated fluorescences are extracted using Agilent software. **B**. Heatmap showing the posterior mean of the inferred prevalence for each (patch, probe) combination in panel A. **C**, **D**. As in panels A and B, but for spots and integrated fluorescences extracted using our own software. In both panels A and C, the upper limit of the colormap corresponds to the 99.9th percentile of the values in the heatmap.

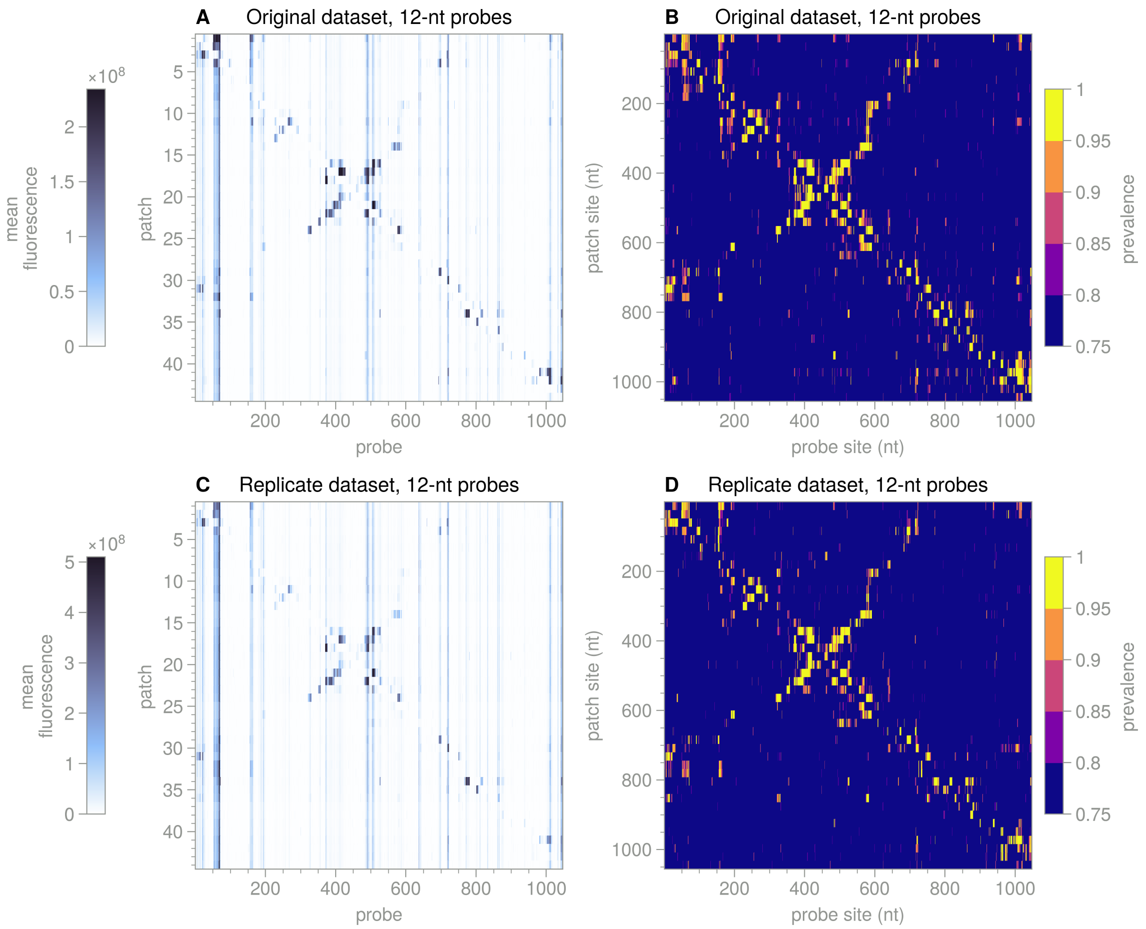


**Figure S19. Comparison of results from replicate patch-probe experiments.** **A**. Heatmap of patch-probe fluorescence data for STMV RNA with 12-nt probes, averaged over spots for each (patch, probe) combination. **B**. Heatmap showing the posterior mean of the inferred prevalence for each (patch, probe) combination in panel A. **C**, **D**. As in panels A and B, but for a replicate measurement taken on a different microarray with the same probe and patch sequences. In both panels A and C, the upper limit of the colormap corresponds to the 99.9th percentile of the values in the heatmap. Spots and their integrated fluorescences are extracted using our own software for both datasets.

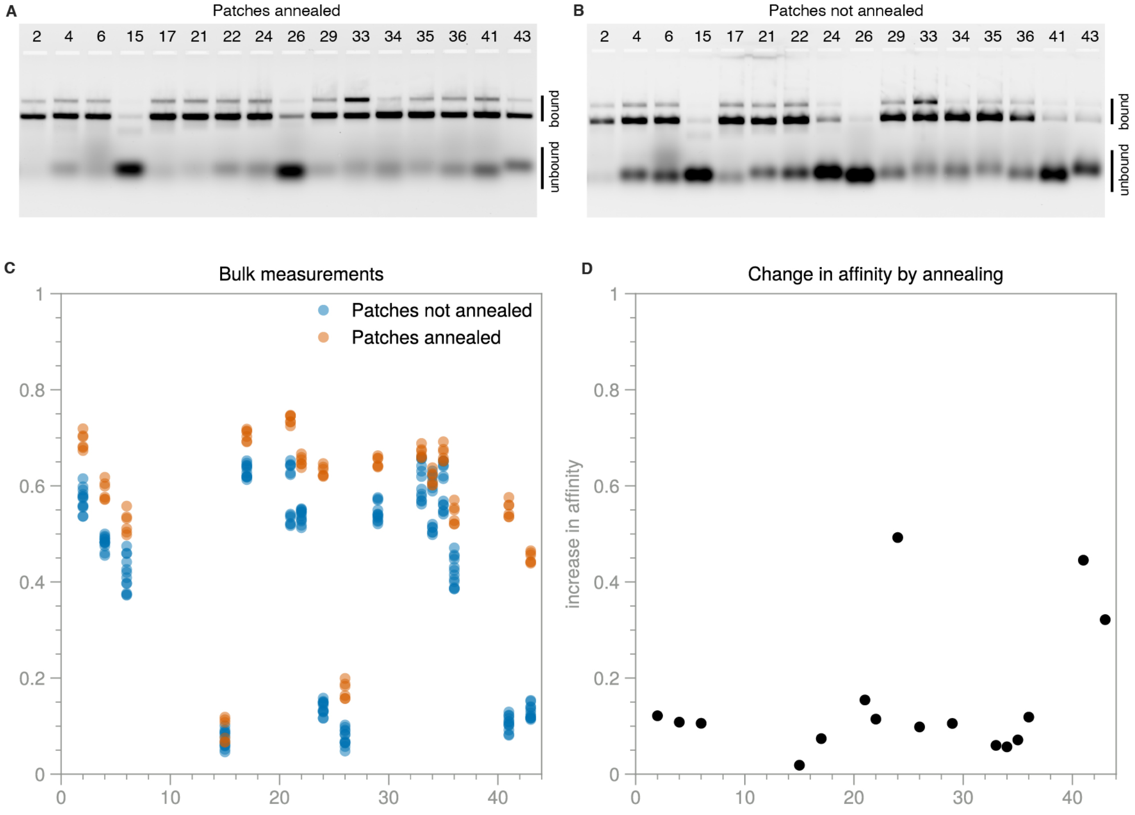


**Figure S20. Bulk measurements of patch affinities. A**. Native 1% agarose gel electrophoresis experiments in which fluorescent patches are annealed with unlabeled RNA. Bars to the right of the gel show the positions of bound (slower migrating) and unbound (faster migrating) patches. The population of bound patches appears as two sharp fluorescent bands: a faster-moving bright band corresponding to patches bound to monomers of RNA, and a slower-moving fainter band corresponding to patches bound to dimers of RNA. Dimers can form in these experiments owing to the high (300 nM) concentration of RNA used. The population of unbound patches appears as a single broad band whose mobility depends on the secondary structure of the patch. **B**. Gel electrophoresis experiments in which the fluorescent patches are not annealed with the RNA. **C**. Plot of binding affinities for each protocol and patch. Taking the ratio of bound fluorescence to total (bound plus unbound) fluorescence for each patch gives the binding affinity. These fluorescence values are measured by integrating the fluorescence along each lane of the gel. To perform the integration, a cutoff position between bound and unbound patches and a background level are manually chosen. To account for slight differences in these manually chosen values, the integration protocol is performed several times, and each replicate integration (3–6 replicates per patch) is shown. **D**. Change in affinity caused by annealing. Each point is the mean of the differences between the two sets of measurements shown in panel C for each patch.

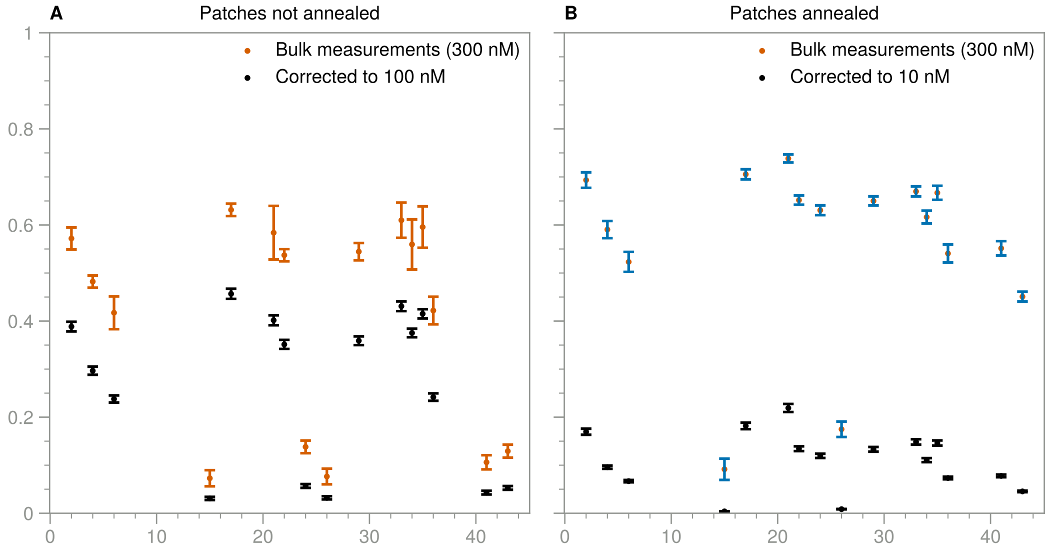


**Figure S21. Comparison of measured and corrected bulk patch affinities. A**. Bulk measurements of patch affinities (mean of replicates, with standard deviation shown by the error bars) and corrected values (mean of posterior, with standard deviation of posterior shown by the error bars) for unannealed patches. Affinities are corrected to 100 nM using the method described in section 3. **B**. As in panel A, but for bulk measurements of patches that are annealed with the RNA. Affinities are corrected to 10 nM.

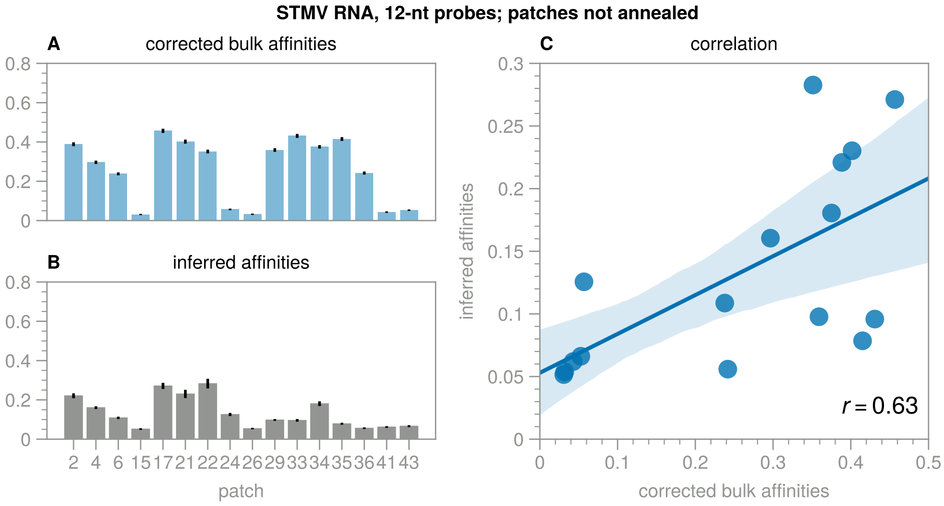


**Figure S22. Comparison of inferred and corrected bulk affinities for unannealed patches. A**. Bulk measurements of patch affinities, corrected to 100 nM using the method described in section 3. Error bars show standard deviation of posterior. **B**. Patch affinities inferred with prevalence model from the microarray data for 12-nt probes, unannealed patches. Bar heights show mean of posterior, and error bars show standard deviation of posterior. **C**. Correlation of inferred and corrected bulk affinities. Pearson correlation coefficient is shown in the bottom right of the plot. Line is the best fit to a linear model, with 95% confidence interval shown by the blue region.

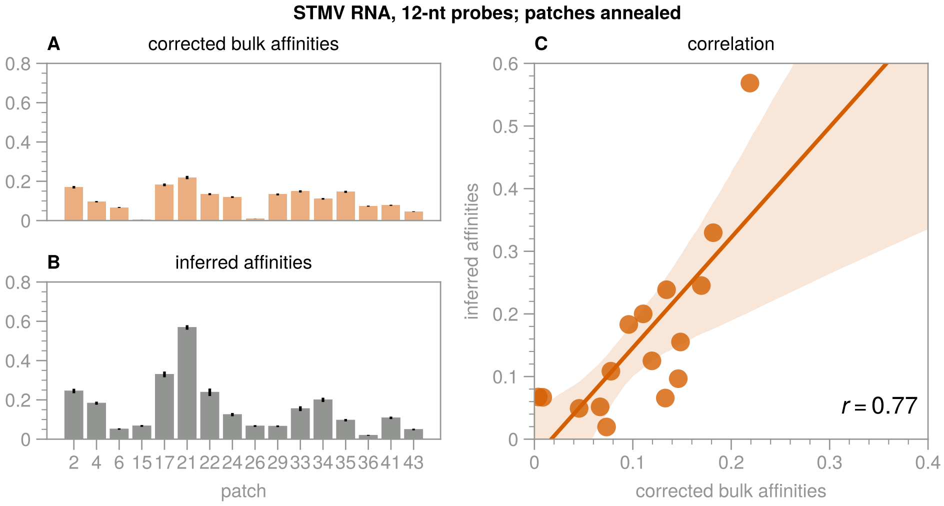


**Figure S23. Comparison of inferred and corrected bulk affinities for annealed patches. A**. As in Figure S22A but showing data for annealed patches corrected to 10 nM. **B**. As in Figure S22B, but showing results inferred from microarray data for annealed patches. **C**. As in Figure S22C, but showing correlation for annealed patches.

### Supplementary Tables

#### Results for hyperparameters of signal model

**Table S1.** Summary of marginalized posteriors of hyperparameters in the Bayesian signal model applied to STMV RNA with 12-nt probes. Hyperparameters are described in section 2. The HDI 3% and 97% values show the bounds of the highest density interval containing 94% of the posterior probability.

| | $\mu_c$ | $\sigma_c$ | $\sigma$ | $\nu_B$ | $\nu_I$ |
|---|---|---|---|---|---|
| posterior mean | 1.04 | 0.11 | 0.12 | 4.97 | 3.90 |
| posterior standard deviation | 0.02 | 0.01 | 0.00 | 0.14 | 0.04 |
| HDI 3% value | 1.01 | 0.08 | 0.12 | 4.69 | 3.82 |
| HDI 97% value | 1.07 | 0.13 | 0.12 | 5.21 | 3.98 |

**Table S2.** Summary of marginalized posteriors of hyperparameters in the Bayesian signal model applied to STMV RNA with 24-nt probes.

| | $\mu_c$ | $\sigma_c$ | $\sigma$ | $\nu_B$ | $\nu_I$ |
|---|---|---|---|---|---|
| posterior mean | 1.02 | 0.10 | 0.03 | 1.22 | 1.61 |
| posterior standard deviation | 0.01 | 0.01 | 0.00 | 0.01 | 0.01 |
| HDI 3% value | 1.00 | 0.08 | 0.03 | 1.19 | 1.60 |
| HDI 97% value | 1.05 | 0.12 | 0.03 | 1.25 | 1.63 |

**Table S3.** Summary of marginalized posteriors of hyperparameters in the Bayesian signal model applied to STMV RNA with 12-nt probes, patches not annealed.

| | $\mu_c$ | $\sigma_c$ | $\sigma$ | $\nu_B$ | $\nu_I$ |
|---|---|---|---|---|---|
| posterior mean | 1.05 | 0.09 | 0.12 | 4.90 | 4.46 |
| posterior standard deviation | 0.01 | 0.01 | 0.00 | 0.13 | 0.05 |
| HDI 3% value | 1.03 | 0.07 | 0.12 | 4.67 | 4.35 |
| HDI 97% value | 1.08 | 0.11 | 0.12 | 5.14 | 4.55 |

**Table S4.** Summary of marginalized posteriors of hyperparameters in the Bayesian signal model applied to STMV RNA with 24-nt probes, patches not annealed.

| | $\mu_c$ | $\sigma_c$ | $\sigma$ | $\nu_B$ | $\nu_I$ |
|---|---|---|---|---|---|
| posterior mean | 0.98 | 0.08 | 0.05 | 1.11 | 1.57 |
| posterior standard deviation | 0.01 | 0.01 | 0.00 | 0.01 | 0.01 |
| HDI 3% value | 0.96 | 0.06 | 0.05 | 1.09 | 1.55 |
| HDI 97% value | 1.00 | 0.10 | 0.05 | 1.14 | 1.59 |

**Table S5.** Summary of marginalized posteriors of hyperparameters in the Bayesian signal model applied to STMV RNA with 12-nt probes, where RNA is pseudouridine substituted.

| | $\mu_c$ | $\sigma_c$ | $\sigma$ | $\nu_B$ | $\nu_I$ |
|---|---|---|---|---|---|
| posterior mean | 0.91 | 0.06 | 0.11 | 6.31 | 5.42 |
| posterior standard deviation | 0.01 | 0.01 | 0.00 | 0.21 | 0.08 |
| HDI 3% value | 0.89 | 0.05 | 0.11 | 5.93 | 5.26 |
| HDI 97% value | 0.92 | 0.07 | 0.11 | 6.72 | 5.56 |

**Table S6.** Summary of marginalized posteriors of hyperparameters in the Bayesian signal model applied to STMV RNA with 24-nt probes, where RNA is pseudouridine substituted.

| | $\mu_c$ | $\sigma_c$ | $\sigma$ | $\nu_B$ | $\nu_I$ |
|---|---|---|---|---|---|
| posterior mean | 0.90 | 0.06 | 0.05 | 2.56 | 2.20 |
| posterior standard deviation | 0.01 | 0.01 | 0.00 | 0.05 | 0.01 |
| HDI 3% value | 0.88 | 0.05 | 0.05 | 2.47 | 2.18 |
| HDI 97% value | 0.92 | 0.07 | 0.05 | 2.64 | 2.23 |

### Results for hyperparameters of prevalence model

**Table S7.** Summary of marginalized posteriors of hyperparameters in the Bayesian prevalence model applied to STMV RNA with 12-nt probes. Hyperparameters are described in section 2. The HDI 3% and 97% values show the bounds of the highest density interval containing 94% of the posterior probability.

| | $\mu_c$ | $\sigma_c$ | $\sigma$ | $\nu_B$ | $\nu_I$ |
|---|---|---|---|---|---|
| posterior mean | 1.02 | 0.09 | 0.12 | 5.03 | 3.36 |
| posterior standard deviation | 0.01 | 0.01 | 0.00 | 0.15 | 0.03 |
| HDI 3% value | 0.99 | 0.07 | 0.12 | 4.74 | 3.30 |
| HDI 97% value | 1.04 | 0.10 | 0.12 | 5.29 | 3.41 |

**Table S8.** Summary of marginalized posteriors of hyperparameters in the Bayesian prevalence model applied to STMV RNA with 12-nt probes, patches not annealed.

| | $\mu_c$ | $\sigma_c$ | $\sigma$ | $\nu_B$ | $\nu_I$ |
|---|---|---|---|---|---|
| posterior mean | 1.04 | 0.09 | 0.12 | 4.29 | 3.30 |
| posterior standard deviation | 0.01 | 0.01 | 0.00 | 0.10 | 0.02 |
| HDI 3% value | 1.02 | 0.07 | 0.12 | 4.09 | 3.26 |
| HDI 97% value | 1.06 | 0.11 | 0.12 | 4.48 | 3.35 |

**Table S9.** Summary of marginalized posteriors of hyperparameters in the Bayesian prevalence model applied to STMV RNA with 12-nt probes, where RNA is pseudouridine substituted.

| | $\mu_c$ | $\sigma_c$ | $\sigma$ | $\nu_B$ | $\nu_I$ |
|---|---|---|---|---|---|
| posterior mean | 0.92 | 0.09 | 0.11 | 5.00 | 3.45 |
| posterior standard deviation | 0.01 | 0.01 | 0.00 | 0.19 | 0.03 |
| HDI 3% value | 0.90 | 0.08 | 0.11 | 4.65 | 3.40 |
| HDI 97% value | 0.94 | 0.11 | 0.11 | 5.33 | 3.51 |

### Supplementary Methods

Here we describe the methods used to infer the signals and prevalences from the data, and to test and validate these inferences. First, we briefly summarize the experimental procedure to highlight the details salient to the inference procedure.

As discussed in the online methods, the microarray consists of spots, each of which contains multiple copies of a probe with a specific sequence. The array contains spots with 12-nt-long and 24-nt-long probes. For each probe sequence, there are many spots that contain that probe and which are dispersed throughout the microarray. The microarray is divided into wells by a gasket. Each well contains many spots, including typically 3–12 spots for each probe sequence. To each well we add fluorescent RNA, either patched or unpatched. There are 44 wells to which we add patched RNA, one for each patch sequence. There are four wells to which we add unpatched RNA. After waiting a period of time for the RNA to bind to the probes, we rinse out all the wells and measure the fluorescence of all the spots that are not occluded by the gasket. The fluorescence is related to the amount of the RNA that remains near each spot, and which we assume to be bound to the probes.

Although we set our pipette to dispense the same volume of RNA (patched or unpatched) into each well, we expect that the actual amount dispensed can vary by about 10% or so, the typical scale of "pipetting noise." The four wells in which we measure the binding to unpatched RNA constitute repeated measurements which we can use to estimate the pipetting noise. The pipetting noise will also affect comparisons of binding between different probes (wells).

#### 1 Data and metadata

##### 1.1 Processing

The Agilent instrument measures the fluorescence at a large number of points (pixels) on the microarray. Each spot covers about 100 pixels. We use either Agilent software or our own software routine, written in Matlab, to extract "features" from the microarrays, which consist of the integrated fluorescence for each spot along with the probe number and well associated with each spot. The resulting data are stored as comma-separated value files, one file for each well and probe size. Within each file, each line corresponds to measurements of fluorescence of each spot corresponding to a particular probe, starting at probe 1 and ending at the last probe. The number of measurements per probe can vary from line to line, because the number of spots within each well varies for each probe.

##### 1.2 Data structures

We use Python to read the data into `xarray Dataset` objects that contain both metadata (such as probe size, patch size, and feature extraction software) and data for each microarray. We use a separate `Dataset` for each probe size, so that data on a single microarray is split up into two `Datasets`, one corresponding to 12-base-long probes and one to 24-base-long probes. We read in all the datasets at once and store them in a dictionary, keyed by a short-form identifier for each dataset.

Within each `Dataset`, we separate the measurements into patch-probe measurements and background measurements (which correspond to wells containing unpatched RNA). The patch-probe measurements are stored in `xarray DataArray` objects which contain integrated fluorescence measurements for each spot, patch, and probe. The number of values in the `spot` coordinate corresponds to the maximum number of spots for any patch-probe combination. We use `NaN` values to pad the data.

We also use a `DataArray` to store measurements on unpatched RNA. The integrated fluorescence measurements are stored for each spot, probe, and "repeat," where each repeat corresponds to a background measurement made in a different well.

The Agilent feature extraction software also returns "errors," corresponding to uncertainties for each integrated fluorescence measurement. If available, we store these uncertainties as a separate `DataArray` in the same `Dataset`. This `DataArray` has the same structure as that used to store the patch-probe data. However, because nearly all of these reported uncertainties are equal to 10% of the measurement, we do not use these values in our analysis. Instead, as described below, we fit for the uncertainty.

### 2 Data analysis

We use a Bayesian approach to analyze the data. First, we aim to infer how much the measured fluorescence changes when we patch the RNA, compared to when the RNA is added to the microarray without a patch. The results of this analysis are the signal values, described in the main text. Second, we aim to determine the prevalence of each connection from the data by inferring and correcting for the patch binding affinities, as discussed in the main text. In both cases, we use Markov-chain Monte Carlo (MCMC) sampling to calculate the posterior probability density of the quantities of interest. The procedure and assumptions are described in detail below.

#### 2.1 Notation

Let $i = 1 : n_{\text{probe}}$ be an index over probe sites ($n_{\text{probe}}$ is the number of probe sites). Let $j = 1 : n_{\text{patch}}$ be an index over patches ($n_{\text{patch}}$ is the number of patches, which is equivalent to the number of wells on the microarray into which patches were added). Let $l = 1 : n_{\text{backgrounds}}$ be an index over measurements of the unpatched RNA ($n_{\text{backgrounds}}$) is the number of wells in which the unpatched RNA binding is measured}.

There are two types of data. We measure the integrated fluorescence $I_{ijk}$ for probe $i$ and patch $j$, where $k = 1 : n_{ij}$ is an index that runs over all microarray spots containing probe $i$ and subjected to RNA with patch $j$ (there are $n_{ij}$ spots for a given probe and patch). We also have the unpatched RNA, or *background*, measurements $B_{ilm}$, where $l = 1 : n_i$ is an index that runs over the microarray spots for probe $i$ in repeat measurement $m$ (there are $n_i$ spots for a given probe and repeat). We let $\mathbf{D}$ represent all the data, that is, all the $I_{ijk}$ and $B_{ilm}$.

We also define the following chemical species:

- $R_{i,j}$: RNA molecule with sites $i$ and $j$ unconnected
- $R_{\underline{i,j}}$: RNA molecule with sites $i$ and $j$ connected
- $R$: naked RNA molecule, irrespective of connections
- $R_{\bar{\imath}}$: RNA molecule with probe attached to site $i$
- $R_{*,\bar{\jmath}}$: RNA molecule with patch attached to site $j$
- $R_{\bar{\imath},\bar{\jmath}}$: RNA molecule with probe attached to site $i$ and patch attached to site $j$

We use square bracket notation to denote the concentrations of these species; for example, $[R]$ is the concentration of RNA molecules.

#### 2.2 Signal

We first seek to calculate the posterior probability density of the *signal* in the data, which is related to how much the integrated fluorescence changes when we patch the RNA compared to when the RNA is unpatched. Let $\hat{B}_i$ be the true background for probe $i$ and for a given, reference amount of RNA. Let $\hat{I}_{ij}$ be the true integrated fluorescence for probe $i$ at the same reference RNA amount, where the RNA is patched with patch $j$. Then, the signal $S_{ij}$ is defined as the difference between the true integrated fluorescence for a given probe and patch and the true background for that probe, divided by the background:

$$S_{ij} = \frac{\hat{I}_{ij} - \hat{B}_i}{\hat{B}_i}. \tag{1}$$

The posterior on $S_{ij}$, $p(S_{ij} \mid \mathbf{D})$, is a quantitative measure of how much the binding of the patch has affected the binding of the probe. We estimate $S_{ij}$ from the mean of $p(S_{ij})$, and we calculate the uncertainty on this estimate from the width of the posterior, as characterized by its standard deviation.

We characterize the significance of each signal value through the ratio of the posterior mean to the posterior standard deviation, a metric that we call the *credibility*. The credibility is a nondimensional metric that describes how many standard deviations the signal differs from zero. The credibility is not necessarily related to the signal magnitude: even small signals may be significant if their posterior is narrow compared to their mean value. We assume that statistically significant signals correspond to situations in which the patch disrupts a connection and leads to an increase in probe binding, thereby providing evidence for a connection in the RNA molecule between the binding site for probe $i$ and the binding site for patch $j$.

#### 2.3 Prevalence

We also aim to calculate the prevalence of each individual connection in the population of RNA molecules. To do this, we must infer the affinities of both the probes and patches of the RNA molecules. The probe affinities are characterized by the true background values $\hat{B}_i$, as discussed above. The patch affinities are characterized by parameters $p_j$, the fraction of the RNA molecules to which patch $j$ is attached.

We use a linear model to infer the prevalences. The simplest linear model we can write for the integrated fluorescence of a microarray spot is

$$\hat{I}_{ij} = \hat{B}_i \left(1 + p_j r_{\underline{i,j}}\right) \tag{2}$$

where

$$r_{\underline{i,j}} = \frac{\left[R_{\underline{i,j}}\right]_0}{\left[R_{i,j}\right]_0} \tag{3}$$

is the ratio of bulk concentrations of RNA molecules with a connection between sites $i$ and $j$ (the "0" subscript denoting that this is the ratio before we add patches or probes). As above, $\hat{B}_i$ is the true background for probe $i$.

The intuitive explanation for (2) is that patching opens up molecules that had a connection between sites $i$ and $j$, thereby making site $i$ more accessible to the probe. The amount by which patching increases the spot intensity is proportional to the fraction of molecules that were patched.

The prevalence is then defined as

$$\begin{aligned} f_{\underline{i,j}} &= \frac{\left[R_{\underline{i,j}}\right]_0}{[R]_0} \\ &= \frac{r_{\underline{i,j}}}{1 + r_{\underline{i,j}}}. \end{aligned} \tag{4}$$

That is, the prevalence corresponds to the ratio $r_{\underline{i,j}}$ mapped to the interval $[0, 1]$. Under a restrictive set of assumptions (see section 4), the inferred prevalence $f_{\underline{i,j}}$ corresponds to the fraction of molecules with a connection between probe site $i$ and patch site $j$, and the ratio $r_{\underline{i,j}}$ corresponds to the true ratio of molecules, as indicated in (4) and (3). However, these assumptions, which include the assumption of equilibrium, may not be true in practice. Nonetheless, we expect the prevalence to be a monotonic function of the fraction of molecules with a connection, since it is corrected for both patch and probe affinity. It is therefore a useful quantity to characterize the population of RNA connections.

When a binding site of a probe $i$ is wholly contained within the binding site of a patch $j$, we say that the probe is *blocked*. We assume that blocked probes cannot attach if a patch has bound to their site. The integrated fluorescence values for these blocked patch-probe combinations should therefore be smaller than the background, since there is no way for the patch to increase the binding of the probe, and consequently the signal values should lie between 0 and $-1$ for blocked probes. In this case, the model becomes

$$\hat{I}_{ij}^{\text{blocked}} = \hat{B}_i \left(1 - p_j\right). \tag{5}$$

The intuitive explanation of (5) is that the probe can bind only to the molecules that are not patched, the fraction of which is $1 - p_j$.

We can then calculate the posterior $p(f_{\underline{i,j}} \mid D)$ using a similar approach to that used for to calculate the posterior on the signal. Here, however, we infer the $p_j$ from measurements on blocked probes.

#### 2.4 Aims

We aim to calculate $p(S_{ij} \mid \mathbf{D})$ and $p(f_{ij} \mid \mathbf{D})$ using *all the data*. Each data set contains more than 200,000 integrated fluorescence measurements, from which we seek to infer about 46,000 values of $S_{ij}$ and the same number of $f_{ij}$ (corresponding to 44 patches and 1047 12-base probes).

While $S_{ij}$ and $f_{ij}$ could be estimated by processing the data, perhaps by binning or averaging quantities, it would be difficult to obtain uncertainties in these quantities through such an approach. Furthermore, discarding, averaging, or binning the measurements carries the risk of introducing systematic errors associated with the choices of thresholds, averaging methods, and bins.

By using a Bayesian model and fitting for $S_{ij}$ and $f_{ij}$ using MCMC sampling, we aim to obtain estimates and uncertainties that are as true to the original data as possible, given our prior information and assumptions. Our Bayesian statistical models are formulated to use all of the data and avoid any processing, as described below.

#### 2.5 Considerations for statistical modeling

A good statistical model is needed because the posterior is informed not only by the measurements for that particular probe $i$ and patch $j$, but also by the measurements for other patches and probes. For example, our estimates of the $S_{ij}$ depend on our estimates of the background $\hat{B}_i$, which are informed not only by the background measurements $B_{ilm}$ but also by the integrated fluorescence measurements $S_{ij'k}$ for all probes $j'$. The existence of these couplings is the reason we write the posterior as $p(S_{ij} \mid \mathbf{D})$ and not $p(S_{ij} \mid I_{ijk}, B_{ilm})$. We seek a model that accounts for these couplings.

The model must also account for other effects seen in the experiments:

1. The amount of RNA can vary from well to well, affecting the overall magnitude of the measured fluorescence.

2. The uncertainty of the measurements may be different from the value (10%) reported by the Agilent software.

3. Some outliers may be present in the background measurements for any particular probe or in the measurements for any particular patch-probe combination. These outliers might arise, for example, from a spot being occluded by the gasket, or from defects or heterogeneities in the microarray to which RNA can stick.

##### 2.6 Bayesian model for signal

The model for the signal can be broken into three blocks. The first corresponds to the likelihood and associated priors:

$$\begin{aligned} B_{ilm} &\sim \text{StudentT}\left(\mu_i, \sigma B_{ilm}, \nu_B\right) \\ I_{ijk} &\sim \text{StudentT}\left(\mu_{ij}, \sigma I_{ijk}, \nu_I\right) \\ \sigma &\sim \text{Exponential}\left(\lambda_\sigma = 10\right) \\ \nu_B &\sim \text{Gamma}\left(\alpha = 2, \beta = 0.1\right) \\ \nu_I &\sim \text{Gamma}\left(\alpha = 2, \beta = 0.1\right). \end{aligned} \tag{6}$$

The second corresponds to the model for the signal:

$$\begin{aligned} \mu_i &= c_l \hat{B}_i \\ \hat{B}_i &\sim \text{Exponential}\left(\lambda_B\right) \\ \mu_{ij} &= c_j \hat{B}_i \left(1 + S_{ij}\right) \\ S_{ij} &\sim \text{Laplace}\left(\lambda_S = 1\right). \end{aligned} \tag{7}$$

And the third corresponds to a hierarchical model accounting for variations in observation due to variations in the amount of RNA dispensed in each well:

$$\begin{aligned} c_{l=1} &= 1 \\ c_{l\neq 1} &\sim \text{Normal}\left(\mu_c, \sigma_c\right) \\ c_j &\sim \text{Normal}\left(\mu_c, \sigma_c\right) \\ \mu_c &\sim \text{Normal}\left(\mu_{\mu_c} = 1, \sigma_{\mu_c} = 0.1\right) \\ \sigma_c &\sim \text{Exponential}\left(\lambda_{\sigma_c} = 10\right). \end{aligned} \tag{8}$$

A graphical description of the model for inferring the signal is shown in Figure S6. We describe each block, its parameters, and the choices of priors and likelihood in detail below.

##### 2.6.1 Likelihood

The first block, (6), defines the likelihood function. We use a Student $T$ distribution to model the measurements because this distribution is less sensitive to outliers than a normal distribution (1). We infer both the relative standard deviation $\sigma$ and the degrees of freedom $\nu_B$ (background) and $\nu_I$ (signal) of this distribution from the measurements. By doing so we avoid having to decide whether a measurement is an outlier or how many outliers are present, while ensuring that the inference remains robust to outliers. The parameters $\sigma$, $\nu_B$, and $\nu_I$ vary to account for both the dispersion and over-dispersion of all the measurements in the data set.

For the relative standard deviation $\sigma$ on the measurements, we use an exponential prior with mean of $1/\lambda_\sigma = 0.1$. The mean of the prior corresponds to the estimate (10%) of the uncertainty on each measurement returned by the Agilent feature extraction software. We use an exponential prior because this is the maximum-entropy distribution for a given mean and a non-negative support.

We use a weakly informative Gamma$(2, 0.1)$ prior on both $\nu_B$ and $\nu_I$, as recommended by the developers of the statistical modeling software Stan (2). When $\nu$ is large, the Student $T$ distribution approaches a normal distribution. When $\nu = 1$, the Student $T$ distribution becomes a Cauchy distribution, with long tails. In between these two limits, the parameter $\nu$ determines how long the tails are. If there are few outliers present, we expect the inferred $\nu$ to be large.

We use the same parameter $\sigma$ for the background and for the integrated fluorescence measurements. The underlying assumption is that all measurements, whether patch-probe or background, have the same relative uncertainty. This assumption is consistent with the fact that the Agilent software reports the same relative uncertainty for all of the measurements.

However, we have no reason to assume that the over-dispersion of the patch-probe data is the same as that of the background data, so we infer separate values of the degrees of freedom $\nu_I$ for the patch-probe data and for the background ($\nu_B$).

##### 2.6.2 Model

The next block, (7), defines how the measurements are related to the true background $\hat{B}_i$ and the true signal $S_{ij}$. The third line of (7) can also be written as

$$\begin{aligned} \mu_{ij} &= c_j \hat{I}_{ij} \\ \hat{I}_{ij} &= \hat{B}_i \left(1 + S_{ij}\right), \end{aligned} \tag{9}$$

where $\hat{I}_{ij}$ is the true intensity of a patch-probe measurement, and the second line follows from the definition of the signal, $S_{ij} = (\hat{I}_{ij} - \hat{B}_i)/\hat{B}_i$.

We include scaling factors between the background measurements and the true background (for which $c_l$ are the scaling factors) and the patch-probe measurements and patch-probe integrated fluorescence (for which $c_j$ are the scaling factors). These scaling factors account for the effects of pipetting noise, which causes the amount of RNA added to a well to vary from well to well. In writing this model we assume that the amount of RNA affects the measured fluorescence linearly. By fitting for the scaling factors, we avoid having to explicitly normalize any of the measurements. We avoid explicitly normalizing the raw data because such a procedure would make the results sensitive to the choices of normalization weights. Inferring these scalings allows us to correctly propagate the uncertainty in their estimates to the signal values.

We choose an exponential prior for the true background values with a mean $1/\lambda_B$ equal to the order of magnitude of the typical background measurement, $\lambda_B$. Again, the choice of an exponential distribution is based on the maximum-entropy principle.

The parameter $\lambda_B$ is the only input to the model. It must be chosen based on the probe size and feature extraction method. We use the following values, which are based on the typical scales that we observe in our measurements:

| probe size | feature extraction method | $\lambda_B$ |
|---|---|---|
| 12 | Agilent | $1/100$ |
| 24 | Agilent | $1/1000$ |
| 12 | our code | $10^{-7}$ |
| 24 | our code | $10^{-8}$ |

The signal $S_{ij}$ can be either positive or negative; it can be negative for blocked probes. We expect that the typical scale for the ratio, which is dimensionless, will be order 1. A Laplace prior is the maximum-entropy prior that satisfies the constraints of support along the real line and known scale (the Laplace distribution is two exponential distributions, one extending in the negative direction and one extending in the positive direction). $\lambda_S$ is the inverse scale (1/mean) of the distribution.

We note that using a Laplace prior adds the square of the L1-norm to the log posterior. In an inverse-problem approach, this would be known as L1-regularization, which is typically done when the expectation is that the reconstructed result will be sparse. Here we are not using an inverse-problem approach: our goal is not to find only the maximum of the log posterior, but to evaluate the full posterior probability. We have not made the explicit assumption that the result will be sparse; instead our choice of prior is motivated by the support required and maximum-entropy arguments. But insofar as this choice of prior is mathematically related to L1-regularization, the inference scheme will favor sparse solutions.

##### 2.6.3 Hierarchical model for scaling factors

The third block, (8), defines a hierarchical model to account for the effects of pipetting noise. The scaling factors $c_l$ account for fluctuations in the amount of unpatched RNA from background well to background well (each well corresponds to a background measurement for all probes) relative to some reference amount. We define the reference to be the amount of RNA in the first well. We do this by setting $c_1 = 1$ for the first well ($l = 1$). The choice of reference is arbitrary and should not affect any results.

Similarly, the $c_j$ scaling factors account for well-to-well variation in the amount of patched RNA added. Note that since we have already selected the reference amount (corresponding to the first well of the background measurement), all scaling factors will be relative to that reference.

We use the same parameters, $\mu_c$ and $\sigma_c$, to describe the variation in the amount of RNA in the background wells as in the patch-probe wells. The underlying assumption is that the scaling factors should all be related to pipetting noise, which is the same for background wells as it is for patch-probe wells.

We define a normal prior on the scalings with mean $\mu_c$ and standard deviation $\sigma_c$. Then we define hyperpriors on each of these parameters. The prior on $\mu_c$ is chosen to be normal, centered at 1 and with a standard deviation of 0.1, reflecting our expectation that the mean of the scalings should generally be close to 1 but can be either higher or lower, depending on the choice of reference amount. We choose an exponential prior on $\sigma_c$ with a mean of $1/\lambda_{\sigma_c} = 0.1$, reflecting our expectation that the standard deviation in the scalings should be approximately the expected value of the pipetting noise (typically about 10% variation in dispensed volume), but might be larger or smaller.

The hierarchical model allows us to determine the scalings from the data without overfitting see chapter 5 of reference 1. It also allows us to estimate the magnitude of the pipetting noise.

#### 2.7 Bayesian model for prevalence

The statistical model for the prevalence the same likelihood (6) and hierarchical scaling model (8) as those of the signal, but the second block (7) is modified to accommodate the linear model for the prevalence:

$$\begin{aligned}
\mu_i &= c_l \hat{B}_i \\
\hat{B}_i &\sim \text{Exponential}\left(\lambda_B\right) \\
\mu_{ij} &= \begin{cases} c_j \hat{B}_i \left(1 + p_j r_{\underline{i,j}}\right) & \text{for blocked probes} \\ c_j \hat{B}_i \left(1 - p_j\right) & \text{otherwise} \end{cases} \\
r_{\underline{i,j}} &= \frac{f_{\underline{i,j}}}{1 - f_{\underline{i,j}}} \\
f_{\underline{i,j}} &\sim \text{Beta}(\alpha = 2, \beta = 2) \\
p_j &\sim \text{Beta}(\alpha = 2, \beta = 2)
\end{aligned} \tag{10}$$

We use a beta distribution as a prior on both $f_{\underline{i,j}}$ and $p_j$ because it has support over the interval $[0, 1]$. We choose the parameters of the beta distribution to facilitate sampling while keeping the prior as uninformative as possible. Compared to a uniform distribution over the same interval, the beta distribution leads to more rapid tuning and sampling in our MCMC approach, which is described below. Because the ratios $r_{\underline{i,j}}$ are related to the prevalences $f_{\underline{i,j}}$, we need only set a prior on only one of the two sets of parameters. We choose $f_{\underline{i,j}}$ because it is simpler to set a prior on the interval $[0, 1]$ than on the interval $[0, \infty)$.

A graphical description of the model for inferring the prevalence is shown in Figure S7.

#### 2.8 Approach

We implement the signal and prevalence models in `pymc` version 4 and use `pymc`'s No U-Turn Sampler (NUTS) to estimate the posterior for both. For the signal model, the full posterior is

$$p\left(\left\{\hat{B}_i\right\}, \left\{S_{ij}\right\}, \sigma, \nu_B, \nu_I, \left\{c_l\right\}, \left\{c_j\right\}, \sigma_c, \mu_c, \sigma_c \mid \mathbf{D}\right). \tag{11}$$

For the prevalence model, the full posterior is

$$p\left(\left\{\hat{B}_i\right\}, \left\{f_{\underline{i,j}}\right\}, \left\{p_j\right\}, \sigma, \nu_B, \nu_I, \left\{c_l\right\}, \left\{c_j\right\}, \sigma_c, \mu_c, \sigma_c \mid \mathbf{D}\right). \tag{12}$$

We use the `jitter+adapt_diag_grad` method to initialize the sampler, and then run four MCMC chains for each dataset. For the signal model, each chain draws 1500 samples, the first 1000 of which are used to tune the sampler and the last 500 of which are retained to estimate the posterior. For the prevalence model, each chain draws 700 samples, 500 tuning samples and 200 after tuning. We choose a smaller number of draws for the prevalence model because the model is more complex. The reduction in the number of tuning samples for the prevalence model does not seem to adversely affect the sampling, in that we recover similar results for each of the chains. With these choices, it takes typically 2–7 hours to analyze each dataset with either model on a Ryzen 9 3900X processor, with one chain per thread.

To estimate and plot parameters such as $S_{ij}$, $\hat{B}_i$, $f_{\underline{i,j}}$, and $p_j$, we work with their marginal distributions. Marginalization allows us to incorporate the uncertainties on all the other parameters into our uncertainty for the parameters of interest. To marginalize, we simply do not bin or segment our samples by any parameter other than the parameter of interest. For example, each dot in

Figure 2 of the main text indicates the mean and credibility for $S_{i'j'}$ marginalized over all the other parameters, including all the other signals $S_{ij}$ where $i, j \neq i', j'$. In this example, we can view our samples as being drawn from the marginalized posterior $p\left(S_{i'j'} \mid \mathbf{D}\right)$. To calculate the mean of the marginalized posterior, we average the samples over chains and iterations. We estimate the uncertainty from the standard deviation of the marginalized posterior, again calculated over chains and iterations. Calculating these summary statistics is automated by the Python package `arviz`.

We note that we infer prevalences only from measurements on 12-nt probes. In datasets containing measurements with 24-nt probes, there is only one blocked probe per patch, since the probes are the same size as the patches. Because the patch affinities $p_j$ are inferred from data on blocked probes, the sparsity of data makes it difficult to infer the $p_j$ and hence the prevalences $\underline{f_{i,j}}$ from 24-nt probe datasets. In practice we observe that sampling is slow on these datasets, and there is wide variation from chain to chain. By contrast, datasets containing measurements on 12-nt probes have data on 13 blocked probes per patch, which leads to better sampling and inferences.

### 3 Measuring and inferring patch affinities

To validate our model, we compare the patch affinities inferred from the microarray data to affinities that we measure in bulk. The bulk experiments involve combining a fluorescently labeled patch oligonucletide with unlabeled RNA, separating the mixture with gel electrophoresis, and measuring the fluorescence across two sets of bands, the slower-running bands corresponding to bound patch, and the faster-running band corresponding to unbound patch. We assume that the fluorescence measurements are linearly proportional to the patch concentrations in each band, such that the measurement yields, up to a scaling factor, the bound and unbound patch concentrations. We measure these concentrations in two types of annealing protocols:

1. We add the patch to the RNA and do not anneal.

2. We add the patch to the RNA, then anneal the mixture.

The first protocol aligns with our protocol for the unannealed microarray datasets. In these experiments, the patch is added directly to the RNA, which has previously been annealed (heated to 90°C and cooled to below room temperature). The second protocol aligns with our protocol for the other microarray datasets. In these experiments, the patch is added to the RNA, and the mixture is annealed.

There are some differences between the patching protocol for the bulk measurements and the patching protocol for the microarray measurements. First, we fluorescently label the patch and not the RNA for the bulk measurements, while we fluorescently label the RNA and not the patch for the microarray measurements. We label the patch in the bulk measurements because we expect to see a larger difference in mobility between bound and unbound patches in the gel than we would for bound and unbound RNA. To reduce nucleobase quenching of the fluorescence signal that can occur when a fluorescently-labeled patch binds to the RNA, we add a 6-T spacer between the 3'-end of the patch sequence and the fluorescent label. Second, when we patch the RNA for the bulk measurements, we use a much higher concentration of patch and RNA (300 nM for both protocols, compared to 100 nM for microarray measurements under protocol 1 and 10 nM for microarray measurements under protocol 2). The higher concentration is needed to yield a significant intensity in the bulk measurements. Third, we test only a subset of patches in the bulk experiments because of the expense involved in procuring the labeled patches. The patches tested in the bulk experiments were arbitrarily chosen.

##### 3.1 Analysis of bulk patch affinity measurements

The results of the bulk patch binding experiments are stored in two CSV files corresponding to the two protocols above. The data are stored in narrow format, one row for each measurement. We load this data and convert to an `xarray Dataset`. These datasets contain 3–6 replicates for each patch. Plots of all the bulk measurements for both protocols are shown in Figure S20. We see that the binding affinity increases, sometimes by a large amount, when the patch and RNA are annealed together.

To compare the bulk measurements to the microarray measurements, we must correct the bulk patch affinities to the concentrations used in the microarrays. We first focus on the annealed system (protocol 2), which we assume reaches an equilibrium during the annealing process. The equilibrium reaction is

$$R + \text{patch } j \xrightleftharpoons{K_{*,\bar{j}}} R_{*,\bar{j}} \tag{13}$$

where $R$ is an unpatched RNA molecule and $R_{*,\bar{j}}$ is an RNA molecule with a patch attached to site $j$. The equilibrium constant is

$$K_{*,\bar{j}} = \frac{[R_{*,\bar{j}}]}{[R]\,[\text{patch } j]}, \tag{14}$$

where the brackets indicate concentrations. In terms of the initial concentrations $[R]_0$ and $[\text{patch } j]_0$,

$$K_{*,\bar{j}} = \frac{[R_{*,\bar{j}}]}{\left([R]_0 - [R_{*,\bar{j}}]\right)\left([\text{patch } j]_0 - [R_{*,\bar{j}}]\right)}. \tag{15}$$

Dividing both numerator and denominator by $[R]_0$ and rewriting in terms of the patch affinity $p_j = [R_{*,\bar{j}}]/[R]$, we have

$$K_{*,\bar{j}} = \frac{p_j}{[R]_0\left(1 - p_j\right)\left(\frac{[\text{patch } j]_0}{[R]_0} - p_j\right)}. \tag{16}$$

For equimolar initial concentrations of patch and RNA, the equilibrium constant reduces to

$$K_{*,\bar{j}} = \frac{p_j}{[R]_0\left(1 - p_j\right)^2}. \tag{17}$$

Because the initial concentrations are equimolar, the amount of fluorescence in the band corresponding to bound patch is proportional to $[R_{*,\bar{j}}]$, and the amount of fluorescence in the unbound patch is proportional to $[\text{patch } j]_0 - [R_{*,\bar{j}}] = [R]_0 - [R_{*,\bar{j}}]$. Therefore we can estimate the patch affinities as

$$p_j \approx \frac{I_j^{\text{bound}}}{I_j^{\text{bound}} + I_j^{\text{unbound}}} \tag{18}$$

where $I_j$ is the measured integrated fluorescence for the bulk experiment on patch $j$. However, estimating the patch affinities directly from the data through (18) does not allow us to calculate uncertainties or to determine the scaling of the affinities with concentration.

Instead, we fit for the equilibrium constant $K_{*,\bar{j}}[R]_0$ using all the data. To do this, we need a model for the observed $p_j$. We solve (17) for $p_j$ in terms of $K_{*,\bar{j}}$:

$$p_j = \frac{2 + \lambda_j - \left((2 + \lambda_j)^2 - 4\right)^{1/2}}{2}, \tag{19}$$

where $\lambda_j = \left([R]_0 K_{*,\bar{j}}\right)^{-1}$, and we have chosen the negative root because $p_j$ must go to 0 as $K_{*,\bar{j}} \to 0$.

We use pymc with its NUTS sampler to infer the $\lambda_j$ from the $p_j$ observed in the bulk experiments, using (19) as our forward model. Our full statistical model is

$$\begin{aligned} p_j^{\text{observed}} &\sim N(\mu, \sigma) \\ \mu &= \frac{2 + \lambda_j - \left((2+\lambda_j)^2 - 4\right)^{1/2}}{2} \\ \lambda_j &\sim \text{Exponential}(1) \\ \sigma &\sim \text{Exponential}(10). \end{aligned} \tag{20}$$

We choose an exponential distribution with a mean of 1 as the prior for $\lambda_j$ because $\lambda_j$ is dimensionless, so our prior expectation is that it is on the order of unity. We choose an exponential distribution for the uncertainty $\sigma$ with a scale of 10 (corresponding to a mean of 1/10) because we expect the uncertainty on each measurement to be on the order of 10%. We fit the model to all the replicate data points simultaneously, with no averaging.

##### 3.2 Correction of patch affinities

To correct the equilibrium patch affinities to a different concentration $[R]'_0$, we use

$$p'_j = \frac{2 + \lambda'_j - \left((2+\lambda'_j)^2 - 4\right)^{1/2}}{2}, \tag{21}$$

where

$$\lambda'_j = \frac{1}{K_{*,\bar{j}}[R]'_0} = \lambda_j \frac{[R]_0}{[R]'_0}. \tag{22}$$

We propagate the uncertainty from MCMC sampling of $\lambda_j$ to the estimates of the corrected affinities in order to calculate their uncertainties.

We perform the same analysis for bulk measurements made with protocol 1 (where the patch is added without annealing), though we note that the system may not reach equilibrium for these conditions. Here we correct to 100 nM, the concentration at which patches are added to the RNA in the microarray experiments using protocol 1.

The results for the corrected bulk affinities are shown in Figure S21.

##### 3.3 Comparison to affinities inferred from microarrays

There are several reasons to expect the bulk measurements to not strongly correlate with the microarray measurements. First, the bulk experiments are done at 300 nM concentration, and the corrected values assume equilibrium, which may not be an accurate approximation. Second, the 6-T spacers and fluorescent labels that are added to the patches in the bulk experiments could affect their binding affinity. Third, the microarray measurements of the patch affinity are indirect. They are inferred from a model (section 2.7) and their values are related to how efficiently a bound probe molecule binds to the patched RNA.

Nonetheless, we find reasonable correlations between the corrected bulk measurements and the inferred patch affinities. Figures S22 and S23 show bar plots and scatter plots of the affinities. For patches that are not annealed, the Pearson correlation coefficient is 0.63, and for patches that are annealed, the Pearson correlation coefficient is 0.77. The higher correlation coefficient for annealed patches might be due to the annealing protocol bringing the patch-RNA binding closer to equilibrium, thereby bringing the bulk affinities closer to the values calculated through the equilibrium model. For both unannealed and annealed patches, the correlations show that the blocked probe measurements from the microarray contain useful information about the patch affinities.

### 4 Equilibrium model for probe binding

We present an equilibrium model for probe binding that leads to the linear model equations (2) and (5). We note that this derivation is only one way to obtain these equations; more generally, the linear model is the simplest model we can write for the prevalences, and its use does not necessarily entail the assumption of equilibrium.

Our goal in this section is only to demonstrate a set of conditions under which the inferred prevalences $f_{\underline{i,j}}$ are equal to the fraction of molecules in the population that contain a connection between probe site $i$ and patch site $j$. The analysis of the data does not require the assumptions in this section.

Consider a long strand of RNA with multiple binding sites. We index the binding sites in two ways: by probe-binding location $i$ and by patch-binding location $j$. For each pair of binding sites $i, j$, we assume there are two states: unconnected and connected. If the binding sites overlap (all of the nucleotides in site $i$ are contained in site $j$), then no connection is possible.

#### 4.1 Notation

- $R_{i,j}$: RNA molecule with sites $i$ and $j$ unconnected,
- $R_{\underline{i,j}}$: RNA molecule with sites $i$ and $j$ connected,
- $R$: unpatched RNA molecule, irrespective of connections,
- $R_{\bar{\imath}}$: RNA molecule with probe attached to site $i$,
- $R_{*,\bar{\jmath}}$: RNA molecule with patch attached to site $j$,
- $R_{\bar{\imath},\bar{\jmath}}$: RNA molecule with probe attached to site $i$ and patch attached to site $j$

#### 4.2 RNA internal equilibrium

Consider a pair of sites $i, j$. We assume the connected and unconnected states of this pair are in equilibrium with equilibrium constant $K^{\mathrm{f}}_{i,j}$:

$$R_{i,j} \overset{K^{\mathrm{f}}_{i,j}}{\rightleftharpoons} R_{\underline{i,j}} \tag{23}$$

where

$$K^{\mathrm{f}}_{i,j} = \frac{[R_{\underline{i,j}}]_0}{[R_{i,j}]_0}, \tag{24}$$

where the subscript "0" indicates that the RNA is in equilibrium with itself in the absence of any patches or probes. We can write $K^{\mathrm{f}}_{i,j}$ in terms of the equilibrium fraction of connected molecules

$$f_{ij} = \frac{[R_{\underline{i,j}}]_0}{[R_{i,j}]_0 + [R_{\underline{i,j}}]_0} = \frac{[R_{\underline{i,j}}]_0}{[R]_0}, \tag{25}$$

where

$$[R]_0 = [R_{i,j}]_0 + [R_{\underline{i,j}}]_0, \tag{26}$$

since by definition any RNA molecule either has site $i$ and $j$ connected or it does not. Written in terms of the equilibrium fractions, the expression for $K^{\mathrm{f}}_{i,j}$ is

$$K^{\mathrm{f}}_{i,j} = \frac{f_{ij}}{1 - f_{ij}}. \tag{27}$$

Note that $R_{\underline{i,j}}$ and $R_{i,j}$ represent many different states of the molecule. In $R_{\underline{i,j}}$, sites $i$ and $j$ are connected, but there are also potentially connections between other pairs of sites on the molecule $i', j'$ for all $i' \neq i$ and $j' \neq j$. In $R_{i,j}$, sites $i$ and $j$ are unconnected. In addition to connections between other pairs of sites $(i', j')$ for all $i' \neq i, j' \neq j$, site $i$ might be connected to $j', j \neq j$, and site $j$ might be connected to site $i', i' \neq i$.

#### 4.3 Probe binding (background model)

In equilibrium with DNA probes, we have

$$\begin{aligned} \text{probe } i + R_{i,j} &\overset{K_{i,j}}{\rightleftharpoons} R_{\bar{i}} \\ \text{probe } i + R_{\underline{i,j}} &\overset{K_{\underline{i,j}}}{\rightleftharpoons} R_{\bar{i}} \\ R_{i,j} &\overset{K^{\mathrm{f}}_{i,j}}{\rightleftharpoons} R_{\underline{i,j}} \end{aligned} \tag{28}$$

We can merge the first two reactions into one reaction for the binding of the probe to site $i$, irrespective of whether site $i$ is connected to $j$ or not:

$$R + \text{probe } i \overset{K_i}{\rightleftharpoons} R_{\bar{i}}. \tag{29}$$

We then assume that $[R_{\bar{i}}] \ll [R]$ and $[\text{probe}_i] \ll [\text{probe}_i]_0$. This approximation linearizes the expression for $K_i$ in terms of $[R_{\bar{i}}]$:

$$K_i = \frac{[R_{\bar{i}}]}{[R]\,[\text{probe}_i]} \approx \frac{[R_{\bar{i}}]}{[R]_0[\text{probe}_i]_0}. \tag{30}$$

We discuss this assumption in detail below. We can then write a model for the background by assuming that the integrated fluorescence is linearly proportional to the concentration of RNA molecules that are bound to probe, with proportionality constant $c$:

$$\begin{aligned} B_i &= c[R_{\bar{i}}] \\ &= c[R]_0[\text{probe}_i]_0 K_i. \end{aligned} \tag{31}$$

Note that the background gives information only about $K_i$, which is the binding spectrum irrespective of connections to any other site $j$. $K_i$ depends on $K_{i,j}$ and $K_{\underline{i,j}}$ but does not determine the equilibrium fractions $f_{ij}$. This is the reason why 1D measurements are difficult to interpret in terms of structure. Even with a model for $K_{i,j}$ and $K_{\underline{i,j}}$, the equilibrium fractions $f_{ij}$ remain unknown, and there are many combinations of $f_{ij}$ that could yield the same $K_i$. The patch-probe approach is needed to obtain information about the $f_{ij}$.

#### 4.4 Patch-RNA binding

We characterize the patch-RNA binding by the final concentration of patched RNA $[R_{*,\bar{j}}]$ for each patch $j$. For what follows all we need to know is $[R_{*,\bar{j}}]$.

#### 4.5 Patch-RNA-probe binding

Now we consider a mixture of patched and unpatched RNA molecules binding to probes. Our assumptions are

1. Bound patches do not detach from the RNA.
2. Probe binding is treated as being in equilibrium.
3. Probe binding to RNA does not significantly reduce either the total probe concentration or total RNA concentration. This assumption allows us to linearize, which we do to simplify the model. Whether this assumption applies in practice is an interesting question but not one that we consider here, because our goal is only to demonstrate the conditions under which the patch-probe integrated fluorescence $\hat{I}_{ij}$ is linearly proportional to the ratio $r_{\underline{i,j}}$, as shown in (4).
4. Competition for RNA among different probes does not significantly affect the distribution of base pairs
5. The equilibrium constant for a probe binding to a patched RNA is approximately equal that for an RNA molecule with no connection between the probe site and patch site.

In a given well of the array, we have a single patch $j$, but a population of RNA molecules. That population could contain many different molecules in which different sites $i$ are connected to the site $j$ of interest. So all the probes compete for the same pool of naked and patched RNA molecules. Consider a probe $i$ and a probe $i'$ whose binding sites overlap. Let's say probe $i$ is more efficient at binding RNA. Then the amount of RNA that could bind to probe $i'$ is less than what would be available if probe $i$ were not in competition.

To keep the model tractable, we ignore effects like these that arise from competition. With this assumption, the model is uncoupled. The concentration we have been calling $[R]_0$ then becomes not necessarily the initial RNA concentration, but rather some effective concentration of RNA molecules (a mean field).

We seek a model for the total measured integrated fluorescence

$$I_{ij} = c(R_{\bar{\imath}} + R_{\bar{\imath},\bar{j}}), \tag{32}$$

where, again, we have assumed that the integrated fluorescence is linearly proportional to concentration of bound RNA with proportionality constant $c$. We have the following reactions:

$$\text{probe } i + R_{i,j} \overset{K_{i,j}}{\rightleftharpoons} R_{\bar{\imath}} \tag{33}$$

$$\text{probe } i + R_{\underline{i,j}} \overset{K_{\underline{i,j}}}{\rightleftharpoons} R_{\bar{\imath}} \tag{34}$$

$$R_{i,j} \overset{K^{\mathrm{f}}_{i,j}}{\rightleftharpoons} R_{\underline{i,j}} \tag{35}$$

$$\text{probe } i + R_{*,\bar{j}} \overset{K_{i,\bar{j}}}{\rightleftharpoons} R_{\bar{\imath},\bar{j}}, \tag{36}$$

Again, we merge the first two reactions into one reaction for the binding of the probe to site $i$, irrespective of whether site $i$ is connected to $j$ or not:

$$\mathrm{R} + \text{probe } i \xrightleftharpoons{K_i} R_{\bar{i}}. \tag{37}$$

The linearized equilibrium equation (30) for $K_i$ is now modified because the total RNA concentration is lowered by $[R_{*,\bar{j}}]$:

$$K_i = \frac{[R_{\bar{i}}]}{[R][\text{probe}_i]} \approx \frac{[R_{\bar{i}}]}{\left([R]_0 - [R_{*,\bar{j}}]\right)[\text{probe}_i]_0}. \tag{38}$$

Using assumption 5, we can write the equilibrium constant for the binding of probe to patched RNA as

$$K_{i,\bar{j}} = K_{i,j} = \frac{[R_{\bar{i},\bar{j}}]}{[R_{*,\bar{j}}][\text{probe}_i]} \approx \frac{[R_{\bar{i},\bar{j}}]}{[R_{*,\bar{j}}][\text{probe}_i]_0}. \tag{39}$$

These two equilibrium constants can be then be used to express the total measured integrated fluorescence as

$$I_{ij} = c[\text{probe}_i]_0 \left\{\left([R]_0 - [R_{*,\bar{j}}]\right) K_i + [R_{*,\bar{j}}]K_{i,j}\right\}. \tag{40}$$

To remove the dependence on the initial concentrations, we divide by the background. The background corresponds to measurements of the integrated fluorescence for unpatched RNA. Dividing (40) by (31) yields

$$\begin{aligned}\frac{I_{ij}}{B_i} &= \frac{c\left[\left([R]_0 - [R_{*,\bar{j}}]\right)[\text{probe}_i]_0 K_i + [R_{*,\bar{j}}][\text{probe}_i]_0 K_{i,j}\right]}{c[R]_0[\text{probe}_i]_0 K_i} \\ &= 1 - \frac{[R_{*,\bar{j}}]}{[R]_0} + \frac{[R_{*,\bar{j}}]}{[R]_0}\frac{K_{i,j}}{K_i} \\ &= 1 + p_j\left(\frac{K_{i,j}}{K_i} - 1\right),\end{aligned} \tag{41}$$

where $p_j = [R_{*,\bar{j}}]/[R]_0$ is the probability that a given free RNA molecule contains patch $j$.

We now need an expression for $\left(K_{i,j}/K_i - 1\right)$. To simplify, we note that

$$K_{i,j} = \frac{[R_{\bar{i}}]}{[R_{i,j}][\text{probe}_i]} \tag{42}$$

$$K_{\underline{i,j}} = \frac{[R_{\bar{i}}]}{[R_{\underline{i,j}}][\text{probe}_i]} \tag{43}$$

$$K^{\text{f}}_{i,j} = \frac{[R_{\underline{i,j}}]}{[R_{i,j}]}, \tag{44}$$

Therefore,

$$\begin{aligned}\frac{1}{K_{i,j}} + \frac{1}{K_{\underline{i,j}}} &= \frac{\left([R_{i,j}] + [R_{\underline{i,j}}]\right)[\text{probe}_i]}{[R_{\bar{i}}]} \\ &= \frac{1}{K_i},\end{aligned} \tag{45}$$

since $[R_{i,j}] + [R_{\underline{i,j}}] = [R]$ (any free RNA molecule must have sites $i$ and $j$ either connected or unconnected). Furthermore, we can express $K_{\underline{i,j}}$ in terms of $K_{i,j}$ and the folding equilibrium constant $K^{\mathrm{f}}_{i,j}$:

$$\begin{aligned} K_{\underline{i,j}} &= \frac{[R_{\bar{i}}]}{[R_{\underline{i,j}}][\mathrm{probe}_i]} \\ &= \frac{[R_{\bar{i}}]}{[R_{i,j}][\mathrm{probe}_i]} \frac{[R_{i,j}]}{[R_{\underline{i,j}}]} \\ &= \frac{K_{i,j}}{K^{\mathrm{f}}_{i,j}}. \end{aligned} \tag{46}$$

Therefore,

$$\begin{aligned} \frac{K_{i,j}}{K_i} - 1 &= K_{i,j}\left(\frac{1}{K_{i,j}} + \frac{1}{K_{\underline{i,j}}}\right) - 1 \\ &= \frac{K_{i,j}}{K_{\underline{i,j}}} \\ &= K^{\mathrm{f}}_{i,j} \end{aligned} \tag{47}$$

Note that in deriving this expression, we have not needed to use a linear approximation. We can substitute this expression into (41) to find

$$\frac{I_{ij}}{B_i} - 1 = p_j K^{\mathrm{f}}_{i,j}. \tag{48}$$

Since, according to (24) and (3), we have $K^{\mathrm{f}}_{i,j} = r_{\underline{i,j}}$, we arrive at the following expression:

$$\boxed{\frac{I_{ij}}{B_i} - 1 = p_j r_{\underline{i,j}},} \tag{49}$$

which is the model in (2).

#### 4.6 Blocked probes

Blocked probes are those that attach to a site that completely overlaps with a patch. To obtain an expression for integrated fluorescence for these probes, we assume they cannot bind to patched RNA. With this assumption, $[R_{\bar{i},\bar{j}}] = 0$, since $R_{\bar{i},\bar{j}}$ represents a molecule with both patch and probe attached. Our model for the measured intensity of (blocked probe, patch) combinations becomes

$$I^{\mathrm{blocked}}_{ij} = c[R_{\bar{i}}]. \tag{50}$$

There is only one reaction in this case, corresponding to probes binding to unpatched RNA:

$$R + \mathrm{probe}\ i \xrightleftharpoons{K_i} R_{\bar{i}}. \tag{51}$$

Since the concentration of unpatched RNA is reduced by the amount of patched RNA, we can write a linearized approximation for $K_i$ by taking $[R] \approx [R]_0 - [R_{*,\bar{j}}]$:

$$K_i = \frac{[R_{\bar{i}}]}{[R][\mathrm{probe}_i]} \approx \frac{[R_{\bar{i}}]}{\left([R]_0 - [R_{*,\bar{j}}]\right)[\mathrm{probe}_i]_0}. \tag{52}$$

Therefore

$$\begin{aligned} I_{ij}^{\text{blocked}} &= c[R_{\bar{\imath}}] \\ &= c[\text{probe}_i]_0 \left([R]_0 - [R_{*,\bar{\jmath}}]\right) K_i . \end{aligned} \tag{53}$$

Dividing by (31) yields

$$\boxed{\frac{I_{ij}^{\text{blocked}}}{B_i} - 1 = -p_j,} \tag{54}$$

which is the model for blocked probes in (5).
